# TidyTron: Reducing lab waste using validated wash-and-reuse protocols for common plasticware in Opentrons OT-2 lab robots

**DOI:** 10.1101/2023.06.09.544400

**Authors:** John A. Bryant, Cameron Longmire, Sriya Sridhar, Samuel Janousek, Mason Kellinger, R. Clay Wright

## Abstract

Every year biotechnology labs generate a combined total of ∼5.5 million tons of plastic waste. As the global bioeconomy expands, biofoundries will inevitably increase plastic consumption in-step with synthetic biology scaling. Decontamination and reuse of single-use plastics could increase sustainability and reduce recurring costs of biological research. However, throughput and variable cleaning quality make manual decontamination impractical in most instances. Automating single-use plastic cleaning with liquid handling robots makes decontamination more practical by offering higher throughput and consistent cleaning quality. However, open-source, validated protocols using low-cost lab robotics for effective decontamination of plasticware—facilitating safe reuse—have not yet been developed. Here we introduce and validate TidyTron: a library of protocols for cleaning micropipette tips and microtiter plates that are contaminated with DNA, *E. coli*, and *S. cerevisiae*. We tested a variety of cleaning solutions, contact times, and agitation methods with the aim of minimizing time and cost, while maximizing cleaning stringency and sustainability. We tested and validated these cleaning procedures by comparing fresh versus cleaned tips and plates for contamination with cells, DNA, or cleaning solutions. We assessed contamination by measuring colony forming units by plating, PCR efficiency and DNA concentration by qPCR, and event counts and debris by flow cytometry. Open source cleaning protocols are available at https://github.com/PlantSynBioLab/TidyTron and hosted on a graphical user interface at https://jbryantvt.github.io/TidyTron/.

## 1. Introduction

Biotechnology labs consume an excessive amount of single-use plastics, which inflates their spending and diminishes sustainability. Single-use plastics are popular because they provide reliably clean and sterile contact surfaces for sensitive reagents^1^. Since this single-use mindset is the norm in biotech, this research has generated shocking volumes of waste. One report estimates that 5.5 million tons of plastic waste is produced annually by the combined global biology, medical, and agricultural research institutions, which is equivalent to 83% of the total amount of plastics recycled in 2012^2^.

Plastics, and laboratory plastics in particular, are currently produced in a linear economy of manufacture and disposal, with limited recycling^3^. Standard recycling of lab materials is often impractical due to chemical and biological contamination^4,5^, so this material usually ends up in landfills and incinerators. Some organizations are beginning to make efforts to reduce plastic consumption, however most of the strategies involve restricting plastic consumption and encouraging conventional recycling of uncontaminated materials^6^. Contaminated plastics are generally overlooked by sustainability initiatives since they are hazardous to the environment, however tip recycling services do exist^7^.

The global bioeconomy is poised for rapid expansion, which will amplify plastic consumption in biotechnology labs. Over forty countries have national strategies for harnessing the economic potential of biotechnology^8^. A common objective in these strategies is to scale synthetic biology capacity to provide innovation platforms for a productive bioeconomy^9^. Institutions around the world are investing in biofoundry labs that implement automation, high-throughput equipment, process scale-up, and computer-aided design to scale synthetic biology^10^. However, the resource-intensive nature of scaled biofoundries will generate excessive plastic waste if single-use conventions are maintained. Reusing plastic resources will be a key factor in generating sustainable biofoundries that expedite bioeconomic growth without associated plastic waste.

Decontamination and re-use of contaminated plastics is a promising solution to mitigate the waste, but the time investment for implementation is unjustifiable for most labs. Plastic reuse platforms have successfully saved thousands of dollars and hundreds of kilograms worth of plastic per year when implemented in a microbiology lab^4^. Furthermore, reusing lab plastics dramatically reduces the CO_2_-equivalent footprint generated by single-use plastics^5^. Several validated cleaning strategies are available for decontaminating lab plastics^11,12^, however manual cleaning is time-consuming, since a single researcher can use hundreds of pipette tips every day. Furthermore, lab plastics have many hard-to-clean crevices, so thoroughness and quality of manual decontamination may vary. Some labs have considered hiring additional personnel, but resources are often scarce in academic labs^13^. If possible, lab personnel should spend their energy on activities with a higher impact than repetitive, manual cleaning procedures.

Low-cost liquid handling robots are widely available at reasonable price-points and can be used to automate consistently high-quality plastic decontamination. The Opentrons OT-2 is an open-source liquid handling robot with an advanced Application Programming Interface library for pipette manipulation and protocol development for a wide array of molecular biology applications. Generating a wash-and-reuse protocol library for the OT-2 would enable labs to mitigate single-use plastic waste without sacrificing time and energy of lab personnel. Many labs are already equipped with OT-2 liquid handling robots that automate more central protocols such as next generation sequencing library preparation^14^, diagnostic testing^15^, DNA assembly^16^, etc. The Opentrons OT-2 liquid handling robot equipped with a multichannel pipette is available for $11,000^17^. While specialized commercial products such as the Tipnovus by Grenova are available for automated pipette tip cleaning, this equipment is several times the price of an OT-2 and has no other automation capabilities.

Here, we validate wash-and-reuse protocols for plastics contaminated by DNA and transgenic *E. coli* and *S. cerevesiae* (yeast) to mitigate the volume of plastic consumption in biological labs. To ensure that our protocols provide consistent cleaning quality and no interference during reuse, we screened cleaned tips and microwell plates alongside fresh, unused tips and plates to check for insufficient cleaning or cleaning/decontamination reagent residue. We used bleach as our primary decontamination solution, since it has been validated as an effective chemical for breaking down DNA and decontaminating lab plastics^11^. Additionally, bleach sterilization and reuse of plastic plates for *Xenopus laevis* embryo culture did not affect development or survival, which suggest that bleach will cause minimal downstream interference during many plastic reuse scenarios^18^. However, aerosolized bleach can react with ambient compounds to form corrosive HCl^19^. To address this, we added 0.2M household baking soda to the bleach in some tests, since NaHCO_3_ mitigates corrosion from these aerosols^11^. We also include the acidic detergent Citrajet in some tests, since it is reportedly effective for breaking down DNA^20^. Finally, chemical neutralizers were tested in conjunction with primary bleach disinfectant and detergent wash solutions as an additional measure to prevent residual interference during reuse. To neutralize bleach, we rinsed plastics with a 3% household H_2_O_2_ solution in order to break down sodium hypochlorite to saltwater and O_2 (g)_ (reaction 1). Additionally, we used a 1M household baking soda solution to rinse and neutralize Citrajet acid detergent. We also tested different agitation repetitions to optimize the contact time necessary for decontamination. Decontamination protocols often call for bleach soaking^21^, so these tests were important for time minimization of high throughput cleaning procedures.

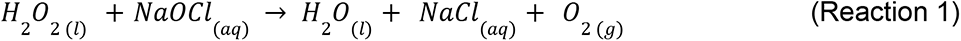

To validate our TidyTron cleaning protocols, we compared decontaminated and fresh plastics to check for residual contamination, bleach, or detergent. For *E. coli*, we compared colony forming units (cfu) for cleaned vs. fresh tips. Additionally, we calculated the initial inoculum concentration of liquid *E. coli* culture, following a microwell plate growth assay, to determine whether residues on cleaned tips kill cells and impact growth. For DNA cleaning, we performed standard PCR to check for residual DNA on cleaned tips. We also performed qPCR with cleaned vs. fresh tips to show that TidyTron protocols remove sub-nanogram contamination below the detection limit of our qPCR assay. These qPCRs also show that cleaned tips do not significantly impact downstream reuse in subsequent qPCRs. Finally, we compared cleaned microwell plates that were previously contaminated with yeast to fresh microwell plates by adding ultrapure water to resuspend residual yeast cell or debris contamination. We screened for contamination of cells and debris with flow cytometry and observed some increased debris events in cleaned plates but no significant difference in cell events between clean and fresh plates. Our results show that TidyTron cleaning protocols effectively decontaminate lab plastics while preventing interference with downstream reuse.

## 2. Methods

### 2.1 Reagents

Media and other reagents were prepared according to Green et al^22^. Lysogeny broth (LB), Miller (Fisher BioReagents) medium with kanamycin at 50 µg/mL was used for growing *E. coli*, and 0.7% (w/v) bactoagar was added for plating. Agarose gels (1% (w/v) in 1x Tris-acetate-EDTA) were stained with Biotium GelRed® Nucleic Acid Gel Stain, and bands were imaged using an iBright imaging system (Thermo Fisher). The 1 kb plus DNA ladder (NEB) was used for fragment length comparison via agarose gel electrophoresis.

### 2.2 *E. coli* Screens

For plating screens (Figure 1A, Figure 1B), *E. coli* was diluted to approximately 20 cells/µL and 10µL of liquid culture was plated. Plasmids used included pIDMv5K-J23100-tsPurple-B1006, pIDMv5K-J23100-YukonOFP-B1006, pIDMv5K-J23100-aeBlue-B1006 and pIDMv5K-J23100-eforCP-B1006 (gifted from Sebastian Cocioba).

**Figure 1:**
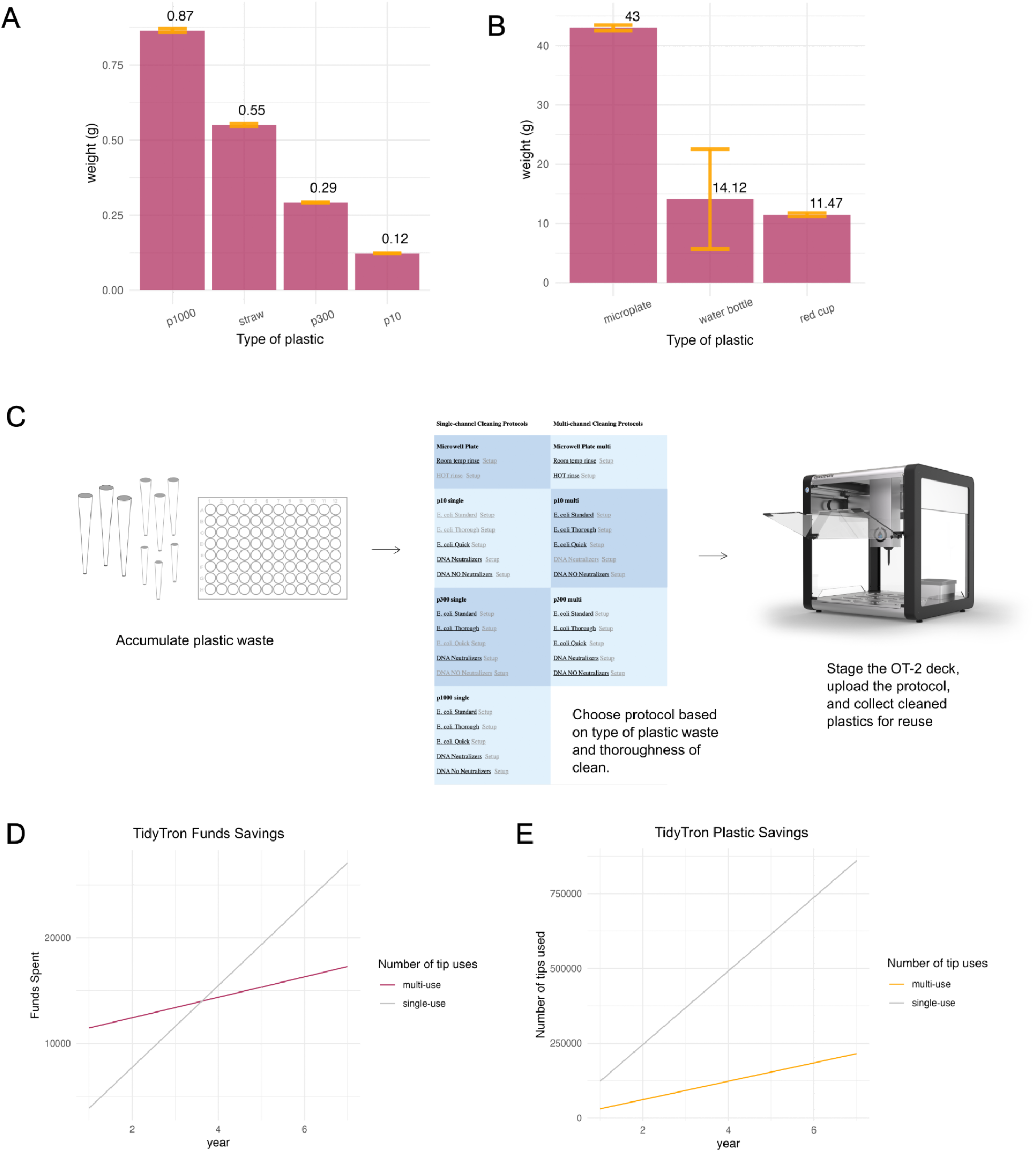
TidyTron saves plastic and funds. A) Average weight of plastic pipette tips plotted with average weight of plastic straws. B) Average weight of 96 well microplates plotted with average weight of red party cups and water bottles. Each mean was obtained from six measurements, and 6 different brands of water bottles were used, which explains the high standard deviation of this mean. C) TidyTron workflow: user accumulates plastic, downloads TidyTron protocol and setup instructions for jbryantvt.github.io/TidyTron, and stages deck according to instructions. D) Plot of money spent on pipette tips over 7 years. The red line depicts tip reuse with TidyTron, and the gray line depicts single-use of tips. The model assumes the following parameters: tip cost = $0.0315, lab personnel = 4, tips per day = 192, tip reuse = 4X, personnel work day per week = 4, personnel work weeks per year = 40, OT-2 cost = $10,500. E) Plot of pipette tip use over 7 years. The orange line depicts tip re-use with TidyTron, and the gray line depicts single use of tips.

For liquid culture growth screens, 10 µL of a dense *E. coli* overnight culture was added to 190 µL of LB with kanamycin. Cultures were grown in a BioTek Synergy HTX microplate reader at 37°C with continuous shaking for approximately 6.5 hrs, and optical density at 600 nm (OD) readings were collected every 15 minutes. A custom R script (Supplemental Data Analysis) was used to extract the time points at which the culture was in the exponential growth phase. OD data was fitted with an exponential model to obtain values for the specific growth rate, *µ_g_*, and the initial OD, *X_0_*.

### 2.3 Standard PCR

Primers were designed using j5 (19) and were purchased from Integrated DNA Technologies. j5 is free for use in academic laboratories (j5.jbei.org). Primers and template plasmids for PCRs are described in Table 1.

**Table 1:**
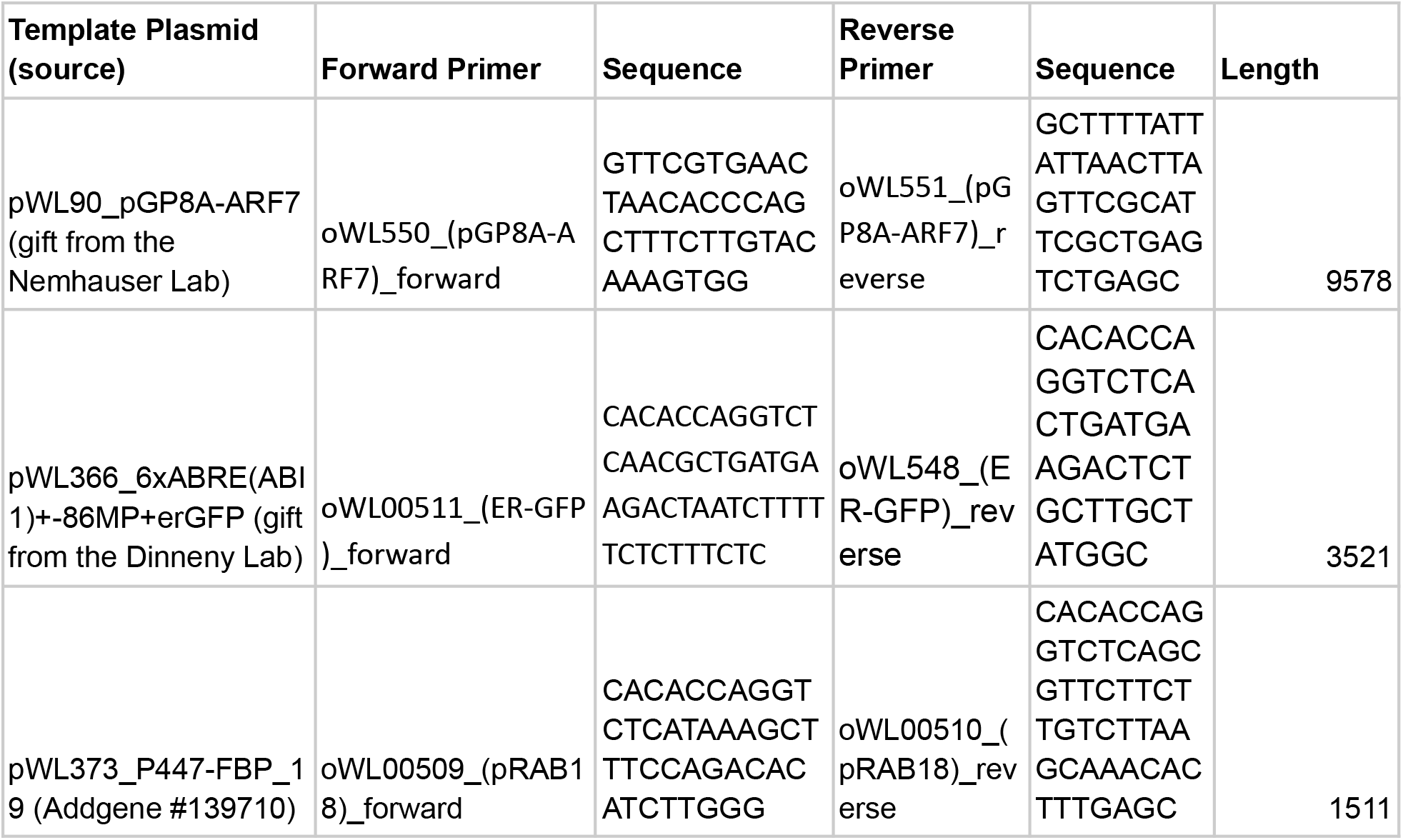
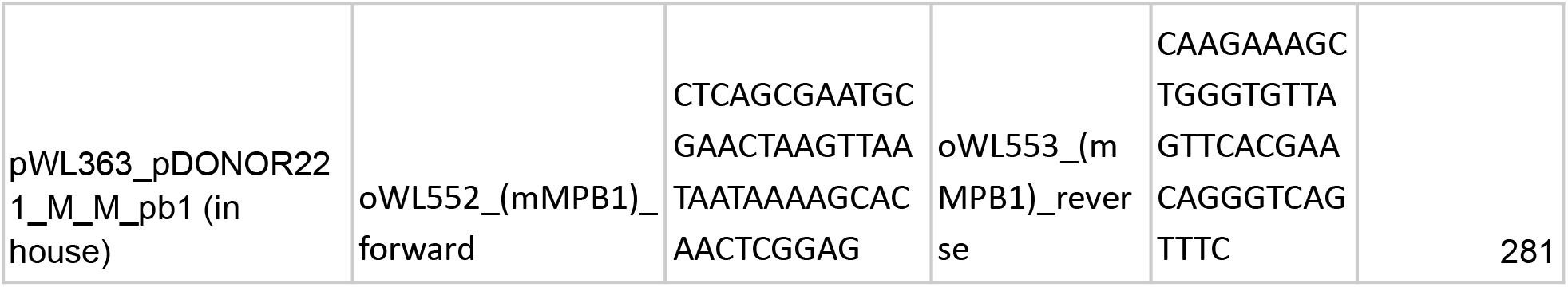
Template plasmids and primers used in PCRs in Figure 3.

### 2.4 Quantitative PCR (qPCR)

Primers for qPCR detection of pIDMv5K were designed with primer3 and were purchased from Integrated DNA Technologies (oWL558_qPCR_KanR_F aaaggtagcgttgccaatga and oWL559_qPCR_KanR_R gcctgagccagacgaaatac). The amplicon was ∼200 base pairs from the kanamycin resistance marker. pIDMv5K-J23100-aeBlue-B1006 was used as the amplicon template. A standard curve ranging from 1 uM to 0.0625 was used for back calculating starting ng in Figure 3D, and a standard curve ranging from 10^-1^ to 10^-8^ was used for Figure 3A. In Figure 3D, we performed three replicates for each measurement. The same tip was used to prepare the initial full PCR and the final post-clean full PCR. Tips used to prepare the -template and -primer PCRs were different tips. However, all tips were contaminated by preparing a 10 µL PCR mix, cleaned on the OT-2 with a TidyTron protocol, and used to prepare screen reactions. It was unnecessary to amplify every initial PCR, since there were three times as many as there were screens. So we only used the initial PCRs that were made up to contaminate tips used in the final post-clean PCR prep.

### 2.5 Flow Cytometry

Yeast cell counts were measured with an Attune NxT B/Y acoustic focusing cytometer with 488 nm blue laser excitation with 488/10 forward-scatter and side-scatter filters, and 561 nm yellow laser excitation with 620/15 nm emission filter for mScarlet measurements. Gates were specified manually with the Attune NxT software interface. Microwell plates were contaminated with 100 µL of dense yeast culture. Haploid and diploid YPH500 yeast strains expressing mScarlet driven by a pGPD constitutive reporter were used for contaminating microwell plates. Since both strains express the same mScarlet protein, the gates still defined live cells from dead cells and debris well, and experiments were aimed at simply removing yeast cells from plates, we combined data from both strains.

## 3. Results

Single-use plastics are a major source of plastic waste that are often overlooked by the scientific community. To highlight this, we measured the weight of the lab plastics cleaned by current TidyTron protocols and plotted them alongside more notorious single-use plastics such as straws, 16.9 fl oz water bottles, and red party cups (Figure 1A). We measured 6 replicates of each plastic type and calculated the average weight and standard deviation of each. We measured 6 different brands of 16.9 fl oz water bottles to account for some of the diversity in available water bottle brands, which explains the higher standard deviation. We found that pipette tips are comparable in weight to plastic straws, as 1.9 p300 pipette tips generate the same plastic waste in grams as 1 plastic straw. We also found that 1 microwell plate generates 3.05 times more plastic waste than a plastic water bottle and 3.75 times more than a red party cup (Figure 1B). Perhaps visualization and deeper consideration of this data will motivate labs to implement automated plastic cleaning with TidyTron.

To implement TidyTron, The user must first accumulate lab plastics over the course of a day with pipette tips being stored in OT-2 compatible tip boxes (Figure 1C). Next, the user selects the appropriate cleaning protocol from the TidyTron protocol library hosted at https://jbryantvt.github.io/TidyTron/ (Figure 1D). Each protocol has a corresponding setup file link directly to the right. Clicking links will trigger downloads of the desired files. Next, the user stages the OT-2 deck according to the setup file and uploads the protocol file to initiate cleaning (Figure 1E).

To further motivate single-use plastic decontamination and re-use with TidyTron, we plotted funds saved (Figure 1F) and plastic saved (Figure 1G) with TidyTron over seven years. These models assume that tip cost is $0.0315, lab is staffed with four personnel, tip consumption is 192 per day, tips are reused four times, personnel work four days/week and 40 days/year, and the initial OT-2 investment is $10,500. In this model, we found that TidyTron savings account for the full initial investment in a new OT-2 within four years of purchase (Figure 1F). We also show that in this scenario TidyTron will reduce tip consumption by millions of tips over a seven year period (Figure 1G). Since the parameters used to calculate this model vary between labs, we host an interactive version of these plots at https://jbryantvt.github.io/TidyTron/. Users can specify parameters that correspond to their lab and explore customized models.

### 3.1 *E. coli* Decontamination

Routine *E. coli* handling is a common source of unnecessary single-use plastic consumption in biotech labs, so we developed cleaning protocols for decontaminating tips of this microorganism. We chose household bleach as our cleaning solution since it is cheap and often used for deactivating live cultures prior to disposal. Household bleach typically provides between 5% and 7.5% sodium hypochlorite (NaHCO_3_), and whenever we specify sodium hypochlorite herein it is from household bleach containing 7.5% sodium hypochlorite. We also tested H_2_O_2_ as a neutralization solution for removing residual bleach from cleaned tips. The cleaning solution used for Figure 2A&B consisted of 1% sodium hypochlorite (NaHCO_3_ from household bleach) and 0.2M household baking soda. The cleaning protocol for these screens incorporated one submersion and wash with the cleaning solution for decontamination, two rinses in H_2_O_2_ for bleach neutralization, and one final rinse with ultrapure water.

**Fig 2:**
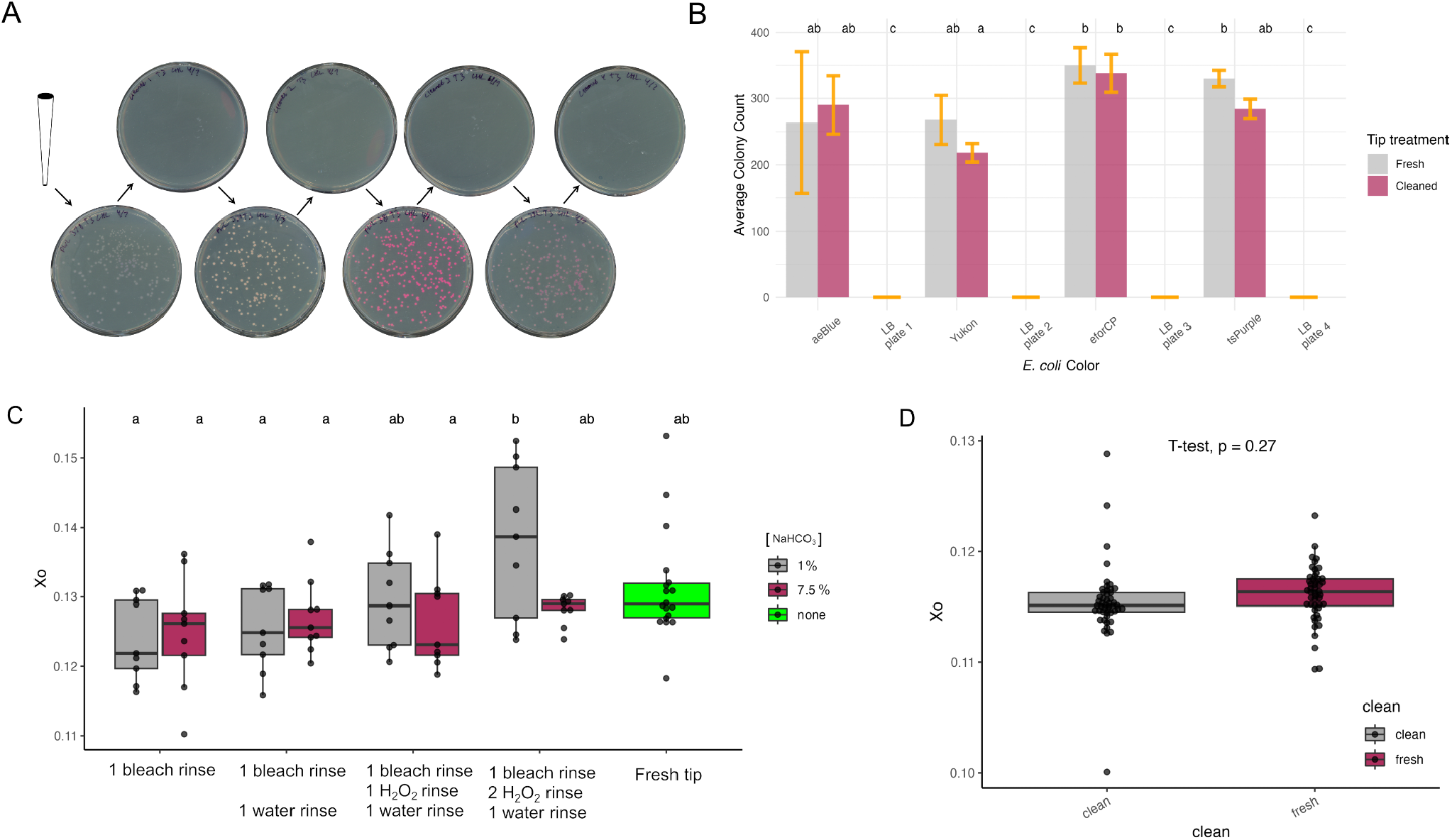
Effective *E. coli* decontamination with TidyTron: A) *E. coli* expressing four unique chromoproteins was plated with a decontaminated tip. Each arrow signifies a tip cleaning and the order in which plates were prepared. A single tip was used to prepare each plate, and there was no carry-over. B) *E. coli* and LB plating experiment was performed in triplicate along with E. coli plates prepared using fresh tips. Colony forming units (cfu) were counted and averaged. ANOVA and Tukey’s post hoc were used to detect significant differences in colony counts. C) Rinse rigor (agitation repeats and rinses) was compared by plotting *X_o_*, which was obtained from *E. coli* growth curves recorded in a shaking and incubating microplate UV/Vis spectrophotometer. *X_o_* was also recorded for fresh tips (green), and ANOVA and Tukey’s post hoc were used to detect significant differences in means. D) Mean *X_o_* was calculated for every 2 tips of a rack cleaned with 1 bleach wash + 2 H_2_O_2_ rinses + 1 water rinse was compared to mean *X_o_* obtained from 48 fresh tips with Student’s T-test. These means are indistinguishable.

In Figure 2A, we provide a qualitative demonstration of TidyTron’s ability to effectively decontaminate *E. coli* from 10 µL pipette tips. To do this, we contaminated and cleaned a single p10 pipette tip eight consecutive times. For plates 1, 3, 5, and 7, we respectively plated *E. coli* expressing blue, orange, pink, and purple chromoproteins to screen for carryover between plates. In between each *E. coli* plate (plates 2, 4, 6, and 8), we included an additional screen for carryover contamination by plating sterile LB media with the cleaned tip (Figure 2A). This screen allowed us to confirm decontamination since there was no contamination on negative control plates 2, 4, 6, or 8. Additionally, there were no colonies expressing unexpected chromoproteins on plates 1, 3, 5, or 7, which provides qualitative validation that TidyTron’s tip cleaning protocol is effective.

To quantify variation in colony counts as a result of cleaned vs. fresh tips, we replicated the experiment from Figure 2A three times to get average colony counts for each strain. To obtain colony counts for fresh tip platings, we plated identical aliquots for each *E. coli* strain with fresh tips for each replicate (Figure 2B). There was no significant difference between colony counts from plates prepared with cleaned versus fresh tips for any of the strains used. This is quantitative evidence that our TidyTron cleaning protocol does not interfere with *E. coli* viability when tips are reused.

Next, we optimized agitation repetition and rinse rigor for our TidyTron *E. coli* cleaning protocol. We applied four rinse treatments to p10 pipette tips that were decontaminated with bleach solutions of either 1% or 7.5% sodium hypochlorite (NaHCO_3_) (Figure 2C). Rinse conditions included two rinses with H_2_O_2_ followed by one water rinse, one H_2_O_2_ rinse followed by one water rinse, one water rinse, and no rinse after bleach contamination. To ensure that each rinse treatment effectively decontaminated *E. coli*, we contaminated tips, cleaned, rinsed, and plated LB on kanamycin plates, where we observed no growth (Figure S1). To compare the impact of different rinse treatments on E. coli viability, treated tips were used to inoculate a liquid culture in an incubating microreader plate, and the optical density of each well at 600 nm was measured every 15 minutes to create growth curves. A custom R script (Supplemental Data Analysis) was used to subset data to points prior to *µ_max_*in the exponential growth phase. We then fit an exponential model to each dataset to calculate an initial E. coli density (*X_0_*) and growth constant (*µ_g_*) (Equation 1). Lower *X_0_*values indicate that residual chemicals on cleaned tips killed E. coli during inoculation. We grouped *X_0_* values by rinse treatment and bleach concentration to find the optimal bleach concentration, agitation repetition, and rinse rigor for maximizing *X_0_*, or minimizing interference. We performed an analysis of variance followed by Tukey’s post hoc analysis on the *X_0_* means to detect significant differences (Figure 2C).

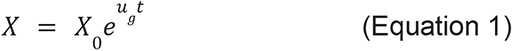

Figure 2C shows that there is very little difference in initial inoculum viability between different wash/rinse treatments. Furthermore, we observed indistinguishable *X_0_* values when tips were fresh compared to any tip cleaning treatment (Figure 2C). The only significant difference we observed is that a wash/rinse treatment consisting of 1% NaHCO_3_ + 2 H_2_O_2_ rinses + 1 water rinse leads to a significantly higher *X_0_* than when there are no H_2_O_2_ rinses or when one H_2_O_2_ rinse is paired with a 7.5% NaHCO_3_ wash. In a replicate of the experiment, we found that one 7.5% NaHCO_3_ wash + 1 H_2_O_2_ rinse + 1 water rinse kills significantly fewer cells in the inoculation than when no rinses are incorporated after a 1% NaHCO_3_ wash (Figure S2). These results provide evidence that an H_2_O_2_ rinse step reduces interference effects from cleaned tips. Interestingly, the 1% NaHCO_3_ + 2 H_2_O_2_ rinses + 1 water rinse is the only *X_0_* value in Figure 2C significantly higher than the no rinse tests. However, since all of the wash/rinse treatments fall in the same Tukey post-hoc grouping as the fresh tips, this indicates that bleach residues may not cause significant interference with *E. coli* viability during reuse, even when left unrinsed. This finding strengthens the case for TidyTron implementation if potential users are concerned about bleach interference. Follow-up experiments with more sensitive assays could further confirm this hypothesis.

To confirm that cleaning quality is consistent over an entire rack of pipette tips, we contaminated and cleaned all 96 tips in a rack. Our cleaning treatment for this screen consisted of 1-7.5% NaHCO_3_ clean + 2 H_2_O_2_ rinses + 1 water rinse. We chose this treatment because O_2_ gas formed during the H_2_O_2_ rinse sporadically occludes water from entering tips during the water rinse step. We hypothesized that of the potential cleaning solution combinations, this one is most likely to lead to variable *E. coli* viability during tip reuse. We recorded growth data for every-other cleaned tip across the rack (tip 1, 3, 5, etc.), which was 48 in total. We simultaneously recorded growth data for 48 cultures inoculated with fresh tips. We then calculated *X_0_* for each well using our R script and exponential growth model. We performed a Student’s T-test on the means of fresh versus cleaned tips and also calculated the coefficient of variation for each group of *X_0_* values. We found no evidence that a rack of cleaned tips led to a lower *X_0_* value than a rack of fresh tips (Figure 2G). Furthermore, we calculated a CV of 3.07% corresponding to cleaned tips and a CV of 2.21% corresponding to fresh tips (Supplementary Data Analysis). These comparable CV’s confirm that tip cleaning does not cause downstream *E. coli* viability to vary and that our cleaning protocol is robust for cleaning full tip racks.

### 3.2 DNA Decontamination

Most biotechnology labs handle DNA for molecular cloning, diagnostics, genetic engineering, etc. Standard autoclaving cycles are not sufficient for removing DNA^23^, so single-use plastics are typically the solution for avoiding DNA contamination. Here, we introduce and validate cleaning protocols for single-use pipette tips that quickly and effectively remove DNA below the limit of detection. Our protocols enable multiple reuses of pipette tips for standard PCR, and they reduce the risks of environmental contamination after disposal. We also provide evidence that cleaned tips do not interfere with PCR amplification efficiency, which means that tips can be reused several times prior to disposal. We used qPCR as our detection screen since it is highly sensitive for detecting DNA in sub-nanogram quantities and can provide quantitative measurements of DNA quantity and amplification efficiency.

First, we used qPCR to detect residual template DNA contamination on used tips to provide evidence that our screen is capable of detecting DNA on contaminated tips. Figure 3A shows that residual DNA on pipette tips is detectable by qPCR and that cleaning is necessary. To screen our protocol’s ability to effectively remove DNA, we used nine tips in successive rounds of cleaning and PCR prep (Figure 3B). We designated three tips for handling template DNA, three for handling primer 1, and three for handling primer 2. All 9 tips were used to prepare the initial PCR. We then cleaned the tips and used them to prepare three PCR screens: one without template to screen for template carryover (-template), one without primers to screen for primer carryover (-primers), and one with a different fragment length to screen for interference of residual cleaning residues with DNA amplification (Residuals). We then cleaned the tips and started over. Overall, we prepared eight rounds of PCRs with seven rounds of cleaning in between, and the cycle is illustrated in Figure 3B. We ran the PCR products on an agarose gel to check for appropriate band lengths and contamination (Figure 3C). The first two lanes in each gel contain a ladder and a control band to show the size expected for each round of screens. Each gel has two rows of labeling, where the first row specifies the clean cycle completed prior to that band’s amplification. The second row specifies whether that band is an initial or screen band (i.e., -template, -primer, residual). Each screen band, or lack thereof, is also marked with a green box.

**Figure 3:**
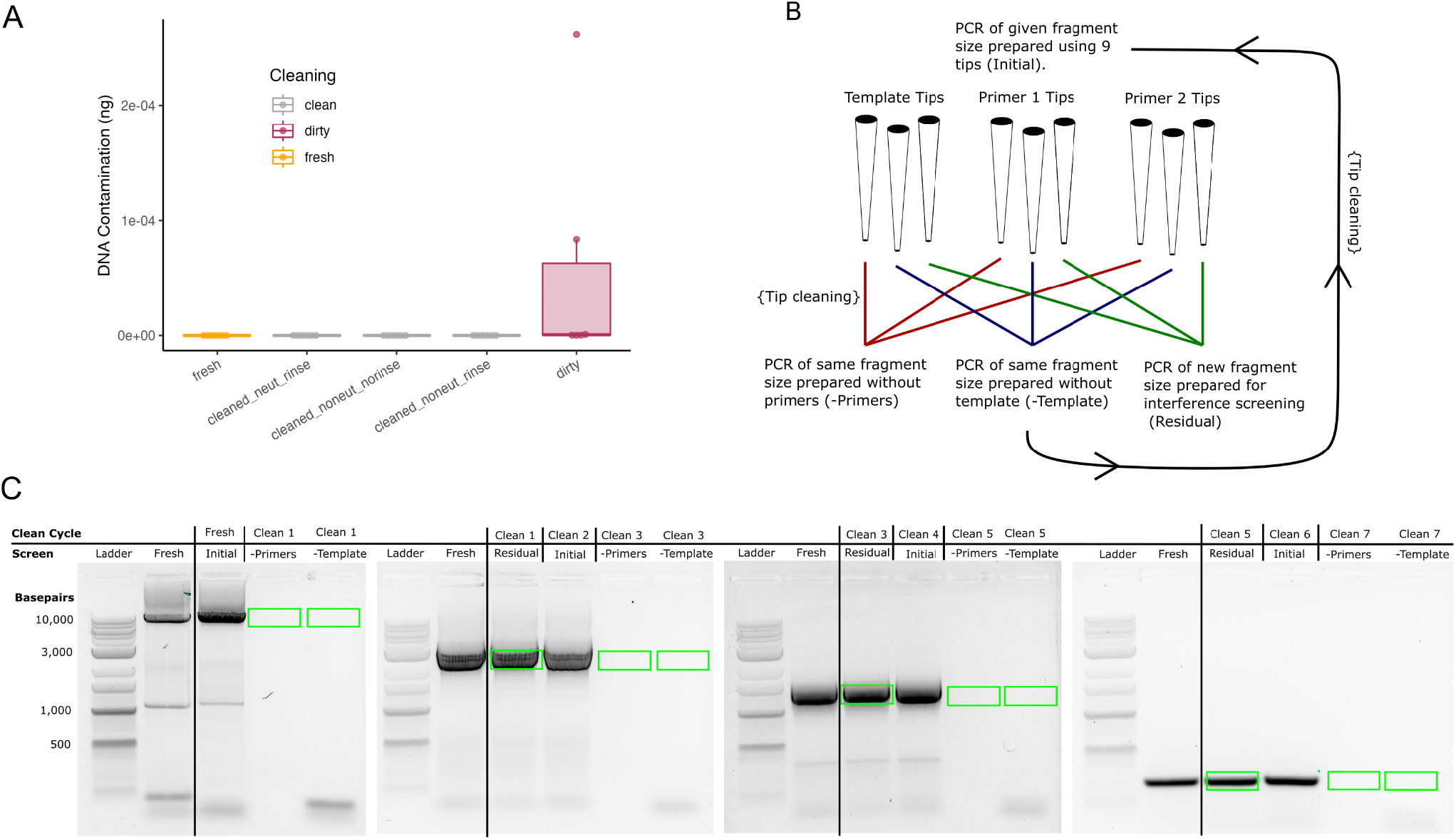
Effective DNA decontamination with TidyTron: A) qPCR shows that template DNA contamination can be detected on pipette tips. Three cleaning combinations remove contamination: clean + neutralize + rinse, clean, and clean + rinse. B) Experimental workflow for qualitative assessment of DNA decontamination: 9 pipette tips are used for successive rounds of cleaning and PCR prep, detailed in this flowchart. C) Amplicon bands from successive rounds of cleaning and PCR prep: There was no band carry-over for -template or -primer screens, and cleaned tips did not interfere with residual screens.

Initially, our cleaning protocol consisted of 3 bleach rinses, 3 H_2_O_2_ rinses, and 3 water rinses. When we screened this protocol, we detected a band for the final -template screen (figure S3). To increase the stringency of our cleaning protocol, we added 3 rinses with a 1% Citrajet acid detergent followed by 3 rinses with a 1M baking soda solution in between the H_2_O_2_ and water rinse steps. Acidic detergents are effective for breaking down DNA^20^, and the baking soda was included to neutralize any acidic residues remaining on the tip after detergent exposure. We also added a second water rinse step to the end of the protocol. As a final measure, we increased the first water rinse to four iterations and the second water rinse to six iterations. These modifications resolved our carryover issue and validated our protocol’s ability to thoroughly remove DNA (Figure 3C). Figure 3A provides quantitative evidence that our bleach + H_2_O_2_ + citrajet + baking soda + water cleaning protocol is effective for eliminating DNA contamination, since residual DNA detected from cleaned tips is indistinguishable from fresh tips.

To ensure that our cleaning procedure does not leave disruptive detergent or other residues on pipette tips, we performed an additional qPCR to compare amplification efficiency of fresh and cleaned tips (Figure 4). To measure this, we used a standard curve for quantifying initial template DNA in each PCR. If amplification efficiency is inhibited by detergent residues, the initial template concentration should appear lower. In Figure 4, the first three measurements are initial PCRs prepared with fresh tips. Measurements 4-6 lack templates (-template), measurements 7-9 lack primers (-primers), and measurements 9-12 contain all PCR components. Post-clean full PCRs were included as a screen for residue interference with PCR efficiency. Tip A was washed by the bleach + H_2_O_2_ + citrajet + baking soda + water cleaning protocol, and we found that there was no template or primer carryover. Additionally, the amplification efficiency for the post-clean full qPCR was indistinguishable from the initial qPCR prepared by fresh tips, which indicates that detergent residues do not inhibit PCR efficiency. To optimize our cleaning protocol, we cleaned Tip B with no H_2_O_2_ and baking soda neutralizers, and we cleaned Tip C with no neutralizers and no water rinse. We found that regardless of neutralization or rinse steps, bleach and citrajet effectively removed DNA from tips (Figure 4). We also found that neutralizers were not necessary for maintaining PCR efficiency between the initial and post-clean PCRs for Tip B (Figure 4). However, post-clean PCR efficiency was significantly lower than initial PCR efficiency for Tip C, perhaps since it was neither neutralized nor rinsed prior to PCR prep. However, as Tip C samples had a slightly higher mean in initial qPCRs, this difference could be attributed to random errors such as evaporation during qPCR amplification. Nevertheless, residual droplets of citrajet detergent were visible on Tip C after cleaning, suggesting possible residue contamination. Because the quantity of template DNA detected from Tip C initial and Tip C washed were still comparable, we consider this evidence that qPCR efficiency is uninhibited by TidyTron cleaning protocols, particularly for use in routine qualitative PCRs.

**Figure 4:**
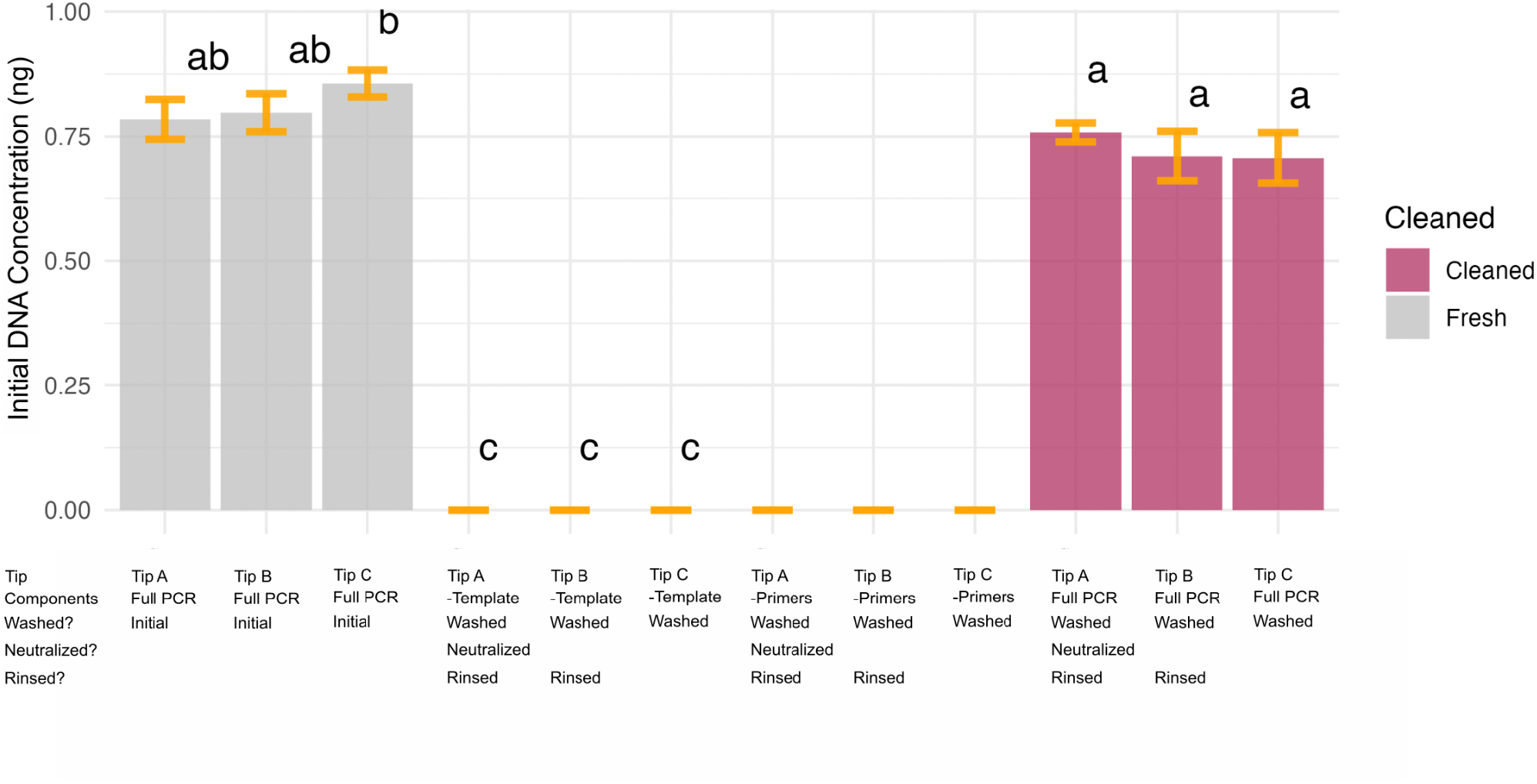
qPCR is not significantly inhibited by TidyTron cleaning protocols. qPCR was used to quantify the initial DNA concentration in a reaction prepared by tip A (cleaned, neutralized, and rinsed), tip B (cleaned and rinsed), and tip C (just cleaned). Gray bars indicate data from fresh tips. Screens without primers (-primers) and without template (-temp) indicate that no residual contamination remained on tips. Tip A, B, and C used to prepare the initial qPCR mixes were cleaned and reused to generate an identical “full qPCR” mix (maroon bars). ANOVA followed by Tukey’s post hoc analysis was performed to detect significant differences between means.

### 3.3 Yeast Decontamination

Some microorganisms, such as yeast, have cell walls that promote adherence to surfaces, which makes cleaning single-use plastics more challenging. On top of this, many microtiter plates are not autoclave-safe, which increases the importance of reliable cleaning procedures that reliably decontaminate each well. To address this need, we developed a TidyTron cleaning procedure for removing yeast from 96-well microtiter plates. Our cleaning procedures consisted of consecutive agitations with combinations of 7.5% or 1% NaHCO_3_ and 0.2M baking soda cleaner, 3% H_2_O_2_ neutralizer, and room temperature or heated ultrapure water rinse. To validate our method, flat-bottom polystyrene microtiter plates were contaminated with 100 µL of a dense liquid culture of yeast. Plates were left standing for at least two hours to allow cells to settle and adhere to the plastic. We then placed the plate directly into the OT-2, where we tested variants of cleaning protocols and solutions side-by-side (Figure 5B).

**Figure 5:**
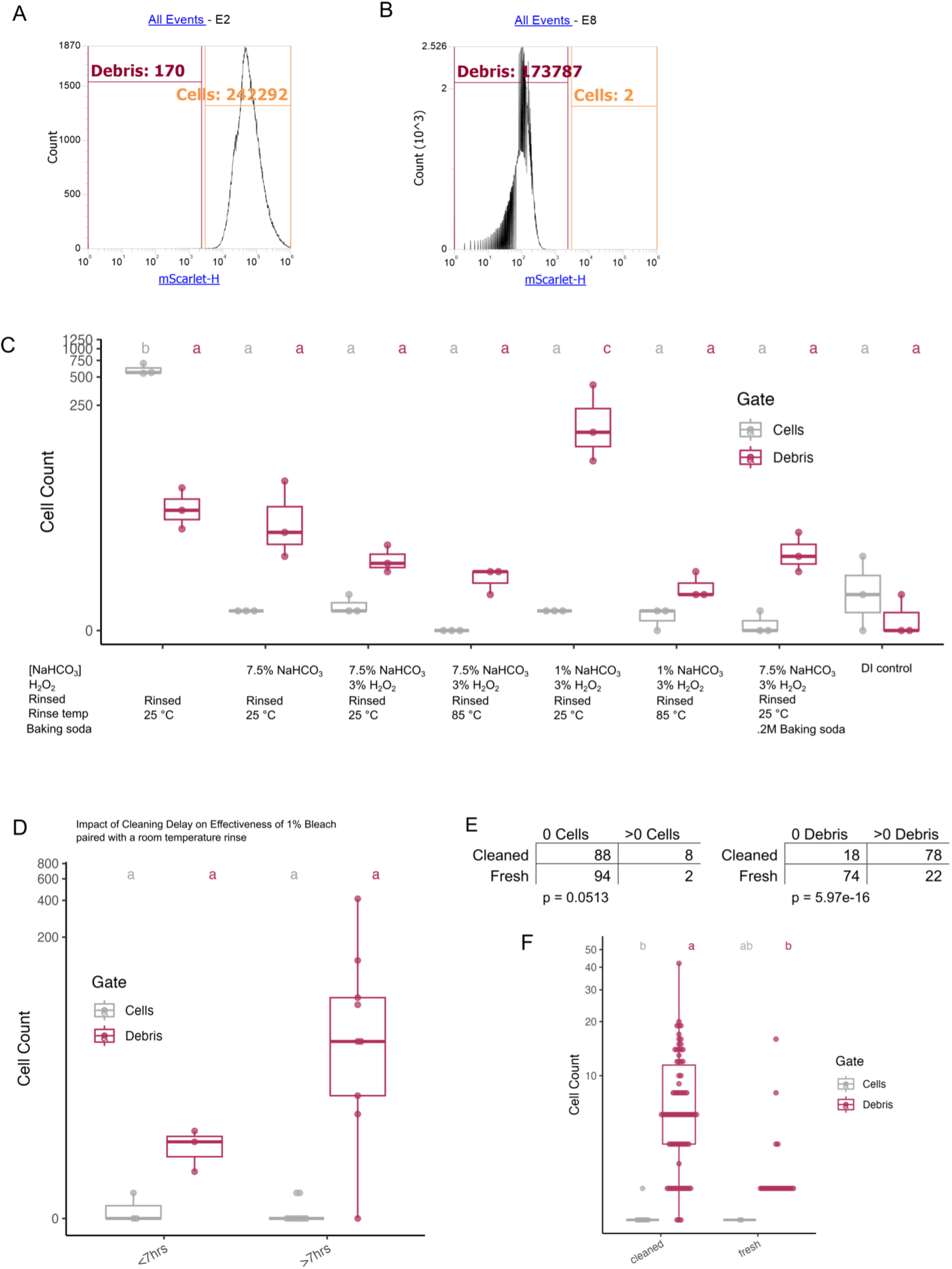
Effective yeast decontamination with TidyTron: A) Gates were applied to mScarlet histogram plots to classify events as live cells. B) A second gate was applied to classify events as debris (or dead cells). C) The three highest cell counts for each cleaning combination were selected and plotted to compare cleaning strategies. An ANOVA and Tukey’s post hoc test were used to screen for significant differences between cell count and debris means. D) Data for the 1% NaHCO_3_ clean + H_2_O_2_ rinse + 25°C rinse was subset into two groups: those set for over 7 hours (>7 hrs) and those that set for less than 7 hours (<7 hrs). ANOVA found no significant differences between means, however a clear trend in variability appeared between the two groups. E) A chi-square test was used to determine whether cleaning had an effect on presence of cells or debris. F) Nonzero cell and debris counts across an entire plate were compared for a fresh plate vs. a cleaned plate. ANOVA and Tukey’s post hoc were used to screen for differences between means.

To screen microtiter plates for cleanliness, we used flow cytometry, which is highly sensitive and capable of detecting single-cell contamination. To differentiate between live cells and debris (dead cells), we used a yeast strain expressing constitutive mScarlet red fluorescent protein to contaminate plates. Since cells express distinctly higher levels of mScarlet while living and healthy than after bleach sterilization, we designed gates based on a histogram plot of mScarlet expression. We designed the “cells” gate based on a live culture of mScarlet yeast (Figure 5A). We then added bleach to this same culture and designed our “debris” gate, which has distinctly lower mScarlet fluorescence (Figure 5B).

To screen cleaned microtiter plates with flow cytometry, we added 100 µL of ultrapure water into wells to resuspend any residual cells or debris prior to screening. We then recorded average event counts for each cleaning procedure, along with event counts for fresh, unused wells. We performed an analysis of variance on cell counts against cleaning method (or fresh wells) and gate followed by Tukey’s post-hoc analysis to screen for significant differences between fresh versus cleaned wells. Since contamination is sporadic, we selected the top three highest event counts for each cleaning procedure to highlight the worst circumstances of residual contamination (Figure 5C). We found that 1% NaHCO_3_ decontamination + H_2_O_2_ rinse + 25°C water rinse left residual contamination distinctly higher than a fresh well (Figure 5B). Nevertheless, live cell counts after this cleaning procedure are indistinguishable from fresh wells, which means that it still produces sterile conditions. The untrimmed data plot is available in Figure S4. We also show that cleaning residues do not elevate event counts compared to the fresh well (DI) control (Figure S5).

We hypothesized that the amount of time that yeast cultures have had to adhere to the plate prior to cleaning contributed to residual contamination observed after the 1% NaHCO_3_+ H_2_O_2_ rinse + 25°C water rinse protocol. To test this, we subset our data into two groups: wells that set for over 7 hours (>7 hrs) and wells that set for less than 7 hours (<7 hrs) (Figure 5D). Based on this comparison, there is distinctly higher variability in cleaning quality when yeast contacts wells for extended periods of time. We recommend avoiding the 1% NaHCO_3_ + H_2_O_2_ rinse + 25°C water rinse protocol, unless plates can be cleaned immediately after use. Aside from this caveat, our data demonstrates that TidyTron provides thorough cleaning quality for flat bottom polystyrene microtiter plates, even when they are used for handling yeast.

The final plate cleaning protocol that we suggest for microtiter decontamination consists of 7.5% NaHCO_3_ decontamination + H_2_O_2_ rinse + 25°C water rinse, since this does not require heating. We integrated tip cleaning into our final protocol so that one tip rack can be designated and re-used for plate filling and cleaning. This protocol uses the p300 multichannel pipette to reduce yeast contact time. A single channel protocol variation is available, but it takes over 35 hours to clean a full plate. Using this single channel protocol, by row six, we found that cultures evaporated and adhered preventing, residual debris from being fully removed (Figure S6). Our final microtiter plate cleaning protocol reuses tips and cleans wells in multiplex, which reduces plastic consumption and produces a thoroughly cleaned 96 well plate in approximately 4.5 hours. We recorded cell and debris counts for wells across the entire plate and compared to a fresh plate. We detected significantly higher debris counts for the cleaned plate than the fresh plate (Figure 5E), however we find this slightly higher debris presence is unavoidable when reusing tips and plates. To show that cleaning does not lead to a significant increase in cell count, we performed a chi-square test on wells where we detected 0 cell events versus wells where we detected >0 cell events. Our results indicate that cleaning is not significantly related to detecting cell events (Figure 5E). We performed a second chi-square test on wells where we detected 0 debris vs. wells where we detected >0 debris to show that cleaning results in higher debris counts (Figure 5E). This test confirms that cleaning increases the debris event count significantly. We plotted all non-zero cell and debris counts and performed ANOVA followed by Tukey’s post hoc on the means. We found that cell count for cleaned vs. fresh plates is indistinguishable. As expected from the chi-square test, debris count was significantly higher for a cleaned plate than a fresh plate.

Debris count is low enough that it will not interfere with downstream plate reuse. For integrated tip and plate cleaning, we recorded 621 cumulative events versus 70 cumulative events for the fresh plate (Figure 5F). This averages out to approximately 0.129 events/µL for the cleaned plate vs 0.015 events/µL for fresh plates. Since yeast samples typically range from 200-2,000 events/µL for cytometry analysis, the residual debris left behind by our integrated plate cleaning protocol is negligible. Moreover, debris is typically removed via gating or thresholding prior to flow cytometry analysis, so any potential interference from debris remaining after TidyTron cleaning would be removed. Taken together these results suggest that the integrated tip and plate cleaning protocol will permit reuse of these tips and plates for routine flow cytometry of yeast cultures.

## 4. Discussion

Single-use plastics are ingrained in modern biotechnology procedures because of the convenience and reliable sterility that they offer. However, the level of waste generated by single use plastics is shockingly substantial. By using two racks of pipette tips in a single experiment, a researcher is likely to generate a higher volume of plastic waste than they would from a year of plastic straw use at restaurants (based on conversion factors from Figure 1A). A single graduate student is likely to generate a higher volume of plastic waste over the course of their PhD than they would from using straws over the course of four lifetimes (Figure 1A, 1B). Despite these disturbing predictions based on our conversion factors from Figure 1A, the ends have justified the unsustainable means of science; until now.

Here we introduce and validate TidyTron to foster sustainability and plastic reuse, while maintaining the convenience and reliability provided by single-use plastics. Our automated wash-and-reuse protocols can mitigate plastic waste and recurring consumable costs for biotechnology labs. We provide data to validate that our protocols effectively remove *E. coli*, DNA, and yeast from common lab plastics. Notably, we did not integrate autoclaving into any of this work. The fact that we were able to achieve such high standards for decontamination prompts an important question: are autoclaves necessary for sterilizing lab plastics prior to use? Industrial autoclaves are low throughput and consume massive amounts of energy to attain high temperature and pressure^24^. Furthermore, standard autoclaving cycles do not eliminate many forms of contamination^23^. Instead, extra-long two hour cycles are necessary to achieve reliable decontamination^25^. TidyTron offers reliable, high throughput decontamination that is far more sustainable and convenient than standard or extended autoclaving. Additionally, TidyTron protocols are compatible with materials such as polystyrene that are not autoclavable.

Long-term implementation of TidyTron is crucial for reaching impactful levels of plastic conservation, so users should consider mitigating long-term corrosion of mechanical pipette due to consistent exposure to bleach aerosols. Cleaning surfaces with bleach is known to elevate levels of aerosolized OH^-^ and Cl^-^ radicals, which can react with indoor volatile compounds to yield corrosive HCl^19^. While the byproducts of this ambient chemistry won’t be immediately evident, over time we predict that routine bleach pipetting could lead to mechanical malfunctions. Adding baking soda as an additive to bleach cleaning solutions is a cheap, precautionary measure for TidyTron users who want to mitigate OT-2 corrosion risk. Our data confirms that adding baking soda to bleach solutions does not inhibit decontamination. The addition of baking soda has also been shown to have no adverse effect on bleach activity^11^. It should be noted that adding 0.2 M baking soda creates a cloudy, precipitous suspension in 7.5% NaHCO_3_, however this did not appear to inhibit decontamination activity, but may have increased residual debris in flow cytometry.

Currently, the TidyTron library is limited to cleaning protocols for pipette tips and microtiter plates, since these single-use plastics generate high volumes of waste and are difficult to clean. In the future, we plan to expand our protocol library to other lab plastics such as deep well culture plates, 5 mL culture tubes, and PCR tubes. We also aim to develop cleaning protocols for standard 15×100mm plastic culture plates or alternatives, which have been referred to as non-reusable lab plastic^13^. Establishing labs that reuse plastic resources will be a key factor in generating sustainable biofoundries capable of meeting global food, feed, and pharmaceutical needs. TidyTron represents a critical step towards altering the single-use mindset that prevails in modern biotechnology.

## Supporting information

Supplemental Data Analysis

## Supplemental Figures

**S1:**
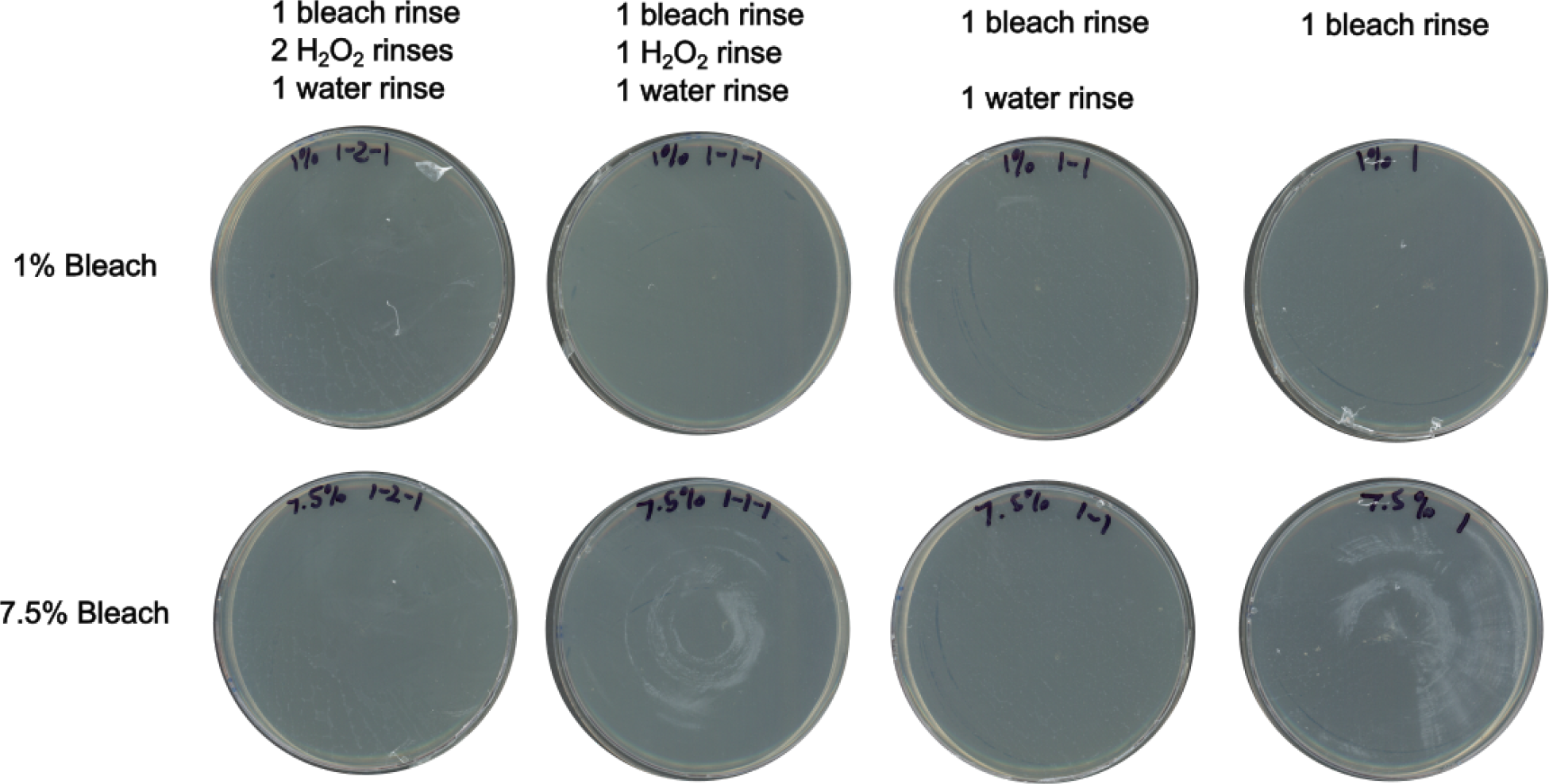
Effective decontamination for optimized tip cleaning. Here we show that successively less thorough cleaning of tips effectively eliminates E. coli decontamination. Each plate corresponds to a growth curve category in Figure 2C.

**S2:**
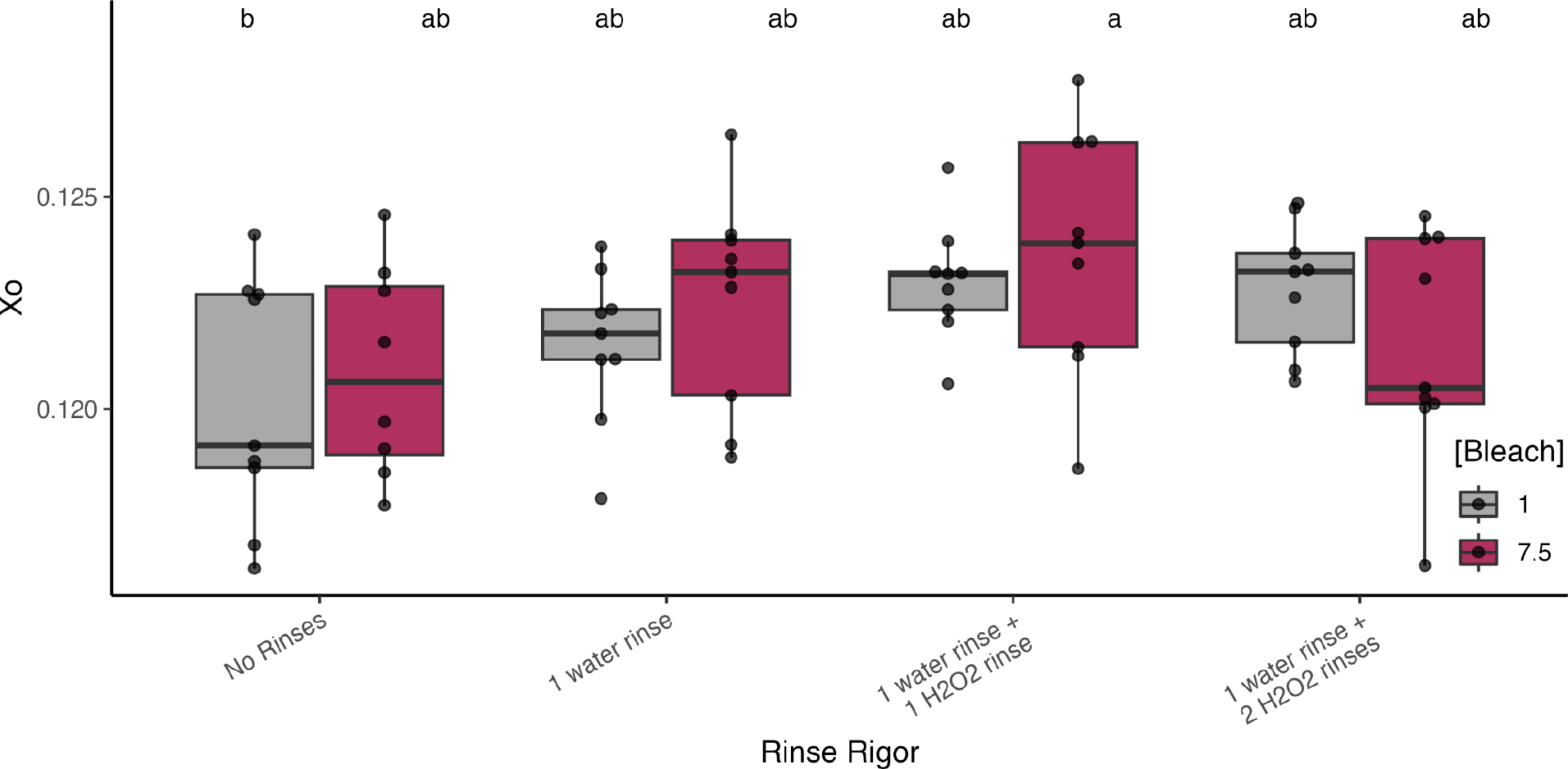
Rinse rigor (Agitation repeats and rinses) was compared by plotting *X_o_*, which was obtained from E. coli growth curves recorded in a shaking and incubating microplate UV/Vis spectrophotometer (trtmnt p= 0.0087, bleach p=0.6738, trtmnt:bleach p= 0.4003). This figure is a replicate experiment of Figure 2C and shows the same trend as the main text, which fortifies our conclusions.

**S3:**
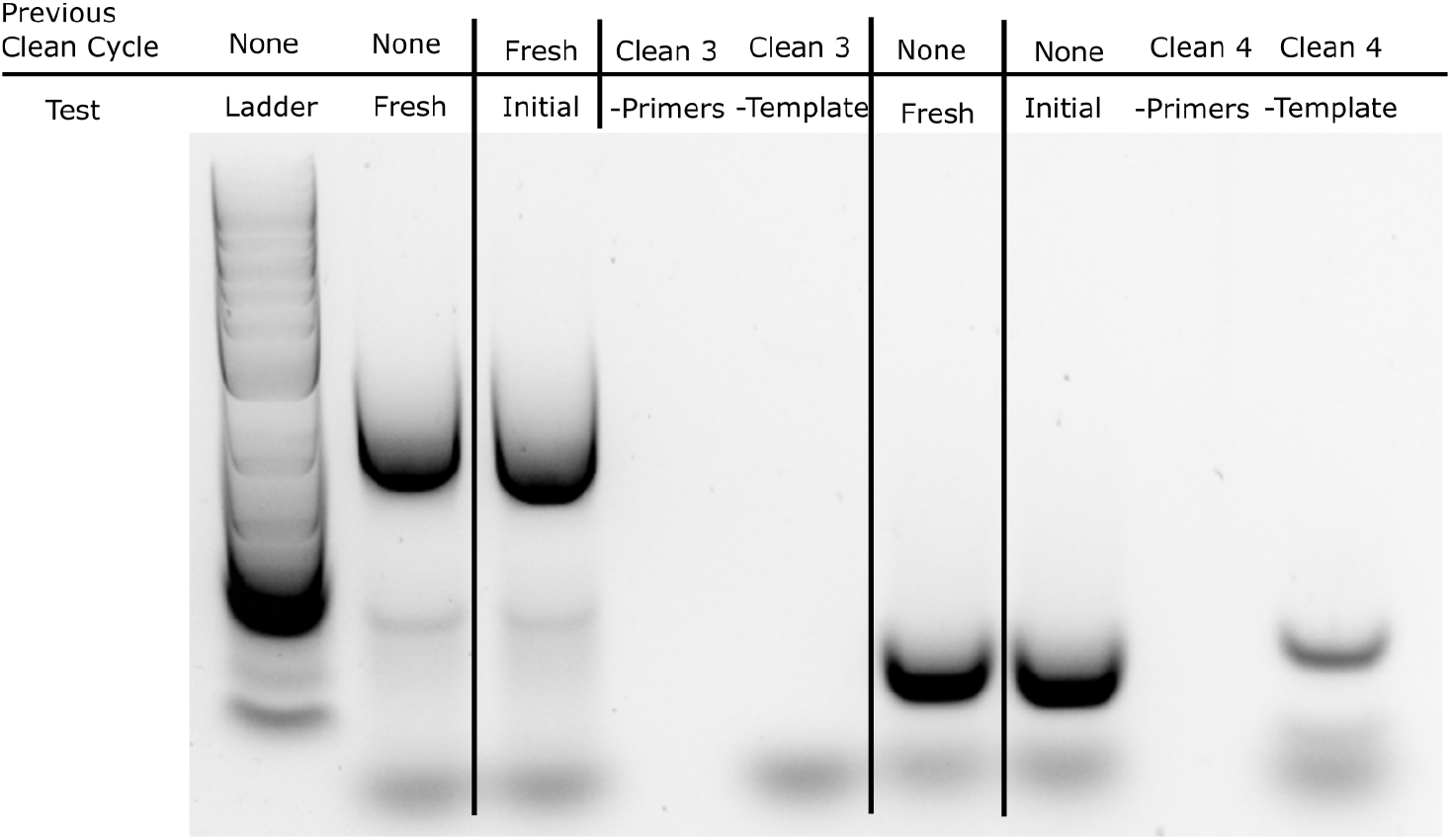
Template carryover failure. Amplicon bands are shown from successive rounds of cleaning and PCR prep: There was carryover after cleaning for the -template screen. This failure incentivized us to add additional Citrajet detergent, baking soda neutralization, and water rinse steps.

**S4:**
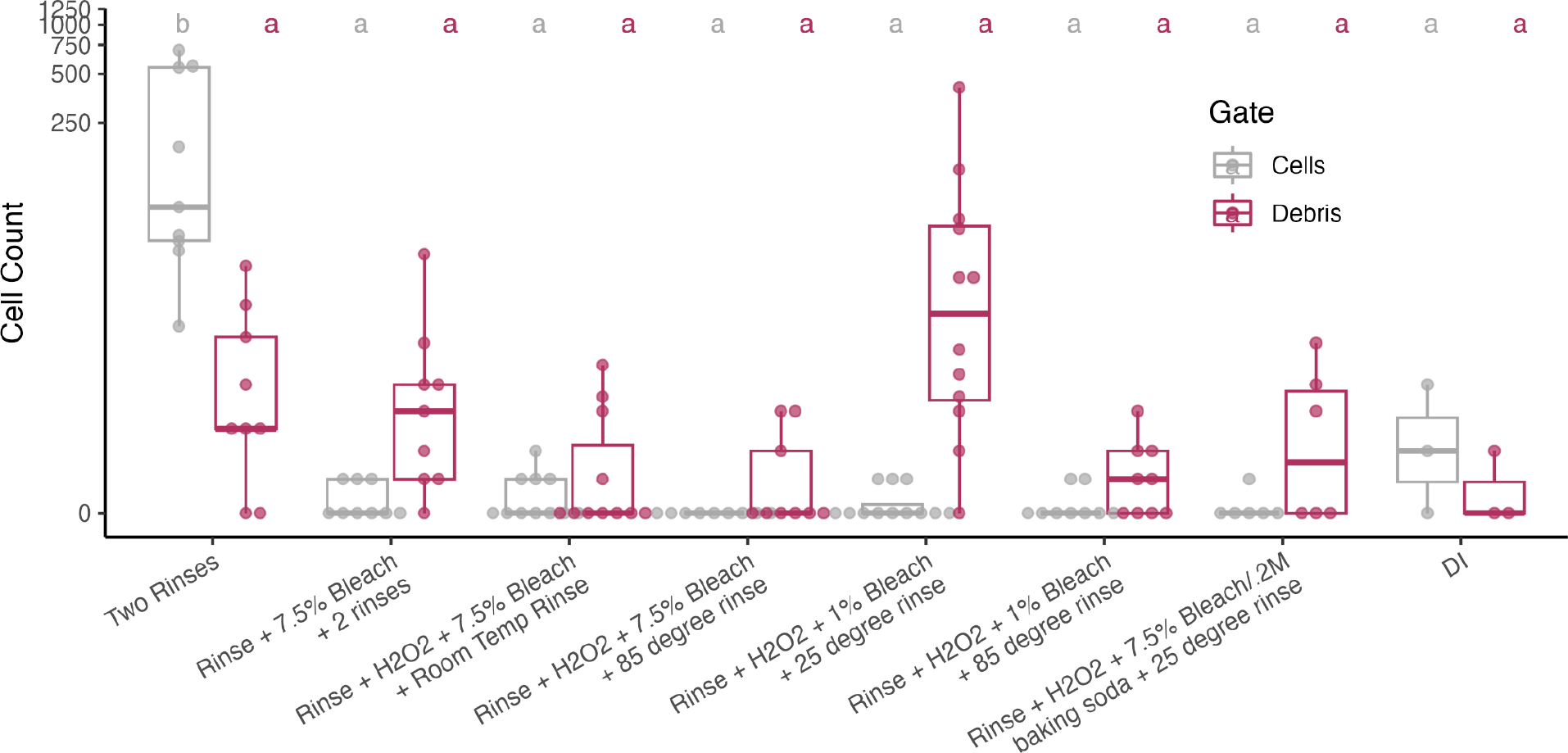
Complete dataset for yeast plate cleaning procedure comparison. All data points for each cleaning combination are plotted to compare cleaning strategies. An ANOVA (clean_method p=1.74e-05, Gate p= 0.169, and clean_method:Gate p =5.04e-06) and Tukey’s post hoc test were applied to screen for significant differences between cell count and debris means. Here, 1% bleach decontamination + H_2_O_2_ neutralization + water rinse is not significantly different from other cleaning procedures, even though a distinct trend is visible. We found that contamination appeared sporadically, so the top 3 event counts from this figure are displayed in Figure 5C.

**S5:**
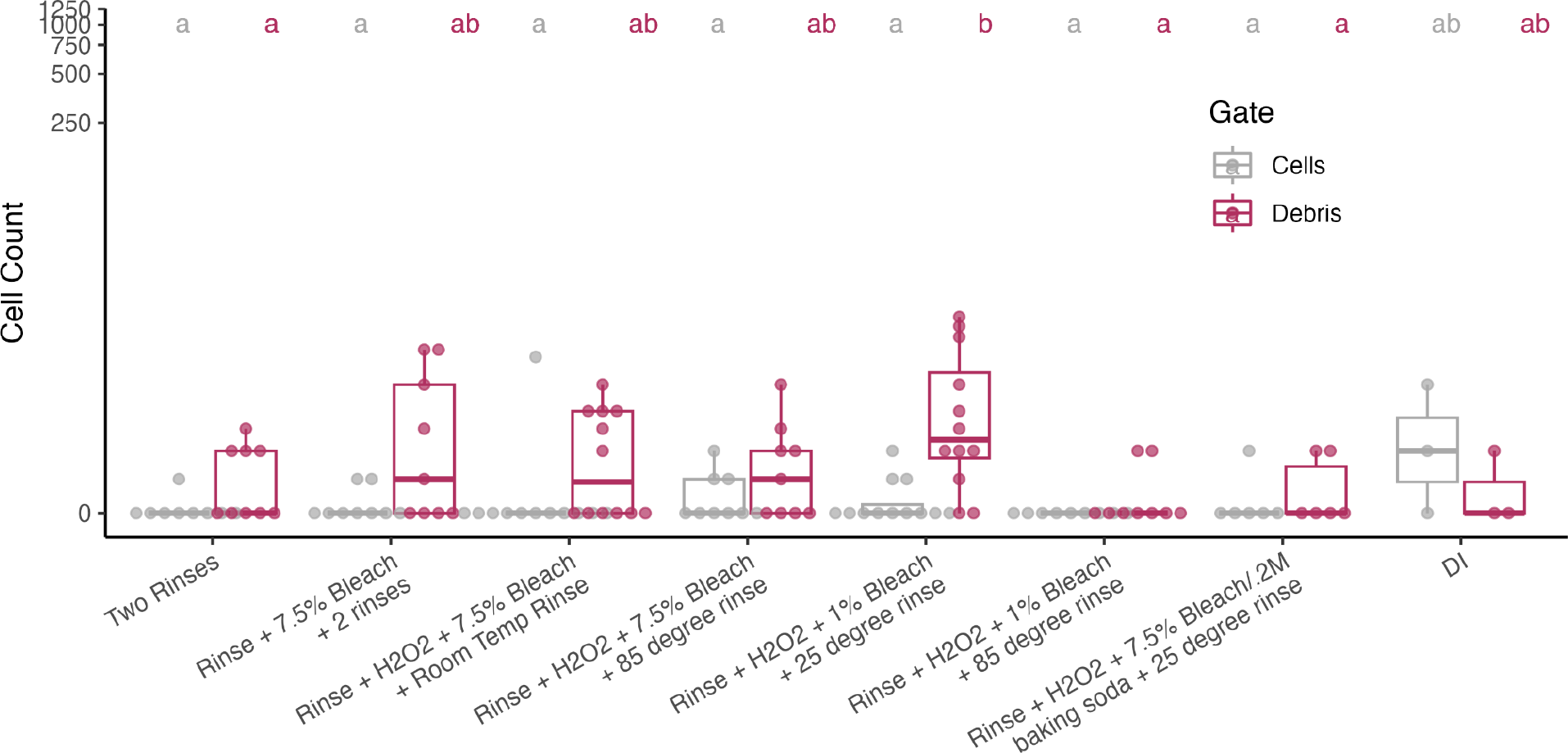
Bleach residue from plate cleaning. To check for bleach residue, we tested each cleaning procedure by cleaning wells with 100 µL of ultrapure water, instead of dense yeast culture. An ANOVA (clean_method p=0.0288, Gate p= 6.18e-05, and clean_method:Gate p =0.0149) and Tukey’s post hoc test were applied to screen for significant differences between means. We found that no cleaning procedure led to event counts significantly different from an uncleaned control (DI).

**S6:**
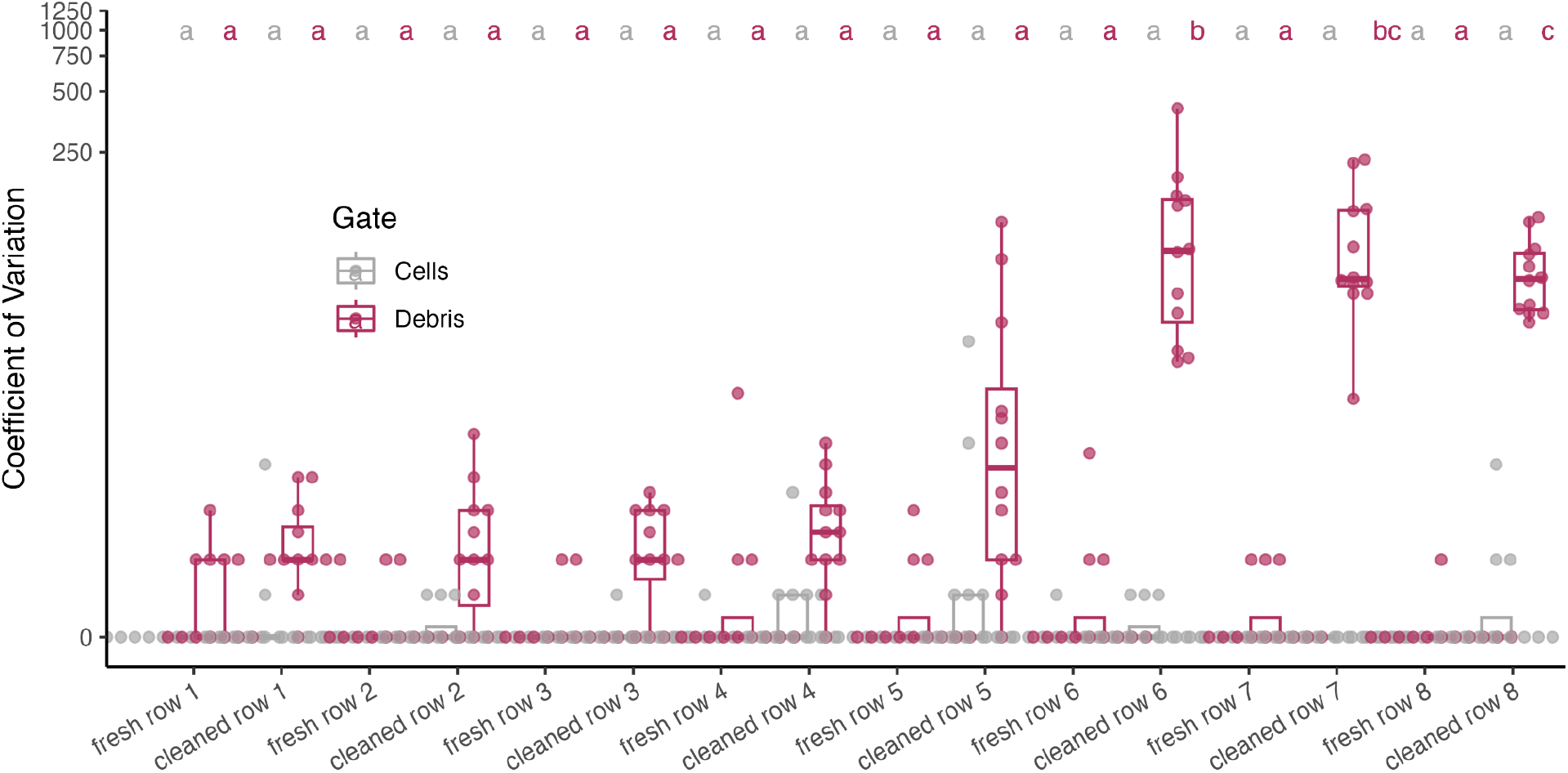
Event data for full 96 well microtiter plate cleaned with a single channel pipette and integrated tip cleaning. After cleaning an entire plate with integrated tip cleaning between each well, we compare event means of each row to event means of a fresh 96 well plate. While single channel cleaning is effective for the first half of the plate, we found that rows 6, 7, 8, and 9 had significantly higher event counts than a fresh place. (grp p = < 2e-16, Gate p = 1.51e-13, grp:Gate p = < 2e-16). This data motivates implementation of the higher throughput multichannel plate cleaning protocol offered in the TidyTron library for cleaning full microtiter plates.

## Acknowledgements

We thank Sebastian Cocioba for providing the plasmids used in this work. We also thank Patarasuda Chaisupa for designing the TidyTron Logo.

