## Supplemental Data Analysis for "TidyTron: Reducing lab waste using validated wash-and-reuse protocols for common plasticware in Opentrons OT-2 lab robots"

John Bryant

2023-05-18

```
library(multcompView)
library(ggplot2)
library(ggbeeswarm)
library(forcats)
library(dplyr)
```

```
##
## Attaching package: 'dplyr'

## The following objects are masked from 'package:stats':
##
##     filter, lag

## The following objects are masked from 'package:base':
##
##     intersect, setdiff, setequal, union
```

```
library(stringr)
library(rstatix)
```

```
##
## Attaching package: 'rstatix'

## The following object is masked from 'package:stats':
##
##     filter
```

```
library(ggpubr)
```

```
load("my_work_space.RData")
```

#Figure 1A

```
data<- read.csv("rawdata/plastics_fig1.csv")

means <- aggregate(.~plasticware,data=data,FUN = mean )
colnames(means) <- c("plasticware", "mean")
sd <- aggregate(.~plasticware,data=data,FUN="sd")
colnames(sd) <- c("plasticware", "sd")
means["sd"] <- sd["sd"]
#means <- merge(means, LABELS, by = "Sample",
 #                  all.x = TRUE)

light <- subset(means, plasticware == "straw" | plasticware == "p10"|
                  plasticware == "p300"| plasticware == "p1000")
```

```
ggplot(data = light, aes(x = fct_relevel(plasticware,"p1000","straw","p300",
                                          "p10"), y = mean,fill="maroon")) +
  geom_bar(stat = "identity", position = position_dodge(), alpha = 0.75) +
  geom_errorbar( aes(x=plasticware, ymin=mean-sd, ymax=mean+sd), width=0.4,
                colour="orange", alpha=0.9, size=1.3,
                position = position_dodge(.9)) +  labs(y = "weight (g)",
                                                         x = "Type of plastic")+
  scale_fill_manual(values=c("maroon","gray","gray","gray"))+ theme_minimal()+
  theme(axis.text.x = element_text(angle = 20, vjust = 0.5, hjust=.2),
        legend.position = "none")+geom_text(aes(label=round(mean,2)),
                                             nudge_x = .11,nudge_y = .05)
```

```
## Warning: Using `size` aesthetic for lines was deprecated in ggplot2 3.4.0.
## i Please use `linewidth` instead.
## This warning is displayed once every 8 hours.
## Call `lifecycle::last_lifecycle_warnings()` to see where this warning was
## generated.
```

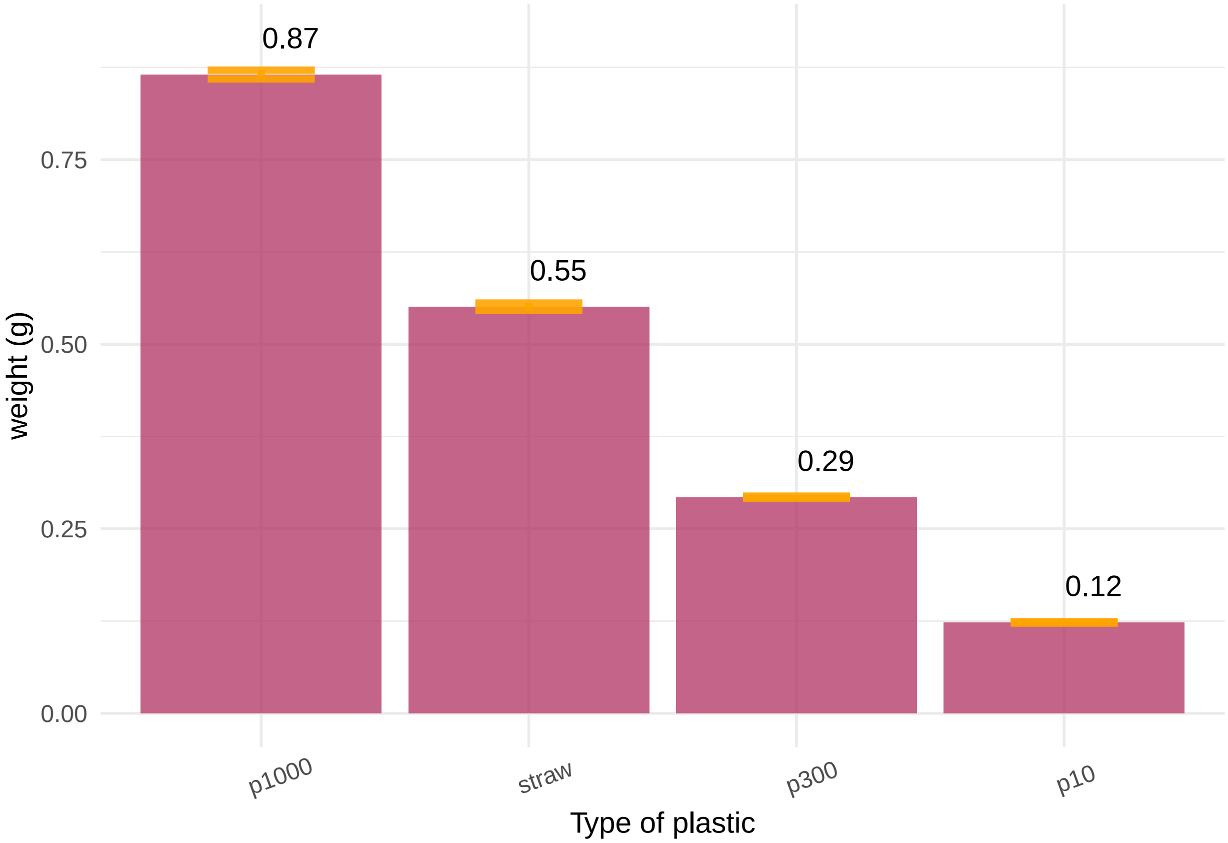


#Figure 1B

```
heavy <- subset(means, plasticware == "microplate" | plasticware == "red cup"|
                  plasticware == "water bottle")

ggplot(data = heavy, aes(x = fct_relevel(plasticware,"microplate","water bottle"
                                          ,"red cup"), y = mean,fill="maroon")) +
  geom_bar(stat = "identity", position = position_dodge(), alpha = 0.75) +
  geom_errorbar( aes(x=plasticware, ymin=mean-sd, ymax=mean+sd), width=0.4,
                colour="orange", alpha=0.9, size=1.3,
```

```
                  position = position_dodge(.9)) +  labs(y = "weight (g)",
                                                       x = "Type of plastic")+
  scale_fill_manual(values=c("maroon","gray","gray","gray"))+
  theme_minimal()+
  theme(axis.text.x = element_text(angle = 20, vjust = 0.5, hjust=.2),
        legend.position = "none")+geom_text(aes(label=round(mean,2)),
                                                nudge_x = .2,nudge_y = 2)
```

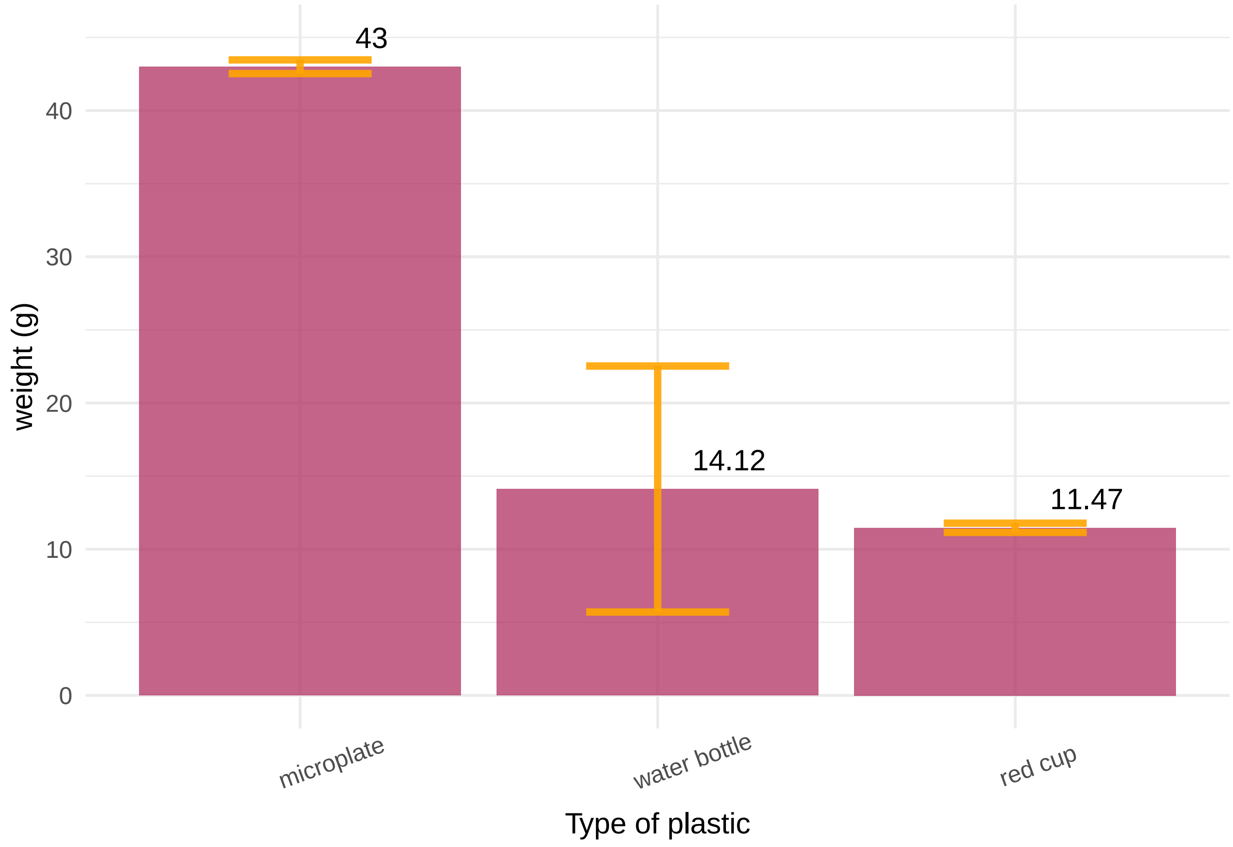


#Figure 1D

```
costsavings<- read.csv("rawdata/costsavingsplot.csv")

ggplot(costsavings, aes(x = year,y=dollars,fill=use)) +
  geom_line(aes(color = use)) +
  scale_color_manual(values = c("maroon", "gray"))+  labs(y = "Funds Spent",
                                                          x = "year")+
  theme_minimal()+labs(color='Number of tip uses')
```

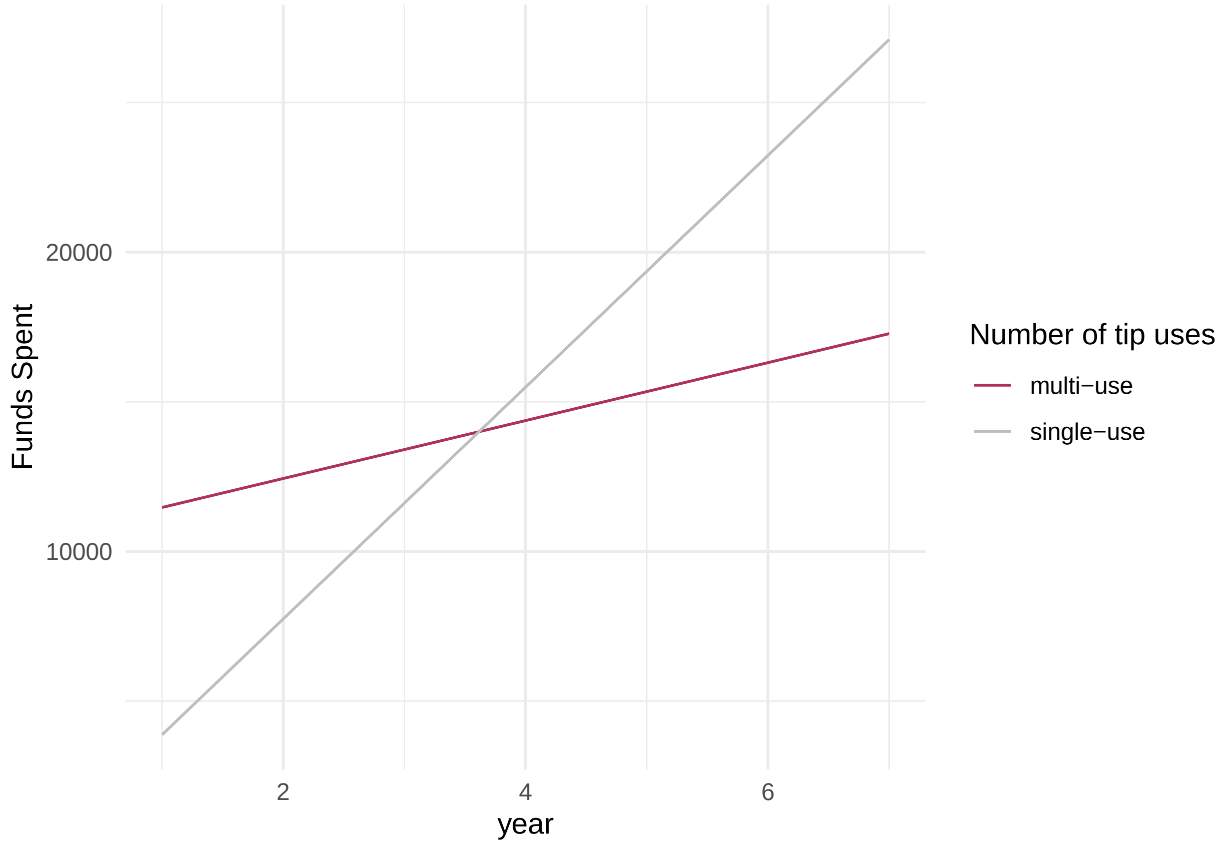


```
#ggsave("~/Google Drive/Shared drives/PlantSynBioLab/Johns Folder/TidyTron/fig1Ccash.png", width = 6, h
```

#Figure 1E

```
tipsavings<- read.csv("rawdata/tipsavingsplot.csv")
ggplot(tipsavings, aes(x = year,y=tips,fill=use)) +
  geom_line(aes(color = use)) +
  scale_color_manual(values = c("orange", "gray"))+  labs(y = "Number of tips used", x = "year")+ theme_
```

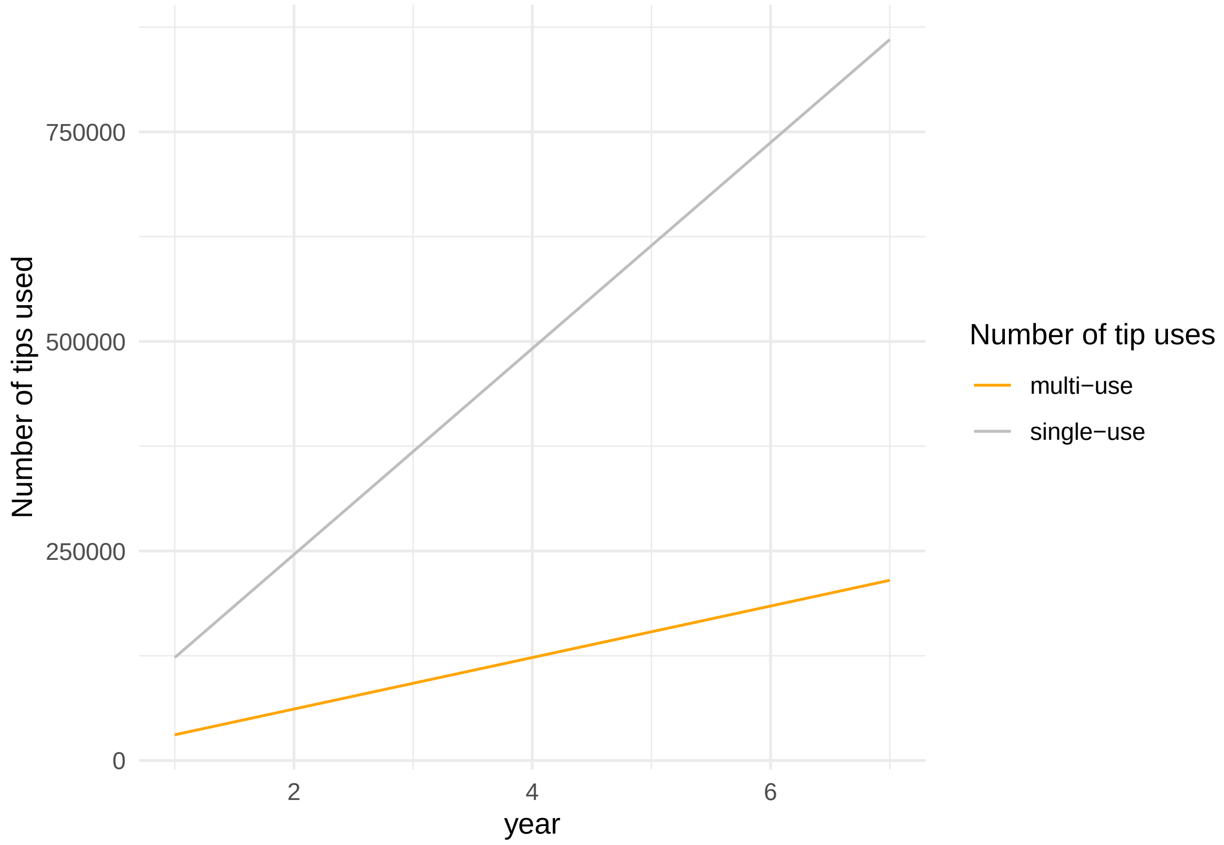


```
#ggsave("~/Google Drive/Shared drives/PlantSynBioLab/Johns Folder/TidyTron/fig1C_tipnum.png", width = 6
```

#Figure 2B

```
data <- read.csv("rawdata/20230412_counts.csv")
data$color <- str_replace_all(data$color,'-','_')
```

Run ANOVA and Tukey's

```
model=lm( data$count ~ data$color )
ANOVA=aov(model)
summary(ANOVA)
```

```
##             Df Sum Sq Mean Sq F value   Pr(>F)    
## data$color  11 727044   66095   47.02 1.64e-13 ***
## Residuals   24  33737    1406                     
## ---
## Signif. codes:  0 '***' 0.001 '**' 0.01 '*' 0.05 '.' 0.1 ' ' 1
```

```
TUKEY <- TukeyHSD(x=ANOVA, 'data$color', conf.level=0.95)
cld <- multcompLetters4(ANOVA, TUKEY)
plot(TUKEY , las=1 , col="brown")
```

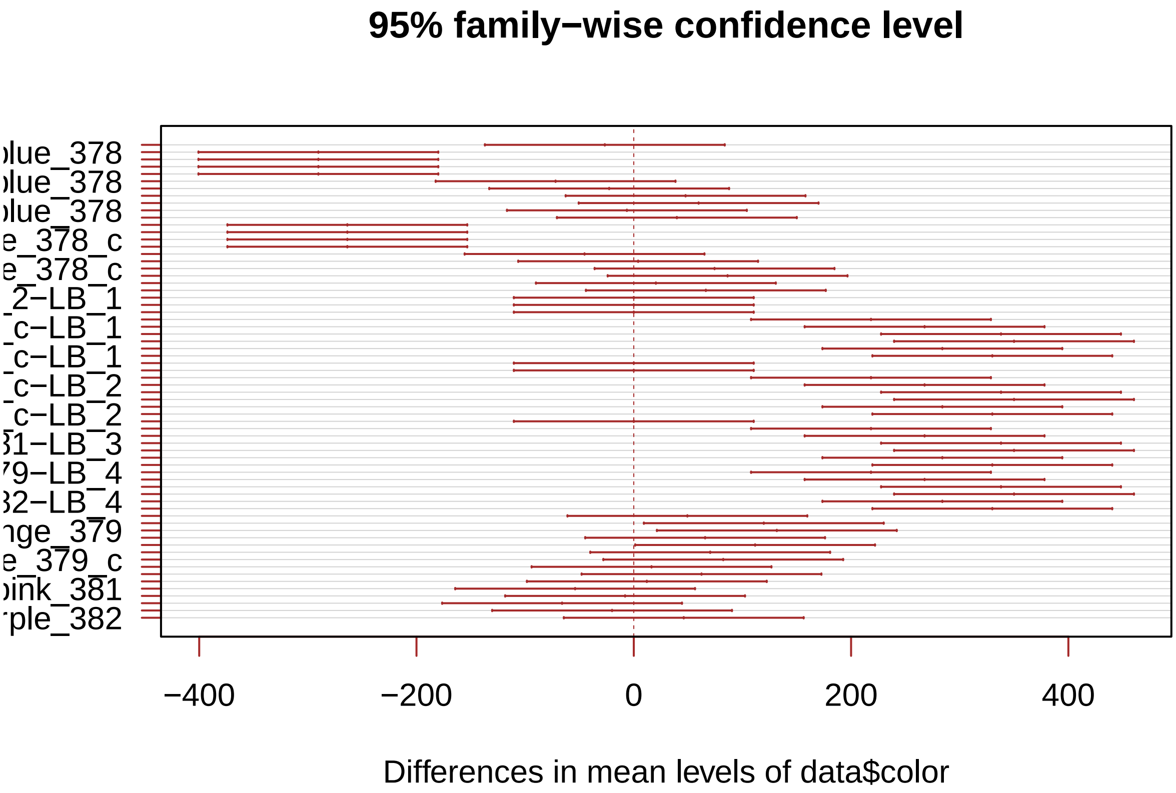


```
# I need to group the treatments that are not different each other together.
generate_label_df <- function(TUKEY, variable){

     # Extract labels and factor levels from Tukey post-hoc
     Tukey.levels <- TUKEY[[variable]][,4]
     Tukey.labels <- data.frame(multcompLetters(Tukey.levels)['Letters'])

     #I need to put the labels in the same order as in the boxplot :
     Tukey.labels$treatment=rownames(Tukey.labels)
     Tukey.labels=Tukey.labels[order(Tukey.labels$treatment) , ]
     return(Tukey.labels)
     }

# Apply the function on my dataset
LABELS <- generate_label_df(TUKEY , "data$color")

colnames(LABELS)<- c("Letters","color")
dataa <- merge(data, LABELS, by = "color",
                  all.x = TRUE)
```

```
data$color <- factor(data$color, levels = c("blue_378_c",'blue_378','LB_1',
                                            'orange_379_c','orange_379', 'LB_2','pink_381_c','pink_381'

Count_summary <- aggregate(.~color,data=data,FUN = mean )
colnames(Count_summary) <- c("color", "mean_count","control")
test1 <- aggregate(.~color,data=data,FUN="sd")
colnames(test1) <- c("color", "sd","control")
Count_summary["sd"] <- test1["sd"]
```

```r
data1 <- subset(dataa, select = -c(count,control))
data1 <- distinct(data1, color, .keep_all = TRUE)
Count_summary <- merge(Count_summary, data1, by = "color",
                  all.x = TRUE)

# create a vector with letters in the desired order
x <- c("blue_378_c",'blue_378','LB_1','orange_379_c','orange_379', 'LB_2','pink_381_c','pink_381','LB_3

Count_summary<- Count_summary %>%
  slice(match(x, color))

Count_summary$control <- as.logical(Count_summary$control)
Count_summary$control <- factor(Count_summary$control, labels = c('Cleaned',
                                                                  'Fresh'))
Count_summary$color <- str_replace_all(Count_summary$color,'_c','')
Count_summary$color <- str_replace_all(Count_summary$color,'-','_')


labelconv<- c("blue_378" = "aeBlue","LB_1"= " LB\nplate 1",
              "orange_379"="Yukon","LB_2" = " LB\nplate 2",
              "pink_381" = "eforCP","LB_3"= " LB\nplate 3",
              "purple_382"="tsPurple","LB_4" = " LB\nplate 4")

ggplot(data = Count_summary, aes(x = color, y = mean_count,
                                 fill=fct_relevel(control,"Fresh"))) +
  geom_bar(stat = "identity", position = position_dodge(), alpha = 0.75) +
  geom_errorbar( aes(x=color, ymin=mean_count-sd, ymax=mean_count+sd),
                 width=0.4, colour="orange", alpha=0.9, size=1.3,
                 position = position_dodge(.9)) +
  scale_x_discrete(labels=labelconv,
                   limits=unique(as.character(Count_summary$color)) ) +
  geom_text(data = Count_summary, aes(x = color, y = 400, label = Letters,),
            size = 3.4, vjust = 0, hjust = -0.5,
            position =position_dodge(width = 1) ) +
  labs(y = "Average Colony Count", x = expression(paste(italic("E. coli"),
                                                        "Color")))+
  scale_fill_manual(values=c("gray","maroon","gray","gray","maroon","gray",
                             "gray","maroon","gray","gray","maroon","gray"))+
  theme_minimal() +labs(fill = "Tip treatment")+
  theme(axis.text.x = element_text(angle = 40, vjust = 0.5, hjust=.2))
```

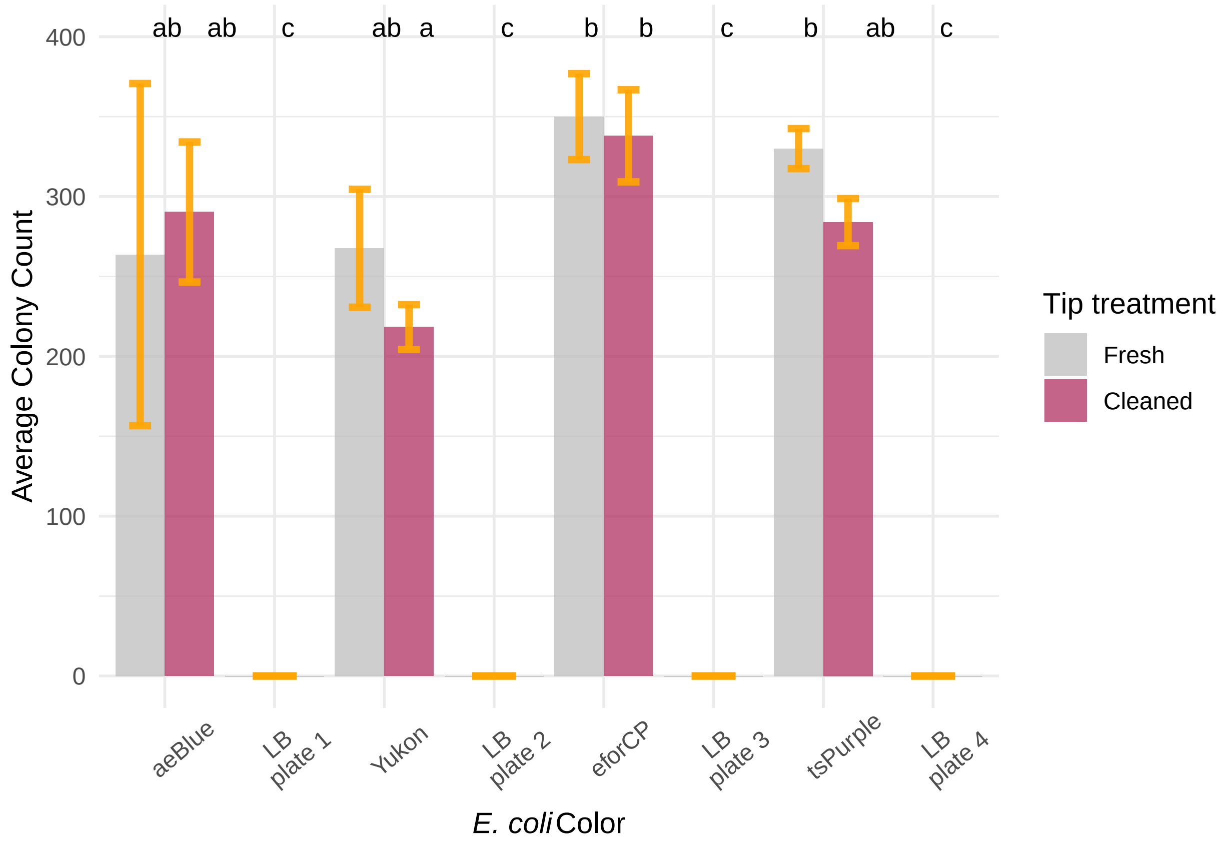


#Figure 2C

```
data <- read.csv("rawdata/20230418_vmax.csv")
data_long <- tidyr::pivot_longer(
  data = data, cols = c('one','two','three','four','five','six','seven',
                        'eight','nine','ten','eleven','twelve'),
  names_to = "Column")
```

Adding tip number and time to each row

```
simpletime <- c('maxv','r','tv','lag')
time1 <- unlist(lapply(simpletime, rep, 12))
clas<-rep(time1,8)

data_long['classification'] <- clas
data_long <-subset(data_long, data_long$value != "")

A <- rep('1-2-1',3)
B<- rep('1-1-1',3)
C<- rep('1-1',3)
D<- rep('1',3)
E<- rep('none',3)

trt1 <- c(A,B,C,D)
trt1<- rep(trt1,4)

trt2 <- c(E,A,B,C)
trt2<- rep(trt2,4)
```

```
trt3 <- c(D,E,A,B)
trt3<- rep(trt3,4)

trt4 <- c(C,D,E,A)
trt4<- rep(trt4,4)

trt5 <- c(B,C,D,E)
trt5<- rep(trt5,4)

trt6 <- c(A,B,C,D)
trt6<- rep(trt6,4)

trt7 <- c(E,A,B,C)
trt7<- rep(trt7,4)

trt8 <- c(D,E)
trt8<- rep(trt8,4)

alltrts<-c(trt1,trt2,trt3,trt4,trt5,trt6,trt7,trt8)

one <- rep('1',48)

A <- rep('none',3)
B <-rep('7.5',9)
C<-c(A,B)
two<-rep(C,4)

A <- rep('7.5',3)
B <-rep('none',3)
C <-rep('1',6)
D<-c(A,B,C)
three<-rep(D,4)

A <- rep('1',6)
B <-rep('none',3)
C <-rep('7.5',3)
D<-c(A,B,C)
four<-rep(D,4)

A <- rep('7.5',9)
B <-rep('none',3)
D<-c(A,B)
five<-rep(D,4)

six<-rep('1',48)

A <- rep('none',3)
B <-rep('7.5',9)
D<-c(A,B)
seven<-rep(D,4)

A <- rep('7.5',3)
B <-rep('none',3)
```

```
D<-c(A,B)
eight<-rep(D,4)

allblch <- c(one,two,three,four,five,six,seven,eight)

data_long['treatment']<- alltrts
data_long['bleach']<-allblch
data_long$combo<-paste(data_long$treatment,data_long$bleach,sep="_")
```

Here you read in the OD600 values, along with the “Time” Column

```
data <- read.csv("rawdata/20230418_od.csv")

data_long1 <- tidyr::pivot_longer(
  data = data, cols = c('A1','A2','A3','A4','A5','A6','A7','A8','A9','A10',
                        'A11','A12','B1','B2','B3','B4','B5','B6','B7','B8',
                        'B9','B10','B11','B12','C1','C2','C3','C4','C5','C6',
                        'C7','C8','C9','C10','C11','C12','D1','D2','D3','D4',
                        'D5','D6','D7','D8','D9','D10','D11','D12','E1','E2',
                        'E3','E4','E5','E6','E7','E8','E9','E10','E11','E12',
                        'F1','F2','F3','F4','F5','F6','F7','F8','F9','F10',
                        'F11','F12','G1','G2','G3','G4','G5','G6','G7','G8',
                        'G9','G10','G11','G12','H1','H2','H3','H4','H5','H6'),
  names_to = "Column")

wells<-c('A1','A2','A3','A4','A5','A6','A7','A8','A9','A10','A11','A12','B1',
         'B2','B3','B4','B5','B6','B7','B8','B9','B10','B11','B12','C1','C2',
         'C3','C4','C5','C6','C7','C8','C9','C10','C11','C12','D1','D2','D3',
         'D4','D5','D6','D7','D8','D9','D10','D11','D12','E1','E2','E3','E4',
         'E5','E6','E7','E8','E9','E10','E11','E12','F1','F2','F3','F4','F5',
         'F6','F7','F8','F9','F10','F11','F12','G1','G2','G3','G4','G5','G6',
         'G7','G8','G9','G10','G11','G12','H1','H2','H3','H4','H5','H6' )

A <- rep('1-2-1',3)
B<- rep('1-1-1',3)
C<- rep('1-1',3)
D<- rep('1',3)
E<- rep('none',3)

trt1 <- c(A,B,C,D)
trt2 <- c(E,A,B,C)
trt3 <- c(D,E,A,B)
trt4 <- c(C,D,E,A)
trt5 <- c(B,C,D,E)
trt6 <- c(A,B,C,D)
trt7 <- c(E,A,B,C)
trt8 <- c(D,E)

alltrts<-c(trt1,trt2,trt3,trt4,trt5,trt6,trt7,trt8)
alltrts<-rep(alltrts,27)
alltrts<-as.data.frame((alltrts))

one<- rep('1',12)
two<-rep('none',3)
```

```
three<-rep('7.5',12)
four<-rep('none',3)
comb<-c(one,two,three,four)
comball<-rep(comb,3)
allblch<-rep(comball,27)
allblch<- as.data.frame(allblch)

alltrts$combo<-paste(alltrts[,1],allblch[,1],sep="_")
vec<-alltrts$combo

data_long1['trt']<-vec
simpletime <- c(1,2,3,4,5,6,7,8,9,10,11,12,13,14,15,16,17,18,19,20,21,22,23,24,
                25,26,27)
time1 <- unlist(lapply(simpletime, rep,90))
data_long1['time'] <- time1
```

Here I'm selecting only growth data before Vmax to accurately fit an exponential growth model

```
allgrowthconstants <- c()
allxo <- c()

data_long$Column <- gsub("one", 1, data_long$Column)
data_long$Column <- gsub("two", 2, data_long$Column)
data_long$Column <- gsub("three", 3, data_long$Column)
data_long$Column <- gsub("four", 4, data_long$Column)
data_long$Column <- gsub("five", 5, data_long$Column)
data_long$Column <- gsub("six", 6, data_long$Column)
data_long$Column <- gsub("seven", 7, data_long$Column)
data_long$Column <- gsub("eight", 8, data_long$Column)
data_long$Column <- gsub("nine", 9, data_long$Column)
data_long$Column <- gsub("ten", 10, data_long$Column)
data_long$Column <- gsub("eleven", 11, data_long$Column)
data_long$Column <- gsub("twelve", 12, data_long$Column)
data_long$well <-paste(data_long$X,data_long$Column,sep="")
data_long$wellncol <-paste(data_long$well,data_long$combo,sep="+")
data_long1$trtncol <-paste(data_long1$Column,data_long1$trt,sep="+")

vec <- data_long1$trtncol
vec <- unique(vec)

for (x in vec) {
tipdat<- subset(data_long1, trtncol == x )
growdat<-subset(data_long, wellncol== x  )
growdat<-subset(growdat, classification=="tv")
time <- gsub(':','',growdat$value)
tipdat$Time<-gsub(':','',tipdat$Time)
growdat$value<-gsub(':','',growdat$value)
tipdat$use = ""

for (row in 1:nrow(tipdat)) {

  if (as.numeric(tipdat[row,"Time"]) - (as.numeric(growdat[1,"value"])) < 0){
  tipdat[row,"use"] <- "yes"}
  else {
```

```
    tipdat[row,"use"] <-"nope"
  }
}
tipdat <- subset(tipdat, tipdat$use == 'yes')

relation <- lm(log(tipdat$value)~tipdat$time)
intermediate <- coef(relation)
growthconstant <- intermediate[2]
xo <- exp(intermediate[1])
allgrowthconstants <-append(allgrowthconstants, growthconstant)
allxo <-append(allxo,xo)
}

growthefficiency1 <- data.frame(vec,allgrowthconstants,allxo)
```

```
ge2<-growthefficiency1
ge2[c('well', 'treatment')] <- str_split_fixed(ge2$vec, '\\+', 2)
ge2[c('trtmnt', 'bleach')] <- str_split_fixed(ge2$treatment, '_', 2)

ge2$treatment <- sub('-','_',ge2$treatment)
ge2$treatment <- sub('-','_',ge2$treatment)

model=lm(allxo ~ treatment, data = ge2)
ANOVA=aov(model)
summary(ANOVA)
```

```
##             Df   Sum Sq   Mean Sq F value  Pr(>F)
## treatment    8 0.001411 1.764e-04   3.405 0.00202 **
## Residuals   81 0.004197 5.181e-05
## ---
## Signif. codes:  0 '***' 0.001 '**' 0.01 '*' 0.05 '.' 0.1 ' ' 1
```

```
TUKEY <- TukeyHSD(x=ANOVA, 'treatment', conf.level=0.95)
plot(TUKEY , las=1 , col="brown")
```

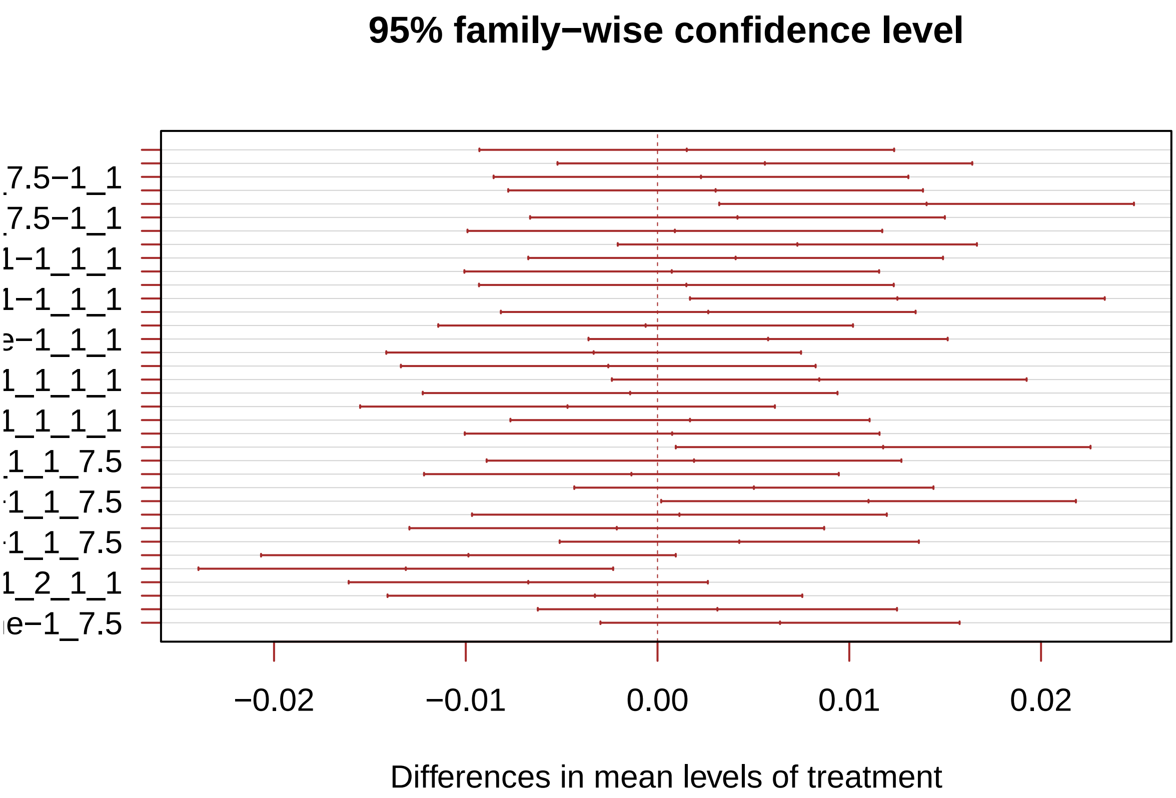


```
# I need to group the treatments that are not different each other together.
generate_label_df <- function(TUKEY, variable){

     # Extract labels and factor levels from Tukey post-hoc
     Tukey.levels <- TUKEY[[variable]][,4]
     Tukey.labels <- data.frame(multcompLetters(Tukey.levels)['Letters'])

     #I need to put the labels in the same order as in the boxplot :
     Tukey.labels$treatment=rownames(Tukey.labels)
     Tukey.labels=Tukey.labels[order(Tukey.labels$treatment) , ]
     return(Tukey.labels)
     }

# Apply the function on my dataset
#LABELS <- generate_label_df(TUKEY , "steady_states$strain_and_trt")
LABELS <- generate_label_df(TUKEY , "treatment")
names(LABELS)[2] ="combo"

ge2["combo"] <-""
ge2$combo<-ge2$treatment
ge2 <- merge(ge2, LABELS, by = "combo",
                  all.x = TRUE)

ggplot(ge2, aes(x =trtmnt, y = allxo, fill=bleach)) +
  geom_boxplot(outlier.shape = NA) +
  geom_point(alpha = 0.7, position = position_beeswarm(dodge.width = 0.75)) +
  scale_y_continuous(trans=scales::pseudo_log_trans(base = 10)) +
  labs(y = "Xo") +
  scale_color_manual(values = c("darkgrey", "maroon","green")) +
```

```
  scale_fill_manual(values = c("darkgrey", "maroon","green")) +
  theme_classic() +
  theme(
        axis.text.x = element_text(angle = 30, hjust = 1, vjust = 1))+
  geom_text(data = ge2, aes(x = trtmnt , y = .155, label = Letters),
            size = 3.4, vjust = 0, hjust = -0.5,
            position =position_dodge(width = 1) )+
  labs(x="Rinse Rigor")+ labs(fill = "[Bleach]")
```

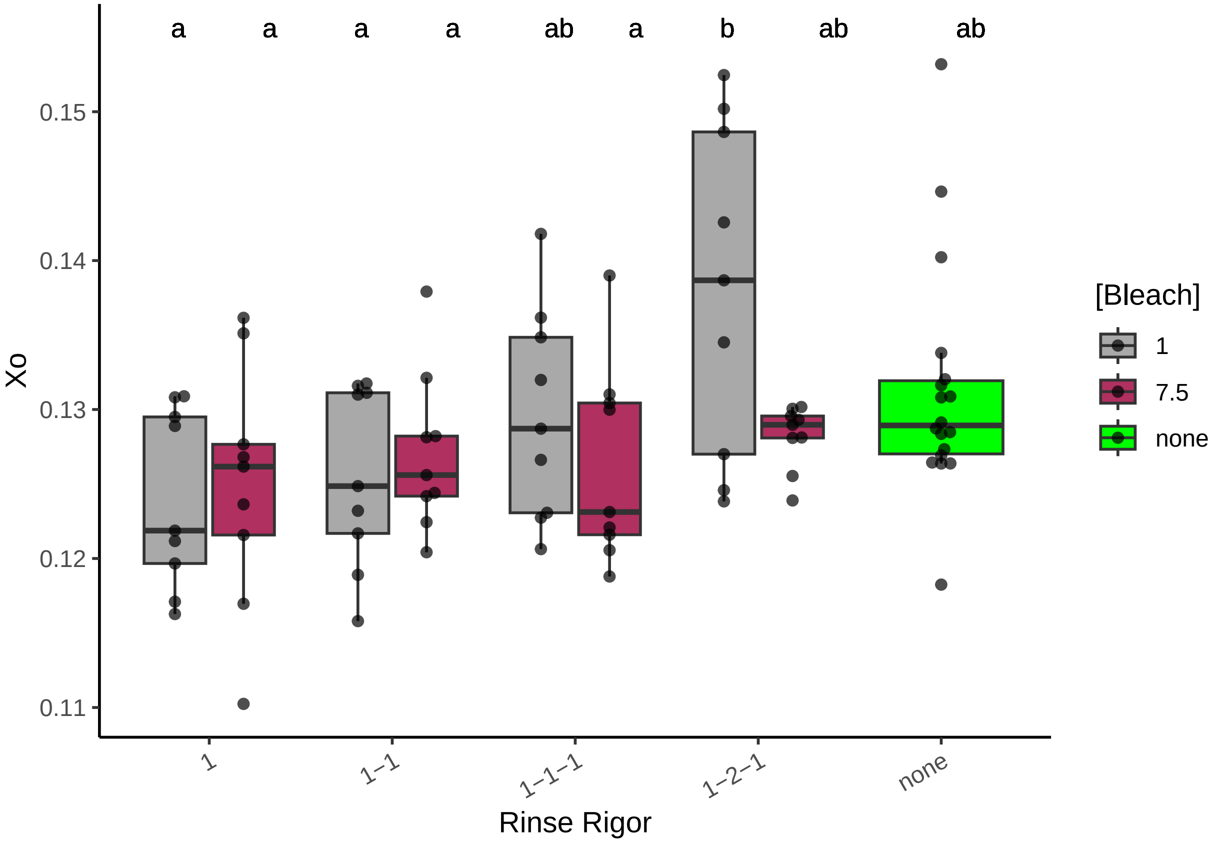


```
#ggsave("fig2C.png", width = 8, height = 4, dpi = 400)
```

#Figure 2D

```
data <- read.csv("rawdata/20230420_vmax.csv")
data_long <- tidyr::pivot_longer(
  data = data, cols = c('one','two','three','four','five','six','seven',
                        'eight','nine','ten','eleven','twelve'),
  names_to = "Column")
```

Adding tip number and time to each row

```
simpletime <- c('maxv','r','tv','lag')
time1 <- unlist(lapply(simpletime, rep, 12))
clas<-rep(time1,8)

data_long['classification'] <- clas
data_long <-subset(data_long, data_long$value != "")
```

```
A <- rep('clean',6)
B<- rep('fresh',6)
C <- c(A,B)
E<- rep(C,32)

data_long['treatment']<- E
data_long$combo<-paste(data_long$treatment,data_long$bleach,sep="_")
```

```
## Warning: Unknown or uninitialised column: `bleach`.
```

Here you read in the OD600 values, along with the "Time" Column

```
data <- read.csv("rawdata/20230420_od.csv")
data_long1 <- tidyr::pivot_longer(
  data = data, cols = c('A1','A2','A3','A4','A5','A6','A7','A8','A9','A10',
                        'A11','A12','B1','B2','B3','B4','B5','B6','B7','B8',
                        'B9','B10','B11','B12','C1','C2','C3','C4','C5','C6',
                        'C7','C8','C9','C10','C11','C12','D1','D2','D3','D4',
                        'D5','D6','D7','D8','D9','D10','D11','D12','E1','E2',
                        'E3','E4','E5','E6','E7','E8','E9','E10','E11','E12',
                        'F1','F2','F3','F4','F5','F6','F7','F8','F9','F10',
                        'F11','F12','G1','G2','G3','G4','G5','G6','G7','G8',
                        'G9','G10','G11','G12','H1','H2','H3','H4','H5','H6',
                        'H7','H8','H9','H10','H11','H12'), names_to = "Column")

wells<-c('A1','A2','A3','A4','A5','A6','A7','A8','A9','A10','A11','A12','B1',
         'B2','B3','B4','B5','B6','B7','B8','B9','B10','B11','B12','C1','C2',
         'C3','C4','C5','C6','C7','C8','C9','C10','C11','C12','D1','D2','D3',
         'D4','D5','D6','D7','D8','D9','D10','D11','D12','E1','E2','E3','E4',
         'E5','E6','E7','E8','E9','E10','E11','E12','F1','F2','F3','F4','F5',
         'F6','F7','F8','F9','F10','F11','F12','G1','G2','G3','G4','G5','G6',
         'G7','G8','G9','G10','G11','G12','H1','H2','H3','H4','H5','H6','H7',
         'H8','H9','H10','H11','H12' )

E<- rep(C,216)
data_long1['treatment']<- E

simpletime <- c(1,2,3,4,5,6,7,8,9,10,11,12,13,14,15,16,17,18,19,20,21,22,23,24,
                25,26,27)
time1 <- unlist(lapply(simpletime, rep,96))
data_long1['time'] <- time1
```

Here I'm selecting only growth data before Vmax to accurately fit an exponential growth model

```
allgrowthconstants <- c()
allxo <- c()

data_long$Column <- gsub("one", 1, data_long$Column)
data_long$Column <- gsub("two", 2, data_long$Column)
data_long$Column <- gsub("three", 3, data_long$Column)
data_long$Column <- gsub("four", 4, data_long$Column)
data_long$Column <- gsub("five", 5, data_long$Column)
data_long$Column <- gsub("six", 6, data_long$Column)
data_long$Column <- gsub("seven", 7, data_long$Column)
data_long$Column <- gsub("eight", 8, data_long$Column)
```

```r
data_long$Column <- gsub("nine", 9, data_long$Column)
data_long$Column <- gsub("ten", 10, data_long$Column)
data_long$Column <- gsub("eleven", 11, data_long$Column)
data_long$Column <- gsub("twelve", 12, data_long$Column)
data_long$well <-paste(data_long$X,data_long$Column,sep="")
data_long$wellncol <-paste(data_long$well,data_long$combo,sep="+")
data_long1$trtncol <-paste(data_long1$Column,data_long1$trt,sep="+")
```

```
## Warning: Unknown or uninitialised column: `trt`.
```

```r
vec <- data_long1$trtncol
vec <- unique(vec)

for (x in wells) {
tipdat<- subset(data_long1, Column == x )
growdat<-subset(data_long, well== x  )
growdat<-subset(growdat, classification=="tv")
time <- gsub(':','',growdat$value)
tipdat$Time<-gsub(':','',tipdat$Time)
growdat$value<-gsub(':','',growdat$value)
tipdat$use = ""

for (row in 1:nrow(tipdat)) {

  if (as.numeric(tipdat[row,"Time"]) - (as.numeric(growdat[1,"value"])) < 0){
  tipdat[row,"use"] <- "yes"}
  else {
    tipdat[row,"use"] <-"nope"
  }
}
tipdat <- subset(tipdat, tipdat$use == 'yes')

relation <- lm(log(tipdat$value)~tipdat$time)
intermediate <- coef(relation)
growthconstant <- intermediate[2]
xo <- exp(intermediate[1])
allgrowthconstants <-append(allgrowthconstants, growthconstant)
allxo <-append(allxo,xo)
}

growthefficiency1 <- data.frame(wells,allgrowthconstants,allxo)
A <- rep('clean',6)
B<- rep('fresh',6)
C <- c(A,B)
E<- rep(C,8)
growthefficiency1$clean <- E
```

```r
ggplot(growthefficiency1, aes(x =clean, y = allxo,fill=clean)) +
  geom_boxplot(outlier.shape = NA) +
  geom_point(alpha = 0.7, position = position_beeswarm(dodge.width = 0.75)) +
  scale_y_continuous(trans=scales::pseudo_log_trans(base = 10)) +
  labs(y = "Xo") +
  scale_color_manual(values = c("darkgrey", "maroon")) +
  scale_fill_manual(values = c("darkgrey", "maroon")) +
```

```
  theme_classic() +
  theme(legend.position = c(.9,.15),
        axis.text.x = element_text(angle = 30, hjust = 1, vjust = 1))+
  stat_compare_means(method = "t.test",label.x = 1.4, label.y = .028)
```

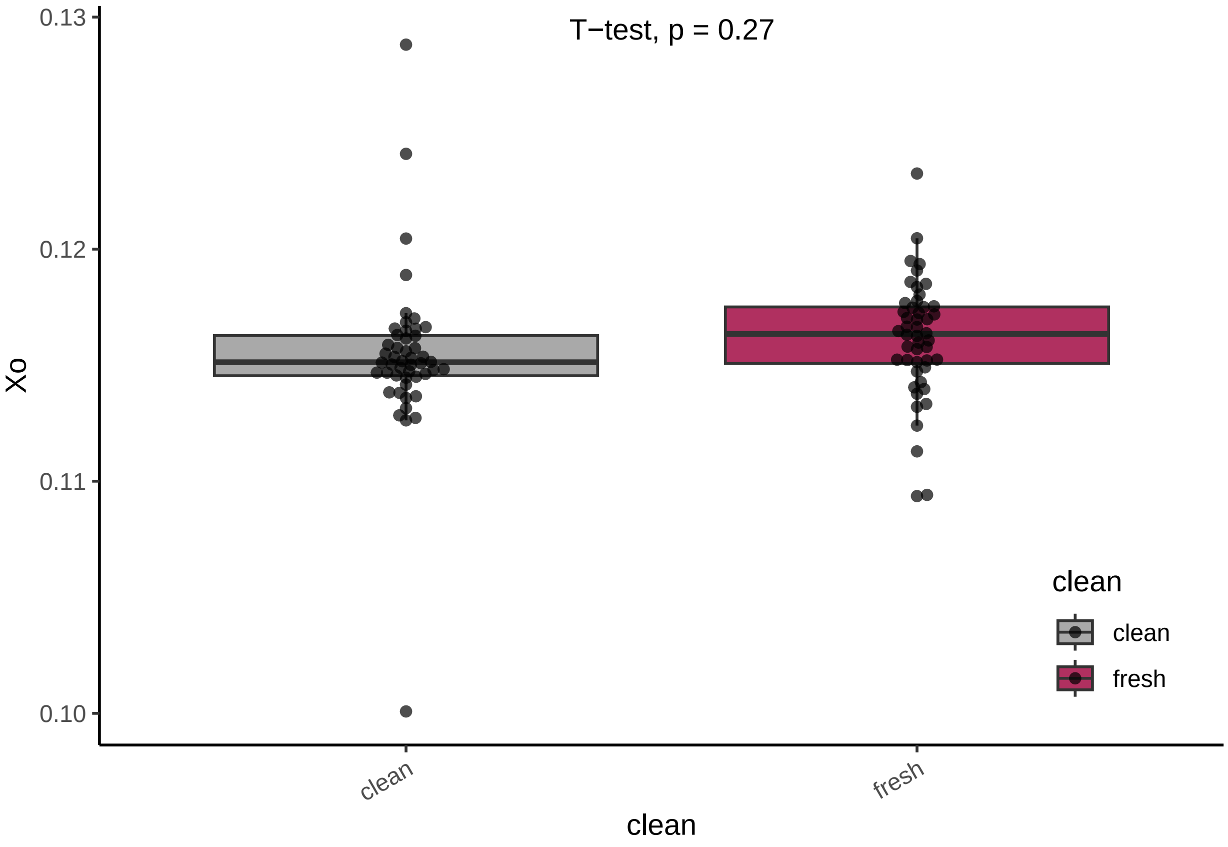


CV calculations

```
ge2<-growthefficiency1
ge <- subset(ge2, select = -c(wells))

means <- aggregate(.~clean,data=ge,FUN = "mean" )
colnames(means) <- c("clean", "meanug","meanxo")
sd <- aggregate(.~clean,data=ge,FUN ="sd")
colnames(sd) <- c("clean", "sdug","sdxo")
means["sdug"] <- sd["sdug"]
means["sdxo"] <- sd["sdxo"]
means["cv"] <- (means["sdxo"]/means["meanxo"])*100
print(means)
```

```
##   clean    meanug    meanxo        sdug        sdxo       cv
## 1 clean 0.1349043 0.1154284 0.011606199 0.003542011 3.068579
## 2 fresh 0.1394889 0.1161361 0.004367283 0.002563587 2.207399
```

#Figure 3A

```
data<- read.csv("rawdata/20230417_tempremoved.csv")
newdata <- subset(data, select = -c(Well,Cq))
cl<-c("fresh","dirty","clean","clean","clean")
cll<-rep(cl,6)
```

```
newdata$Cleaning<-cll
ggplot(newdata, aes( y = SQ, x=fct_relevel(Sample, "fresh"),
                     color=Cleaning,fill=Cleaning)) +
  geom_boxplot(outlier.shape = NA, fill=c("gray","gray","gray","gray","maroon"),
               alpha=0.3) +
  geom_point(alpha = 0.7, position = position_beeswarm(dodge.width = 0.75)) +
  scale_y_continuous(trans=scales::pseudo_log_trans(base = 10)) +
  labs(y = "DNA Contamination (ng)") +
  scale_color_manual(values = c("darkgray", "maroon","orange")) +
  scale_fill_manual(values = c("darkgray", "maroon","orange")) +
  theme_classic() +
  theme(legend.position = c(.3, .7),
        axis.text.x = element_text(angle = 30, hjust = 1, vjust = 1),
        axis.title.x = element_blank())
```

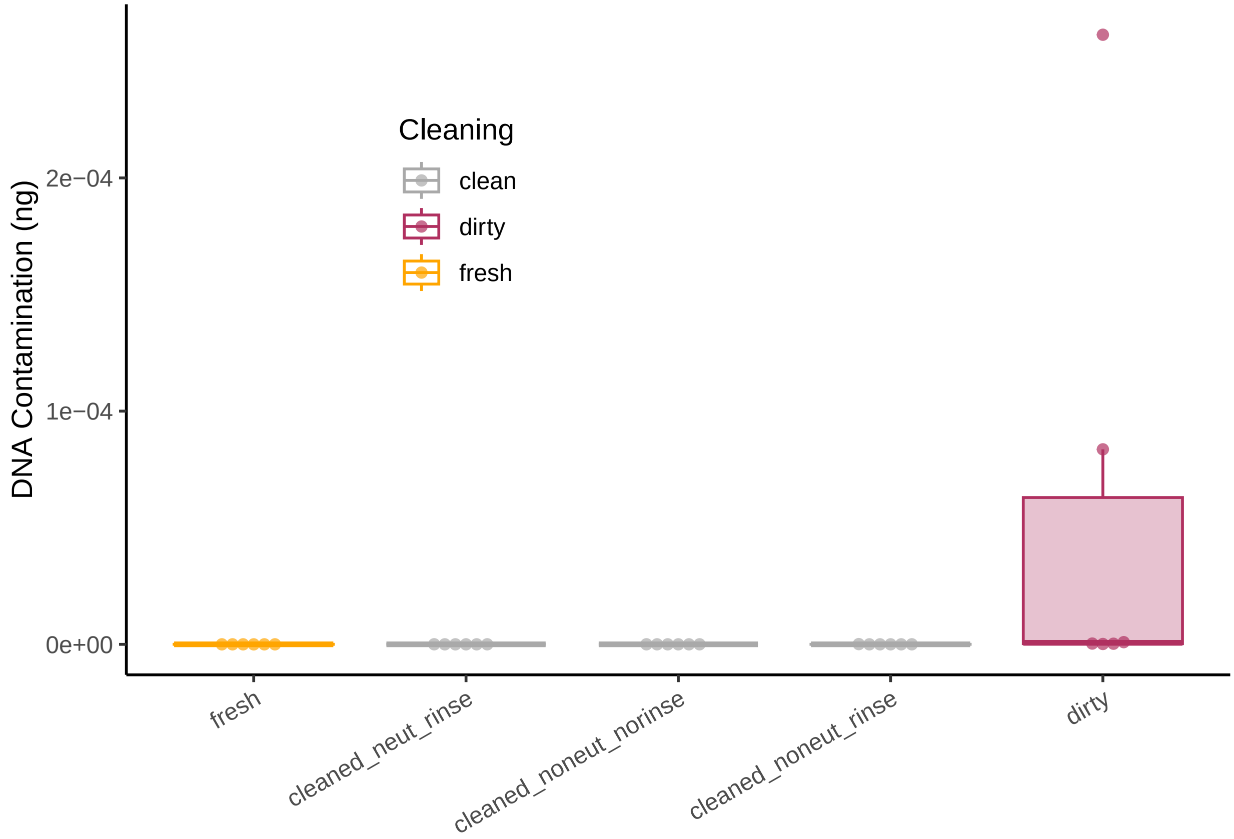


#Figure 4A

```
data<- read.csv("rawdata/20230414_qpcr_analyzed.csv")
newdata <- subset(data, Sample != "dirty_notemp" )
newdata <- subset(newdata, Sample != "notemp_control")
newdata <- subset(newdata,  Sample != "dirty_normalefficiencytest")
newdata <- subset(newdata, Sample != "dirty_noprim" )
newdata<- newdata %>%
  mutate(across(SQ, na_if, "#VALUE!")) %>%
  mutate(across(Cq, na_if, "Undetermined"))
```

```
## Warning: There was 1 warning in `mutate()`.
## i In argument: `across(SQ, na_if, "#VALUE!")`.
```

```
## Caused by warning:
## ! The `...` argument of `across()` is deprecated as of dplyr 1.1.0.
## Supply arguments directly to `.fns` through an anonymous function instead.
##
##   # Previously
##   across(a:b, mean, na.rm = TRUE)
##
##   # Now
##   across(a:b, \(x) mean(x, na.rm = TRUE))
```

```
model=lm( newdata$SQ ~ newdata$Sample )
ANOVA=aov(model)
summary(ANOVA)
```

```
##                 Df Sum Sq Mean Sq F value Pr(>F)
## newdata$Sample  8  3.596  0.4495   436.2 <2e-16 ***
## Residuals      18  0.019  0.0010
## ---
## Signif. codes:  0 '***' 0.001 '**' 0.01 '*' 0.05 '.' 0.1 ' ' 1
## 9 observations deleted due to missingness
```

```
TUKEY <- TukeyHSD(x=ANOVA, 'newdata$Sample', conf.level=0.95)
cld <- multcompLetters4(ANOVA, TUKEY)
```

```
## Warning in mean.default(x = c("0.746129899", "0.74857267", "0.780089315"):
## argument is not numeric or logical: returning NA

## Warning in mean.default(x = c("0.738625495", "0.811492642", "0.803149803"):
## argument is not numeric or logical: returning NA

## Warning in mean.default(x = c("4.14624E-08", "4.7956E-09", "1.579E-09")):
## argument is not numeric or logical: returning NA

## Warning in mean.default(x = c("0.748578333", "0.722145282", "0.649527855"):
## argument is not numeric or logical: returning NA

## Warning in mean.default(x = c("0.87948675", "0.862281497", "0.826530988"):
## argument is not numeric or logical: returning NA

## Warning in mean.default(x = c("2.82865E-10", "1.51653E-10", "3.78423E-09"):
## argument is not numeric or logical: returning NA

## Warning in mean.default(x = c("0.76456393", "0.665300679", "0.701425267"):
## argument is not numeric or logical: returning NA

## Warning in mean.default(x = c("0.770641718", "0.841135084", "0.781468506"):
## argument is not numeric or logical: returning NA

## Warning in mean.default(x = c("4.1721E-10", "2.27137E-09", "1.24226E-10"):
## argument is not numeric or logical: returning NA
```

```
print(cld)
```

```
## $`newdata$Sample`
##        base_wash_efficiency           base_wash_initial
##                         "a"                        "ab"
##            base_wash_notemp no_neut_or_rinse_efficiency
##                         "c"                         "a"
##    no_neut_or_rinse_initial     no_neut_or_rinse_notemp
##                         "b"                         "c"
```

```
##   no_neutralizers_efficiency     no_neutralizers_initial
##                          "a"                        "ab"
##       no_neutralizers_notemp
##                          "c"
```

```r
# Tuckey test representation :
plot(TUKEY , las=1 , col="brown")
```

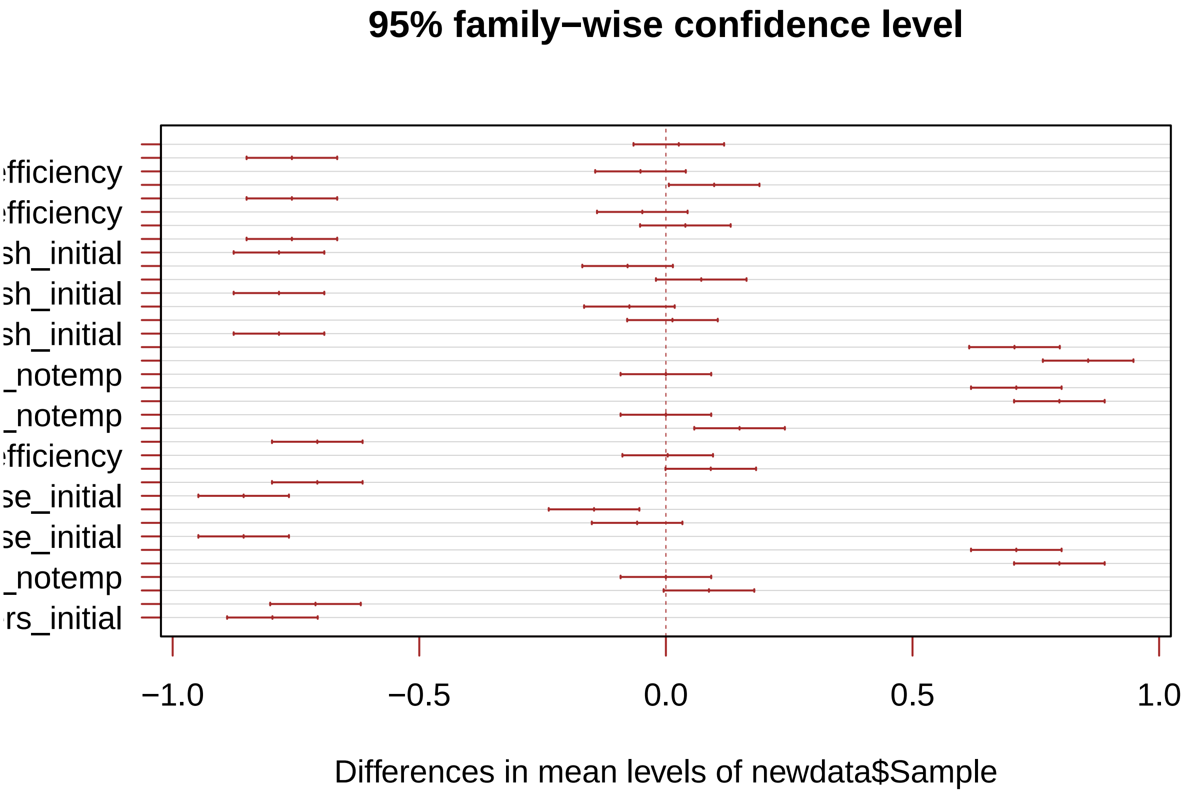


```r
# I need to group the treatments that are not different each other together.
generate_label_df <- function(TUKEY, variable){

     # Extract labels and factor levels from Tukey post-hoc
     Tukey.levels <- TUKEY[[variable]][,4]
     Tukey.labels <- data.frame(multcompLetters(Tukey.levels)['Letters'])

     #I need to put the labels in the same order as in the boxplot :
     Tukey.labels$treatment=rownames(Tukey.labels)
     Tukey.labels=Tukey.labels[order(Tukey.labels$treatment) , ]
     return(Tukey.labels)
     }

# Apply the function on my dataset
LABELS <- generate_label_df(TUKEY , "newdata$Sample")
colnames(LABELS)<- c("Letters","Sample")

newdata <- subset(newdata, select = -c(Cq))
newdata[is.na(newdata)] <- 0
newdata$SQ = as.numeric(as.character(newdata$SQ))

means <- aggregate(.~Sample,data=newdata,FUN = "mean" )
```

```
colnames(means) <- c("Sample", "meanSQ")
sd <- aggregate(.~Sample,data=newdata,FUN="sd")
colnames(sd) <- c("Sample", "sd")
means["sd"] <- sd["sd"]
means <- merge(means, LABELS, by = "Sample",
                  all.x = TRUE)
x <- c("base_wash_initial","no_neutralizers_initial",
       "no_neut_or_rinse_initial","base_wash_notemp","no_neutralizers_notemp",
       "no_neut_or_rinse_notemp","base_wash_noprim","no_neutralizers_noprim",
       "no_neut_or_rinse_noprim","base_wash_efficiency",
       "no_neutralizers_efficiency","no_neut_or_rinse_efficiency")
means<- means %>%
  slice(match(x, Sample))
means$Sample <- factor(means$Sample, levels = means$Sample)

cl<-c("Fresh","Fresh","Fresh","Cleaned","Cleaned","Cleaned","Cleaned",
      "Cleaned","Cleaned","Cleaned","Cleaned","Cleaned")
means$Cleaned <- cl

labelconv<- c("base_wash_initial" = "Wash+Neutralize\n+Rinse",
              "no_neutralizers_initial"= "Wash+Rinse",
              "no_neut_or_rinse_initial"="Wash","base_wash_notemp" = "Wash+Neutralize\n+Rinse","no_neut
              "no_neut_or_rinse_efficiency"="Wash")

ggplot(data = means, aes(x = Sample, y = meanSQ,fill=Cleaned)) +
  geom_bar(stat = "identity", position = position_dodge(), alpha = 0.75) +
  geom_errorbar( aes(x=Sample, ymin=meanSQ-sd, ymax=meanSQ+sd),
                 width=0.4, colour="orange", alpha=0.9, size=1.3,
                 position = position_dodge(.9))   +
  geom_text(data = means, aes(x = Sample, y = meanSQ+.1, label = Letters,),
            size = 5, vjust = 0, hjust = -0.5,
            position =position_dodge(width = 1) ) +
  labs(y = "Amplification Efficiency", x = "Wash Rigor" )+ scale_fill_manual(values=c("maroon","gray","
  theme_minimal() +
  scale_x_discrete(labels=labelconv)+
  theme(axis.text.x = element_text(angle = 90, vjust = 0.5, hjust=.2))
```

```
## Warning: Removed 3 rows containing missing values (`geom_text()`).
```

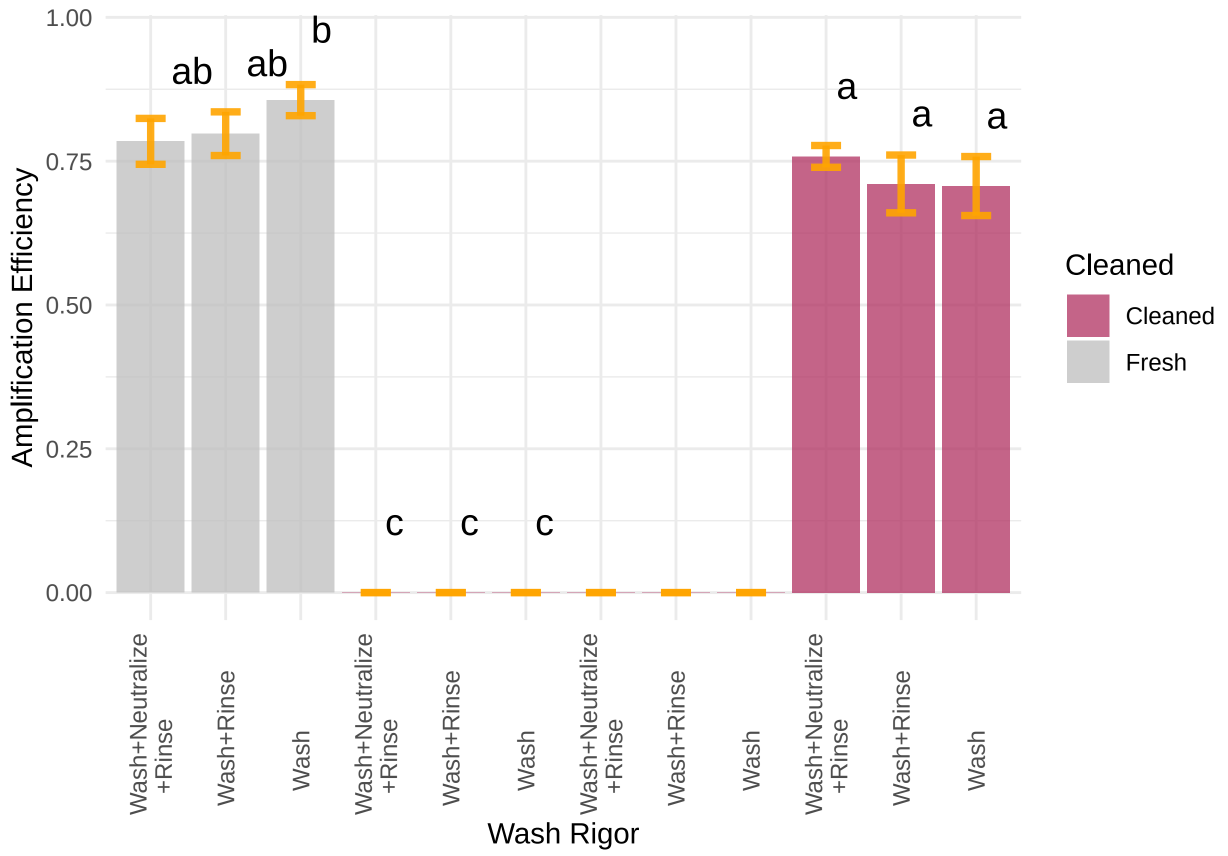


#Figure 5C reading in and annotating data

```
data1a <- read.csv("rawdata/20230215_plateclean.csv")
data2a <- read.csv("rawdata/20230403_plateclean.csv")
data3a <- read.csv("rawdata/20230406_plateclean.csv")
data4a <- read.csv("rawdata/20230412_plateclean.csv")

data1a <- data1a[!grepl("All Events", data1a$Gate),] #taking away an
#unnecessary datapoint

data1a[43,"Sample"] <- "C2"
data1a[44,"Sample"] <- "C2"
data1a <- data1a[order(data1a$Sample),]
data1a$rep<-'1'

data2a <- distinct(data2a)
data2a[97,"Sample"] <- "C2"
data2a[98,"Sample"] <- "C2"
data2a <- data2a[!grepl("All Events", data2a$Gate),]
data2a <- data2a[!grepl(4, data2a$Sample),]
data2a <- data2a[!grepl(5, data2a$Sample),]
data2a <- data2a[!grepl(9, data2a$Sample),]
data2a <- data2a[!grepl(10, data2a$Sample),]
data2a <- data2a[order(data2a$Sample),]
data2a$rep<-'2'
```

```r
data3a <- data3a[!grepl("All Events", data3a$Gate),]
data3a<- data3a %>%
  mutate(Gate = str_replace(Gate, "R1", "Debris"))
data3a<- data3a %>%
  mutate(Gate = str_replace(Gate, "R2", "Cells"))
data3a[41,"Sample"] <- "C2"
data3a[42,"Sample"] <- "C2"
data3a <- data3a[!grepl("All Events", data3a$Gate),]
data3a <- data3a[!grepl(4, data3a$Sample),]
data3a <- data3a[!grepl(5, data3a$Sample),]
data3a <- data3a[!grepl(9, data3a$Sample),]
data3a <- data3a[!grepl(10, data3a$Sample),]
data3a <- data3a[order(data3a$Sample),]
data3a$rep<-'3'

data4a <- data4a[!grepl("All Events", data4a$Gate),]
data4a[37,"Sample"] <- "C2"
data4a[38,"Sample"] <- "C2"
data4a <- data4a[order(data4a$Sample),]
data4a$rep<-'4'

###############################################################################
#putting data in order and annotating
#data1 <- data1[seq(dim(data1)[1],1),]
clean_method <- c("Rinsee", "Rinse+Bleach","Bleach+H2O2+25degrinse",
                  "Bleach+H2O2+85degrinse","1%Bleach+H2O2+85degrinse")
clean_method <- rep(clean_method, each=12)
conts <- c("20X yeast","Bleached yeast","DI")
conts <- rep(conts, each=6)
clean_method <- append(clean_method,conts)

contents_b4_cleaned <- c("yeast","DI","yeast","DI","yeast","DI","yeast","DI",
                         "yeast","DI","20X yeast","Bleached yeast","none")
contents_b4_cleaned <- rep(contents_b4_cleaned, each=6)

data1a <- cbind(data1a, clean_method, contents_b4_cleaned)


#############################################################################
#data2 <- data2[seq(dim(data2)[1],1),]
clean_method <- c("Rinsee", "Rinse+Bleach","Bleach+H2O2+25degrinse",
                  "Bleach+H2O2+85degrinse","1%Bleach+H2O2+85degrinse",
                  "1%Bleach+H2O2+25degrinse","1%Bleach+H2O2+25degrinse")
clean_method <- rep(clean_method, each=12)
conts <- c("20X yeast","Bleached yeast")
conts <- rep(conts, each=6)
clean_method <- append(clean_method,conts)

contents_b4_cleaned <- c("yeast","DI","yeast","DI","yeast","DI","yeast","DI",
                         "yeast","DI","yeast","DI","yeast","DI","20X yeast",
                         "Bleached yeast")
contents_b4_cleaned <- rep(contents_b4_cleaned, each=6)
```

```
data2a <- cbind(data2a, clean_method, contents_b4_cleaned)
data2a <- data2a[!grepl("C2", data2a$Sample),] #there was no data collected
#for this point

############################################################################
#data3 <- data3[seq(dim(data3)[1],1),]
clean_method <- c("Bleach+H2O2+25degrinse","1%Bleach+H2O2+25degrinse",
                  "Bleach+.2Mbakingsoda+H2O2+25degrinse",
                  "Bleach+.2Mbakingsoda+H2O2+25degrinse")
clean_method <- rep(clean_method, each=12)
conts <- c("20X yeast","Bleached yeast")
conts <- rep(conts, each=6)
clean_method <- append(clean_method,conts)

contents_b4_cleaned <- c("yeast","DI","yeast","DI","yeast","DI","yeast","DI",
                         "20X yeast","Bleached yeast")
contents_b4_cleaned <- rep(contents_b4_cleaned, each=6)

data3a <- cbind(data3a, clean_method, contents_b4_cleaned)

######################################################################
#putting data in order and annotating
#data1 <- data1[seq(dim(data1)[1],1),]
clean_method <- c("Rinsee", "Rinse+Bleach","Bleach+H2O2+25degrinse",
                  "Bleach+H2O2+85degrinse","1%Bleach+H2O2+85degrinse",
                  "1%Bleach+H2O2+25degrinse")
clean_method <- rep(clean_method, each=12)

contents_b4_cleaned <- c("yeast","DI","yeast","DI","yeast","DI","yeast","DI",
                         "yeast","DI","yeast","DI")
contents_b4_cleaned <- rep(contents_b4_cleaned, each=6)

data4a <- cbind(data4a, clean_method, contents_b4_cleaned)
```

subsetting and grouping

```
newdata1 <- subset(data1a,contents_b4_cleaned == "yeast" |
                     contents_b4_cleaned == "none")
newdata2 <- subset(data2a, contents_b4_cleaned == "yeast" )
#newdata2["time_set"]<- ">7hrs"
newdata3 <- subset(data3a, contents_b4_cleaned == "yeast")
#newdata3["time_set"]<- "<7hrs"
newdata4 <- subset(data4a, contents_b4_cleaned == "yeast")
#newdata4["time_set"]<- ">7hrs"

neww<- rbind(newdata1,newdata2,newdata3,newdata4)
saveneww<-neww
```

Taking the top 3 highest event counts

```
a <- subset(neww, Gate == "Cells" )
b <- subset(neww, Gate == "Debris" )

data_new2 <- a %>%                            # Top N highest values by group
  arrange(desc(Count)) %>%
```

```
  group_by(clean_method) %>%
  slice(1:3)
data_new2
```

```
## # A tibble: 24 x 7
## # Groups:   clean_method [8]
##    Sample Gate  Count X.Total rep   clean_method              contents_b4_cleaned
##    <chr>  <chr> <int> <chr>   <chr> <chr>                     <chr>
##  1 F1     Cells     1 12.5    2     1%Bleach+H2O2+25degrinse yeast
##  2 F3     Cells     1 8.333   2     1%Bleach+H2O2+25degrinse yeast
##  3 B1     Cells     1 16.667  3     1%Bleach+H2O2+25degrinse yeast
##  4 E1     Cells     1 50.000  1     1%Bleach+H2O2+85degrinse yeast
##  5 E2     Cells     1 50.000  1     1%Bleach+H2O2+85degrinse yeast
##  6 E3     Cells     0 N/A     1     1%Bleach+H2O2+85degrinse yeast
##  7 D3     Cells     1 8.333   3     Bleach+.2Mbakingsoda+H2~ yeast
##  8 C1     Cells     0 N/A     3     Bleach+.2Mbakingsoda+H2~ yeast
##  9 C2     Cells     0 0.000   3     Bleach+.2Mbakingsoda+H2~ yeast
## 10 C3     Cells     2 100.000 1     Bleach+H2O2+25degrinse   yeast
## # i 14 more rows
```

```
datt<- b %>%                                  # Top N highest values by group
  arrange(desc(Count)) %>%
  group_by(clean_method) %>%
  slice(1:3)

yo<- rbind(data_new2,datt)
```

ANOVA and Tukey's

```
model=lm(Count ~ clean_method*Gate, data = yo )
ANOVA=aov(model)
summary(ANOVA)
```

```
##                   Df Sum Sq Mean Sq F value   Pr(>F)
## clean_method       7 502264   71752  27.510 8.04e-12 ***
## Gate               1  22620   22620   8.673  0.00598 **
## clean_method:Gate  7 542972   77567  29.739 2.81e-12 ***
## Residuals         32  83464    2608
## ---
## Signif. codes:  0 '***' 0.001 '**' 0.01 '*' 0.05 '.' 0.1 ' ' 1
```

```
TUKEY <- TukeyHSD(x=ANOVA, 'clean_method:Gate', conf.level=0.95)

# Tuckey test representation :
plot(TUKEY , las=1 , col="brown")
```

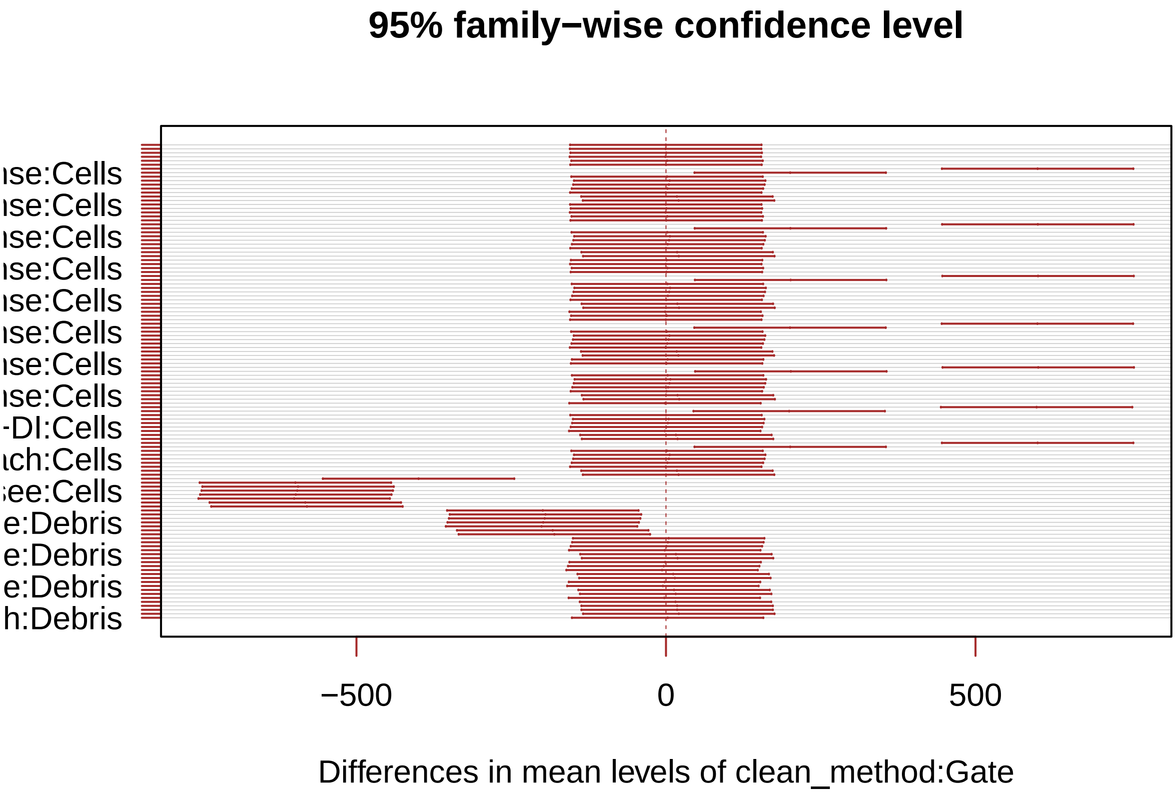


```
generate_label_df <- function(TUKEY, variable){

     # Extract labels and factor levels from Tukey post-hoc
     Tukey.levels <- TUKEY[[variable]][,4]
     Tukey.labels <- data.frame(multcompLetters(Tukey.levels)['Letters'])

     #I need to put the labels in the same order as in the boxplot :
     Tukey.labels$treatment=rownames(Tukey.labels)
     Tukey.labels=Tukey.labels[order(Tukey.labels$treatment) , ]
     return(Tukey.labels)
     }

# Apply the function on my dataset
#LABELS <- generate_label_df(TUKEY , "steady_states$strain_and_trt")
LABELS <- generate_label_df(TUKEY , "clean_method:Gate")

names(LABELS)[2] ="combo"

yo["combo"] <-""
yo$combo<-paste(yo$clean_method,yo$Gate,sep=":")
yo<- merge(yo, LABELS, by = "combo",
                  all.x = TRUE)

ggplot(yo, aes(x =fct_relevel(clean_method,"20X yeast","Bleached yeast", "Rinsee","Rinse+Bleach","Bleac
               y = Count, color = Gate)) +
  geom_boxplot(outlier.shape = NA) +
  geom_point(alpha = 0.7, position = position_beeswarm(dodge.width = 0.75)) +
  scale_y_continuous(trans=scales::pseudo_log_trans(base = 10)) +
  labs(y = "Cell Count") +
```

```
  scale_color_manual(values = c("darkgrey", "maroon")) +
  scale_fill_manual(values = c("darkgrey", "maroon")) +
  theme_classic() +
  theme(legend.position = c(.8, .7),
        axis.text.x = element_text(angle = 30, hjust = 1, vjust = 1),
        axis.title.x = element_blank())+
  geom_text(data = yo, aes(x = clean_method , y = 900, label = Letters),
            size = 3.4, vjust = 0, hjust = -0.5,
            position =position_dodge(width = 1) )+
  scale_x_discrete(labels=labelconv)
```

```
## Warning: 2 unknown levels in `f`: 20X yeast and Bleached yeast
## 2 unknown levels in `f`: 20X yeast and Bleached yeast
```

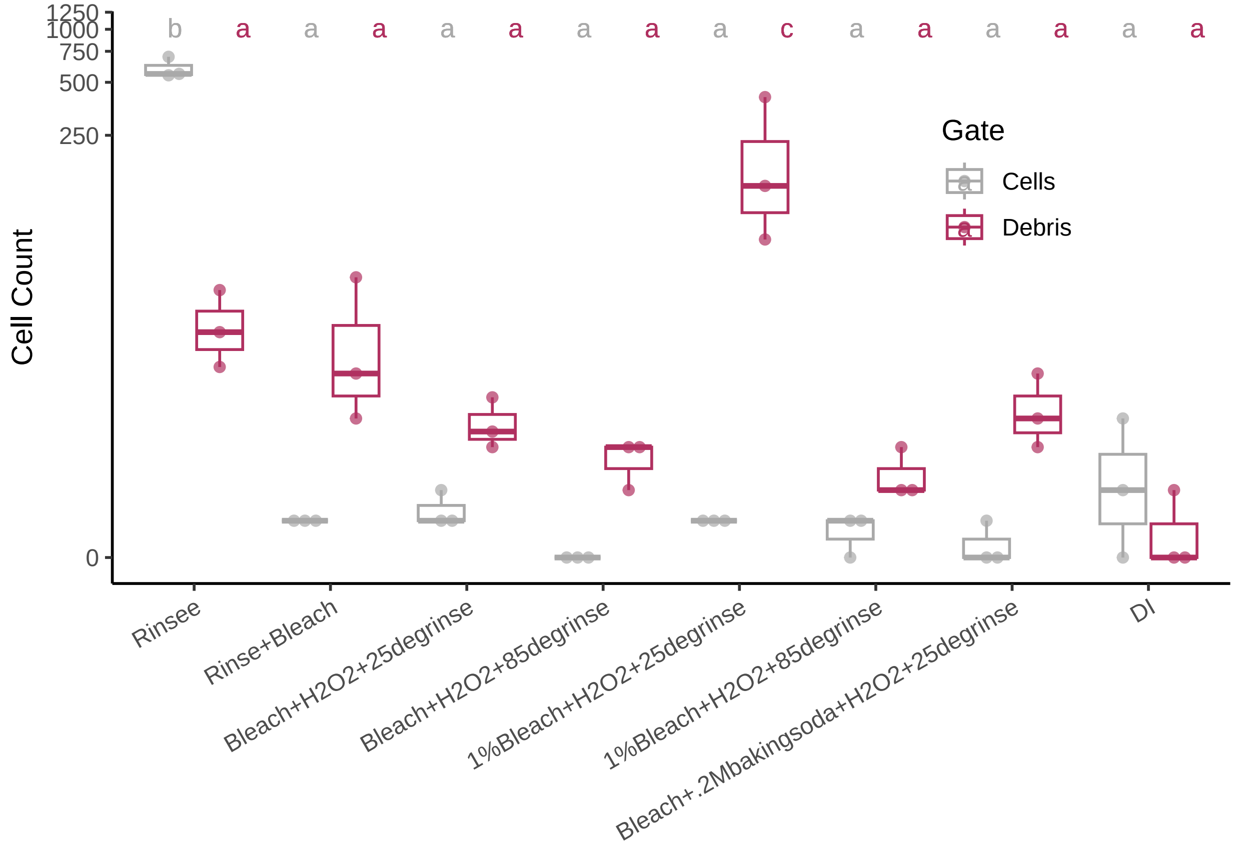


##Figure 5D

HERE I'LL look at difference between set out times

```
newdata2 <- subset(data2a, contents_b4_cleaned == "yeast" &
                   clean_method == "1%Bleach+H2O2+25degrinse")
newdata2["time_set"]<- ">7hrs"
newdata3 <- subset(data3a, contents_b4_cleaned == "yeast" &
                   clean_method == "1%Bleach+H2O2+25degrinse")
newdata3["time_set"]<- "<7hrs"
newdata4 <- subset(data4a, contents_b4_cleaned == "yeast" &
                   clean_method == "1%Bleach+H2O2+25degrinse")
newdata4["time_set"]<- ">7hrs"
neww<- rbind(newdata2,newdata3,newdata4)
```

ANOVA and Tukey's

```
model=lm(Count ~ time_set*Gate, data = neww )
ANOVA=aov(model)
summary(ANOVA)
```

```
##                Df Sum Sq Mean Sq F value Pr(>F)
## time_set        1   6806    6806   1.005 0.3282
## Gate            1  22940   22940   3.387 0.0806 .
## time_set:Gate   1   6844    6844   1.010 0.3268
## Residuals      20 135479    6774
## ---
## Signif. codes:  0 '***' 0.001 '**' 0.01 '*' 0.05 '.' 0.1 ' ' 1
```

```
TUKEY <- TukeyHSD(x=ANOVA, 'time_set:Gate', conf.level=0.95)

# Tuckey test representation :
plot(TUKEY , las=1 , col="brown")
```

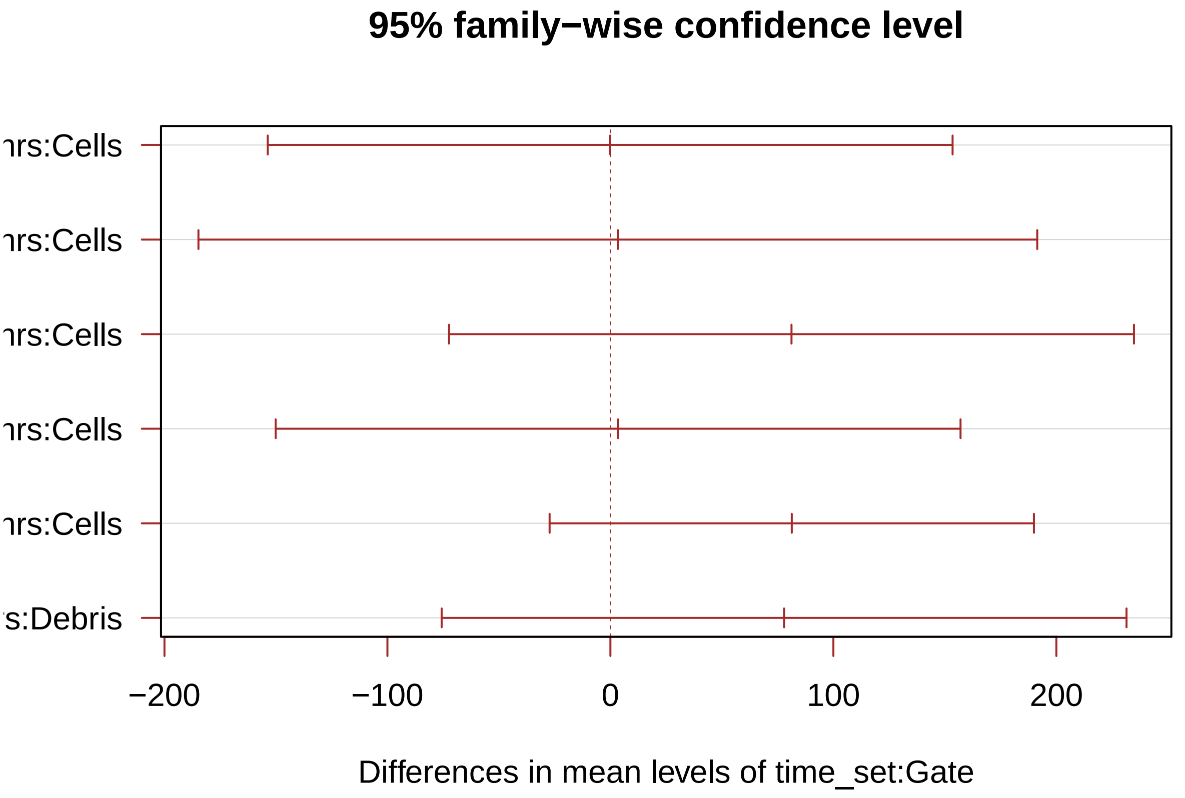


```
# I need to group the treatments that are not different each other together.
generate_label_df <- function(TUKEY, variable){

     # Extract labels and factor levels from Tukey post-hoc
     Tukey.levels <- TUKEY[[variable]][,4]
     Tukey.labels <- data.frame(multcompLetters(Tukey.levels)['Letters'])

     #I need to put the labels in the same order as in the boxplot :
     Tukey.labels$treatment=rownames(Tukey.labels)
     Tukey.labels=Tukey.labels[order(Tukey.labels$treatment) , ]
     return(Tukey.labels)
```

```r
    }

LABELS <- generate_label_df(TUKEY , "time_set:Gate")

names(LABELS)[2] ="combo"
neww["combo"] <-""
neww$combo<-paste(neww$time_set,neww$Gate,sep=":")
neww <- merge(neww, LABELS, by = "combo",
                  all.x = TRUE)


#Variation with 1% based on how long it sits.

ggplot(neww, aes(x =time_set, y = Count, color = Gate)) +
  geom_boxplot(outlier.shape = NA) +
  geom_point(alpha = 0.7, position = position_beeswarm(dodge.width = 0.75)) +
  scale_y_continuous(trans=scales::pseudo_log_trans(base = 10)) +
  labs(y = "Cell Count") +
  scale_color_manual(values = c("darkgrey", "maroon")) +
  scale_fill_manual(values = c("darkgrey", "maroon")) +
  theme_classic() +
  theme(legend.position = c(.17, .6),
        axis.text.x = element_text(angle = 30, hjust = 1, vjust = 1),
        axis.title.x = element_blank())+
  geom_text(data = neww,
            aes(x = time_set , y = 600, label = Letters),
            size = 3.4, vjust = 0, hjust = -0.5,
            position =position_dodge(width = 1) )
```

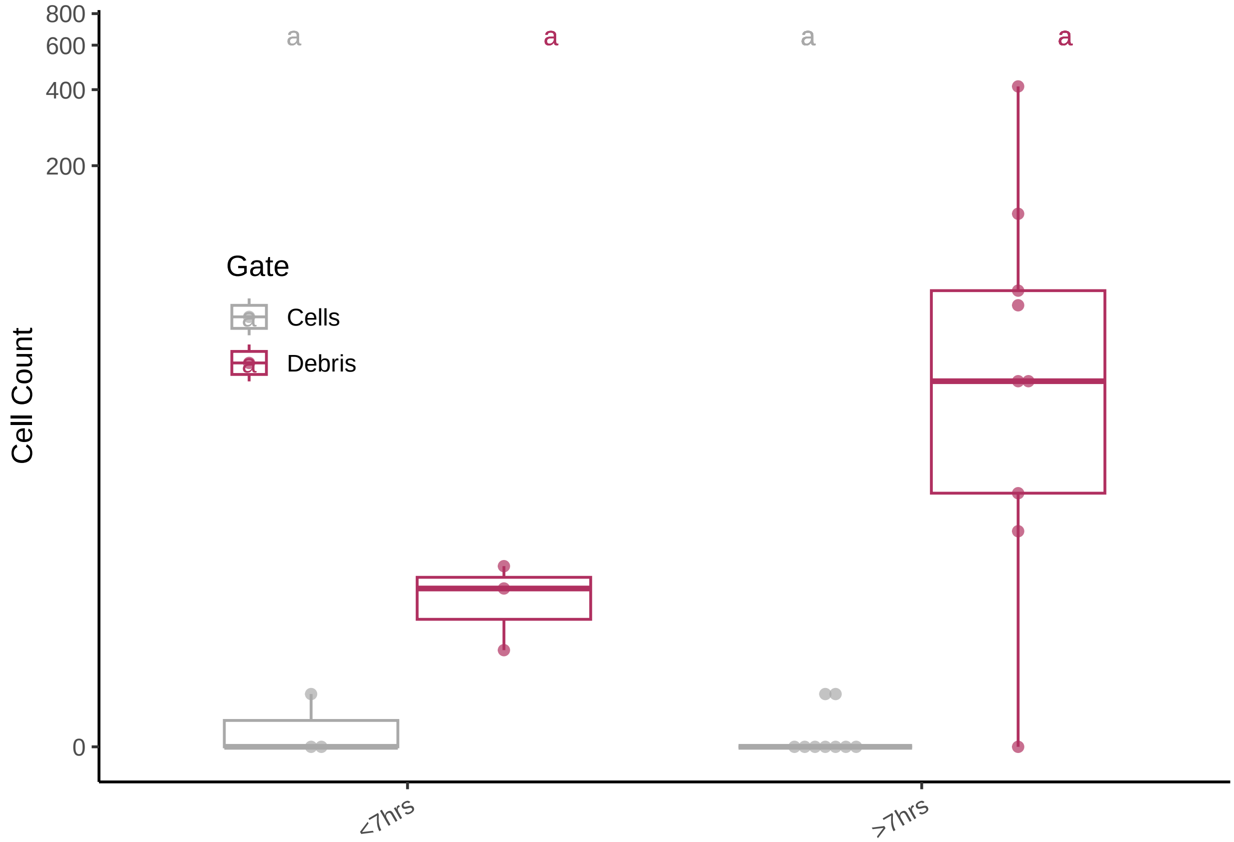


#Figure 5E

reading in and annotating data

```
#Cleaned Plate
data11 <- read.csv("rawdata/20230517_fig4E.csv")

#Fresh Plate
data22 <- read.csv("rawdata/20230423_fullclean.csv")

data11 <- data11[!grepl("All Events", data11$Gate),] #taking away an
#unnecessary datapoint
data11 <- data11[order(data11$Sample),]
data11$cleaned<-'cleaned'


data22 <- data22[!grepl("All Events", data22$Gate),]
data22 <- data22[order(data22$Sample),]
data22$cleaned<-'fresh'

newww<- rbind(data11,data22)
```

chi square for 0 cells vs. some cells.

```
#cells

#clean
ddd<- data11[,c(1,3)]
com <- data11[!grepl("Debris", data11$Gate),]
```

```r
com$clean <- "cleaned"

#fresh
ddd<- data22[,c(1,3)]
com2 <- data22[!grepl("Debris", data22$Gate),]
com2$clean <- "fresh"

rdd <- rbind(com,com2)
z<-rdd$Count > 0
rdd$events <- z
table(rdd$clean, rdd$events)
```

```
##
##           FALSE TRUE
##   cleaned    88    8
##   fresh      94    2
```

```r
chisq.test(rdd$clean, rdd$events, correct=FALSE)
```

```
##
##  Pearson's Chi-squared test
##
## data:  rdd$clean and rdd$events
## X-squared = 3.7978, df = 1, p-value = 0.05132
```

chi square for 0 debris vs. some debris.

```r
#debris

#clean
ddd<- data11[,c(1,3)]
com <- data11[!grepl("Cells", data11$Gate),]
com$clean <- "cleaned"

#fresh
ddd<- data22[,c(1,3)]
com2 <- data22[!grepl("Cells", data22$Gate),]
com2$clean <- "fresh"

rdd <- rbind(com,com2)
z<-rdd$Count > 0
rdd$events <- z
table(rdd$clean, rdd$events)
```

```
##
##           FALSE TRUE
##   cleaned    18   78
##   fresh      74   22
```

```r
chisq.test(rdd$clean, rdd$events, correct=FALSE)
```

```
##
##  Pearson's Chi-squared test
##
## data:  rdd$clean and rdd$events
## X-squared = 65.447, df = 1, p-value = 5.97e-16
```

#Figure 5F

```
newb <- subset(newww,Count> 0)

model=lm(Count ~ cleaned*Gate, data = newb, na.rm=T)
```

```
## Warning: In lm.fit(x, y, offset = offset, singular.ok = singular.ok, ...) :
##  extra argument 'na.rm' will be disregarded
```

```
ANOVA=aov(model,na.rm=T)
```

```
## Warning: In lm.fit(x, y, offset = offset, singular.ok = singular.ok, ...) :
##  extra argument 'na.rm' will be disregarded
```

```
summary(ANOVA)
```

```
##                 Df Sum Sq Mean Sq F value  Pr(>F)
## cleaned          1    348   347.6  10.990 0.00125 **
## Gate             1    304   304.4   9.624 0.00246 **
## cleaned:Gate     1     31    31.4   0.992 0.32153
## Residuals      106   3353    31.6
## ---
## Signif. codes:  0 '***' 0.001 '**' 0.01 '*' 0.05 '.' 0.1 ' ' 1
```

```
TUKEY <- TukeyHSD(x=ANOVA, 'cleaned:Gate', conf.level=0.95)

# Tuckey test representation :
plot(TUKEY , las=1 , col="brown")
```

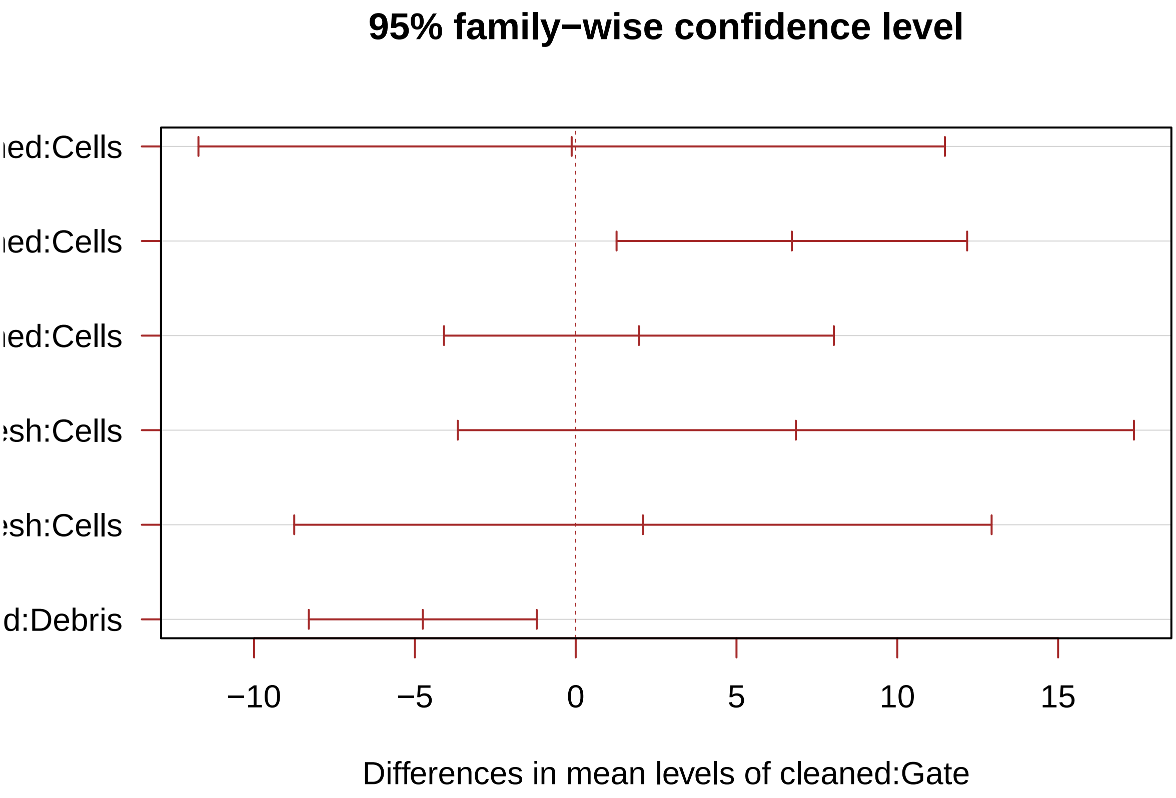


```
# I need to group the treatments that are not different each other together.
generate_label_df <- function(TUKEY, variable){
```

```r
    # Extract labels and factor levels from Tukey post-hoc
    Tukey.levels <- TUKEY[[variable]][,4]
    Tukey.labels <- data.frame(multcompLetters(Tukey.levels)['Letters'])

    #I need to put the labels in the same order as in the boxplot :
    Tukey.labels$treatment=rownames(Tukey.labels)
    Tukey.labels=Tukey.labels[order(Tukey.labels$treatment) , ]
    return(Tukey.labels)
    }

# Apply the function on my dataset
LABELS <- generate_label_df(TUKEY , "cleaned:Gate")

# Attach labels to dataframe
names(LABELS)[2] ="combo"
newb["combo"] <-""
newb$combo<-paste(newb$cleaned,newb$Gate,sep=":")
newb <- merge(newb, LABELS, by = "combo",
                  all.x = TRUE)

#plot Data with Tukey's labels
ggplot(newb, aes(x =cleaned, y = Count, color = Gate)) +
  geom_boxplot(outlier.shape = NA) +
  geom_point(alpha = 0.7, position = position_beeswarm(dodge.width = 0.75)) +
  scale_y_continuous(trans=scales::pseudo_log_trans(base = 10)) +
  labs(y = "Cell Count") +
  scale_color_manual(values = c("darkgray", "maroon")) +
  scale_fill_manual(values = c("darkgray", "maroon")) +
  theme_classic() +
  theme(
        axis.text.x = element_text(angle = 30, hjust = 1, vjust = 1),
        axis.title.x = element_blank()) +
  geom_text(data = newb,
            aes(x =cleaned , y = 50, label = Letters),
            size = 3.4, vjust = 0,
            hjust = -0.5,position =position_dodge(width = 1) )
```

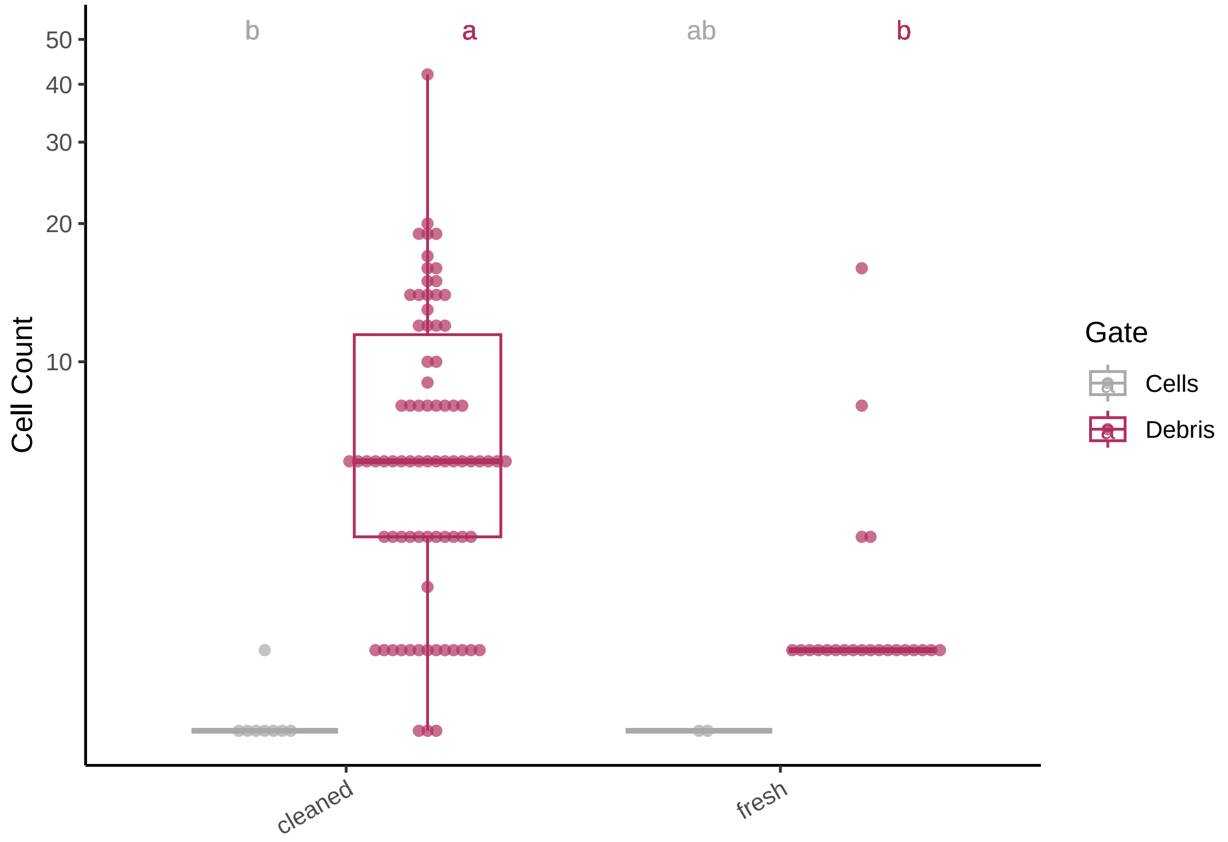


#Figure S2

```
data <- read.csv("rawdata/20230413_vmax.csv")

data_long <- tidyr::pivot_longer(
  data = data, cols = c('one','two','three','four','five','six','seven','eight','nine','ten','eleven','
```

```
simpletime <- c('maxv','r','tv','lag')

time1 <- unlist(lapply(simpletime, rep, 12))
clas<-rep(time1,6)

data_long['classification'] <- clas

A <- rep('1-2-1',3)
B<- rep('1-1-1',3)
C<- rep('1-1',3)
D<- rep('1',3)
trt <- c(A,B,C,D)
alltrts <- rep(trt,24)

one <- rep('1',48)
sevenfive <-rep('7.5',48)
allblch <- c(one,sevenfive,one,sevenfive,one,sevenfive)
data_long['treatment']<- alltrts
data_long['bleach']<-allblch
data_long$combo<-paste(data_long$treatment,data_long$bleach,sep="_")
```

Here you read in the OD600 values, along with the "Time" Column

```
data <- read.csv("rawdata/20230413_od.csv")

data_long1 <- tidyr::pivot_longer(
  data = data, cols = c('A1','A2','A3','A4','A5','A6','A7','A8','A9','A10','A11','A12','B1','B2','B3','

wells<-c('A1','A2','A3','A4','A5','A6','A7','A8','A9','A10','A11','A12','B1','B2','B3','B4','B5','B6','

A <- rep('1-2-1',3)
B<- rep('1-1-1',3)
C<- rep('1-1',3)
D<- rep('1',3)
trt <- c(A,B,C,D)
alltrts <- rep(trt,6)

alltrts<-as.data.frame((alltrts))

one <- rep('1',12)
sevenfive <-rep('7.5',12)
allblch <- c(one,sevenfive,one,sevenfive,one,sevenfive)
allblch<- as.data.frame(allblch)

alltrts$combo<-paste(alltrts[,1],allblch[,1],sep="_")
vec<-alltrts$combo

long <- rep(vec,74)

data_long1['trt']<-long
simpletime <- c(1,2,3,4,5,6,7,8,9,10,11,12,13,14,15,16,17,18,19,20,21,22,23,24,25,26,27,28,29,30,31,32,
time1 <- unlist(lapply(simpletime, rep,72))
data_long1['time'] <- time1
```

Here I'm selecting only growth data before Vmax to accurately fit an exponential growth model

```
allgrowthconstants <- c()
allxo <- c()

data_long$Column <- gsub("one", 1, data_long$Column)
data_long$Column <- gsub("two", 2, data_long$Column)
data_long$Column <- gsub("three", 3, data_long$Column)
data_long$Column <- gsub("four", 4, data_long$Column)
data_long$Column <- gsub("five", 5, data_long$Column)
data_long$Column <- gsub("six", 6, data_long$Column)
data_long$Column <- gsub("seven", 7, data_long$Column)
data_long$Column <- gsub("eight", 8, data_long$Column)
data_long$Column <- gsub("nine", 9, data_long$Column)
data_long$Column <- gsub("ten", 10, data_long$Column)
data_long$Column <- gsub("eleven", 11, data_long$Column)
data_long$Column <- gsub("twelve", 12, data_long$Column)
data_long$well <-paste(data_long$X,data_long$Column,sep="")
data_long$wellncol <-paste(data_long$well,data_long$combo,sep="+")
data_long1$trtncol <-paste(data_long1$Column,data_long1$trt,sep="+")

vec <- data_long1$trtncol
```

```
vec <- unique(vec)

for (x in vec) {
tipdat<- subset(data_long1, trtncol == x )
growdat<-subset(data_long, wellncol== x  )
growdat<-subset(growdat, classification=="tv")
time <- gsub(':','',growdat$value)
tipdat$Time<-gsub(':','',tipdat$Time)
growdat$value<-gsub(':','',growdat$value)
tipdat$use = ""

for (row in 1:nrow(tipdat)) {

  if (as.numeric(tipdat[row,"Time"]) - (as.numeric(growdat[1,"value"])) < 0){
  tipdat[row,"use"] <- "yes"}
  else {
    tipdat[row,"use"] <-"nope"
  }
}

tipdat <- subset(tipdat, tipdat$use == 'yes')

relation <- lm(log(tipdat$value)~tipdat$time)
intermediate <- coef(relation)
growthconstant <- intermediate[2]
xo <- exp(intermediate[1])
allgrowthconstants <-append(allgrowthconstants, growthconstant)
allxo <-append(allxo,xo)
}

growthefficiency <- data.frame(vec,allgrowthconstants,allxo)
growthefficiency <- growthefficiency[!grepl(0.06435705, growthefficiency$allxo),]
```

```
ge2<-growthefficiency
ge2[c('well', 'treatment')] <- str_split_fixed(ge2$vec, '\\+', 2)
ge2[c('trtmnt', 'bleach')] <- str_split_fixed(ge2$treatment, '_', 2)
#ge <- subset(ge, select = -c(vec,treatment))
ge2 <- subset(ge2, select = -c(vec,well,treatment))
```

```
ge2$trtmnt <- sub('-','_',ge2$trtmnt)
ge2$trtmnt <- sub('-','_',ge2$trtmnt)

model=lm(allxo ~ trtmnt*bleach, data = ge2 )

ANOVA=aov(model)

TUKEY <- TukeyHSD(x=ANOVA, 'trtmnt:bleach', conf.level=0.95)
plot(TUKEY , las=1 , col="brown")
```

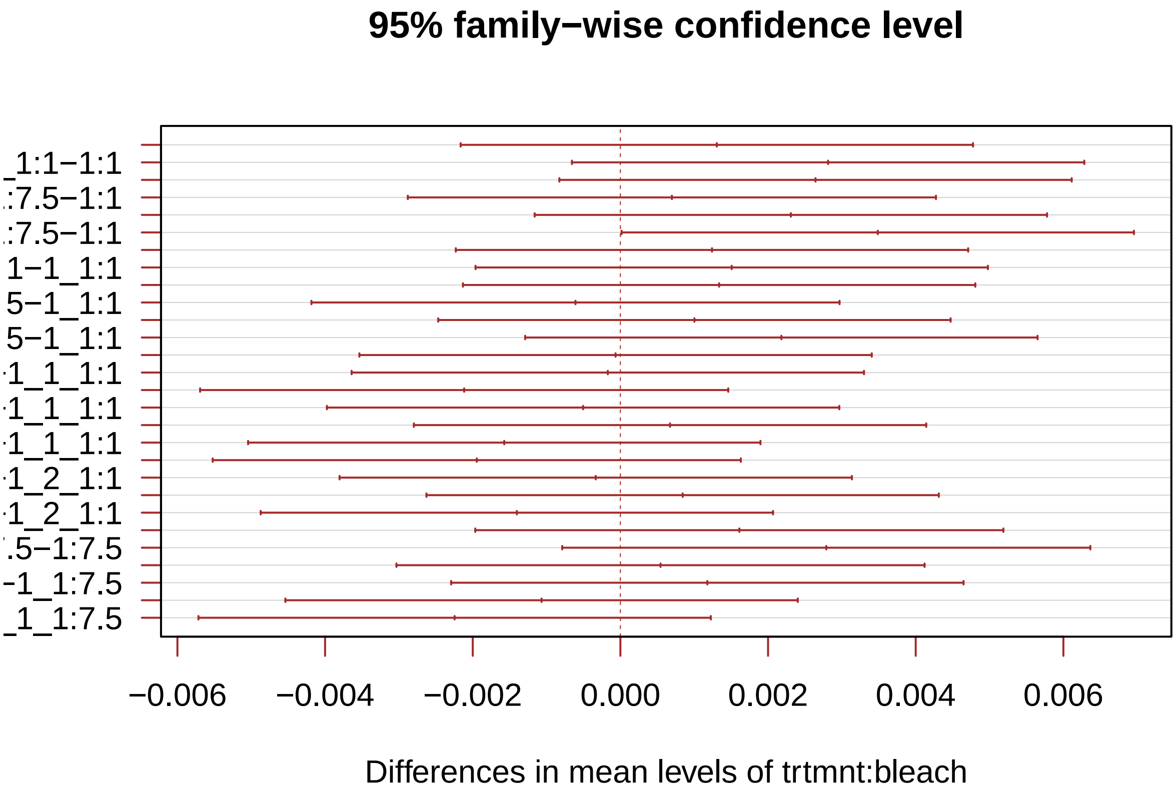


```
generate_label_df <- function(TUKEY, variable){

     # Extract labels and factor levels from Tukey post-hoc
     Tukey.levels <- TUKEY[[variable]][,4]
     Tukey.labels <- data.frame(multcompLetters(Tukey.levels)['Letters'])

     #I need to put the labels in the same order as in the boxplot :
     Tukey.labels$treatment=rownames(Tukey.labels)
     Tukey.labels=Tukey.labels[order(Tukey.labels$treatment) , ]
     return(Tukey.labels)
     }

LABELS <- generate_label_df(TUKEY , "trtmnt:bleach")

names(LABELS)[2] ="combo"



ge2["combo"] <-""
ge2$combo<-paste(ge2$trtmnt,ge2$bleach,sep=":")
ge2 <- merge(ge2, LABELS, by = "combo",
                  all.x = TRUE)
LABELS[c('trtmnt', 'bleach')] <- str_split_fixed(LABELS$combo, ':', 2)
```

```
labelconv<- c("1" = "No Rinses","1_1"= "1 water rinse","1_1_1"="1 water rinse +\n1 H2O2 rinse","1_2_1" =

ggplot(ge2, aes(x =trtmnt, y = allxo, fill=bleach)) +
  geom_boxplot(outlier.shape = NA) +
  geom_point(alpha = 0.7, position = position_beeswarm(dodge.width = 0.75)) +
```

```
  scale_y_continuous(trans=scales::pseudo_log_trans(base = 10)) +
  labs(y = "Xo") +
  scale_color_manual(values = c("darkgrey", "maroon")) +
  scale_fill_manual(values = c("darkgrey", "maroon")) +
  theme_classic() +
  theme(legend.position = c(.96,.15),
        axis.text.x = element_text(angle = 30, hjust = 1, vjust = 1)) +geom_text(data = LABELS, aes(x =
```

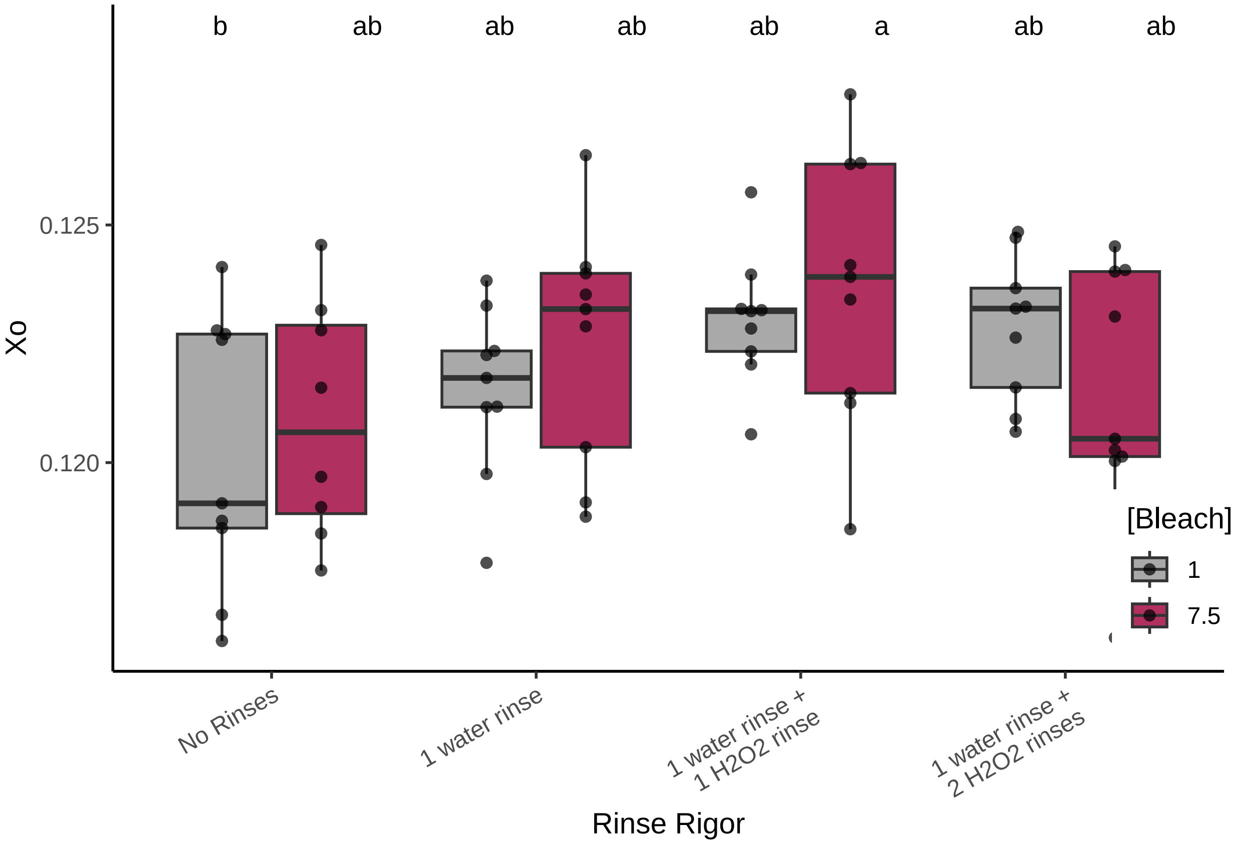


#Figure S4

ANOVA and Tukey's Using the untrimmed dataset from Figure 5

```
neww<-saveneww
model=lm(Count ~ clean_method*Gate, data = neww )
ANOVA=aov(model)
summary(ANOVA)
```

```
##                    Df Sum Sq Mean Sq F value   Pr(>F)
## clean_method        7 240312   34330   5.483 1.74e-05 ***
## Gate                1  12009   12009   1.918    0.169
## clean_method:Gate   7 263760   37680   6.018 5.04e-06 ***
## Residuals         120 751381    6262
## ---
## Signif. codes:  0 '***' 0.001 '**' 0.01 '*' 0.05 '.' 0.1 ' ' 1
```

```
TUKEY <- TukeyHSD(x=ANOVA, 'clean_method:Gate', conf.level=0.95)

# Tuckey test representation :
```

```r
plot(TUKEY , las=1 , col="brown")
```

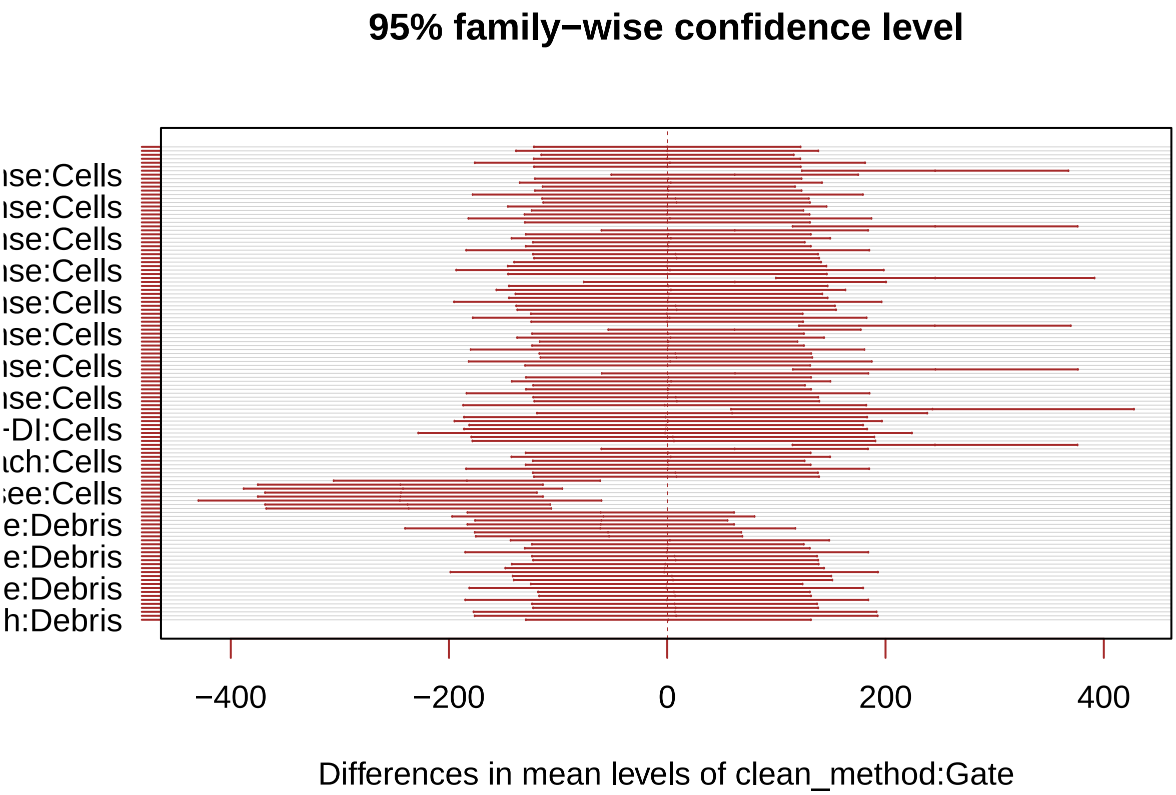


```r
generate_label_df <- function(TUKEY, variable){

     # Extract labels and factor levels from Tukey post-hoc
     Tukey.levels <- TUKEY[[variable]][,4]
     Tukey.labels <- data.frame(multcompLetters(Tukey.levels)['Letters'])

     #I need to put the labels in the same order as in the boxplot :
     Tukey.labels$treatment=rownames(Tukey.labels)
     Tukey.labels=Tukey.labels[order(Tukey.labels$treatment) , ]
     return(Tukey.labels)
     }

# Apply the function on my dataset
#LABELS <- generate_label_df(TUKEY , "steady_states$strain_and_trt")
LABELS <- generate_label_df(TUKEY , "clean_method:Gate")

names(LABELS)[2] ="combo"

neww["combo"] <-""
neww$combo<-paste(neww$clean_method,neww$Gate,sep=":")
neww<- merge(neww, LABELS, by = "combo",
                  all.x = TRUE)

ggplot(neww, aes(x =fct_relevel(clean_method,"20X yeast","Bleached yeast", "Rinsee","Rinse+Bleach","Blea
               y = Count, color = Gate)) +
  geom_boxplot(outlier.shape = NA) +
```

```
  geom_point(alpha = 0.7, position = position_beeswarm(dodge.width = 0.75)) +
  scale_y_continuous(trans=scales::pseudo_log_trans(base = 10)) +
  labs(y = "Cell Count") +
  scale_color_manual(values = c("darkgrey", "maroon")) +
  scale_fill_manual(values = c("darkgrey", "maroon")) +
  theme_classic() +
  theme(legend.position = c(.8, .7),
        axis.text.x = element_text(angle = 30, hjust = 1, vjust = 1),
        axis.title.x = element_blank())+
  geom_text(data = neww, aes(x = clean_method , y = 900, label = Letters),
            size = 3.4, vjust = 0, hjust = -0.5,
            position =position_dodge(width = 1) )+
  scale_x_discrete(labels=labelconv)
```

```
## Warning: 2 unknown levels in `f`: 20X yeast and Bleached yeast
## 2 unknown levels in `f`: 20X yeast and Bleached yeast
```

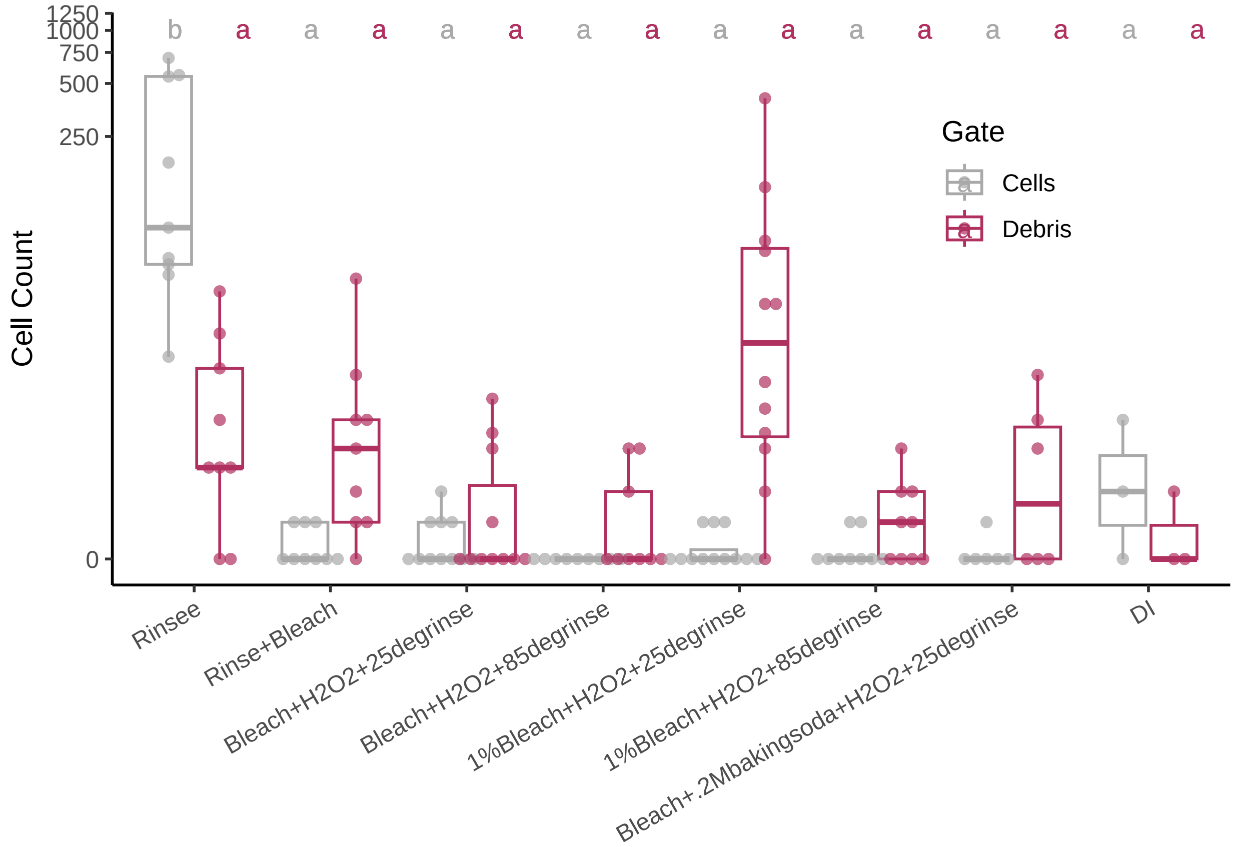


#Figure S5 subsetting and grouping the Ultrapure water samples

```
newdata1 <- subset(data1a,contents_b4_cleaned == "DI" |
                     contents_b4_cleaned == "none")
newdata2 <- subset(data2a, contents_b4_cleaned == "DI" )
#newdata2["time_set"]<- ">7hrs"
newdata3 <- subset(data3a, contents_b4_cleaned == "DI")
#newdata3["time_set"]<- "<7hrs"
newdata4 <- subset(data4a, contents_b4_cleaned == "DI")
#newdata4["time_set"]<- ">7hrs"
```

```r
neww<- rbind(newdata1,newdata2,newdata3,newdata4)
```

ANOVA and Tukey's Using the untrimmed dataset from Figure 5

```r
model=lm(Count ~ clean_method*Gate, data = neww )
ANOVA=aov(model)
summary(ANOVA)
```

```
##                    Df Sum Sq Mean Sq F value   Pr(>F)    
## clean_method        7   95.6   13.65   2.331   0.0288 *  
## Gate                1  100.9  100.90  17.226 6.18e-05 ***
## clean_method:Gate   7  107.3   15.33   2.618   0.0149 *  
## Residuals         122  714.6    5.86                     
## ---
## Signif. codes:  0 '***' 0.001 '**' 0.01 '*' 0.05 '.' 0.1 ' ' 1
```

```r
TUKEY <- TukeyHSD(x=ANOVA, 'clean_method:Gate', conf.level=0.95)

# Tuckey test representation :
plot(TUKEY , las=1 , col="brown")
```

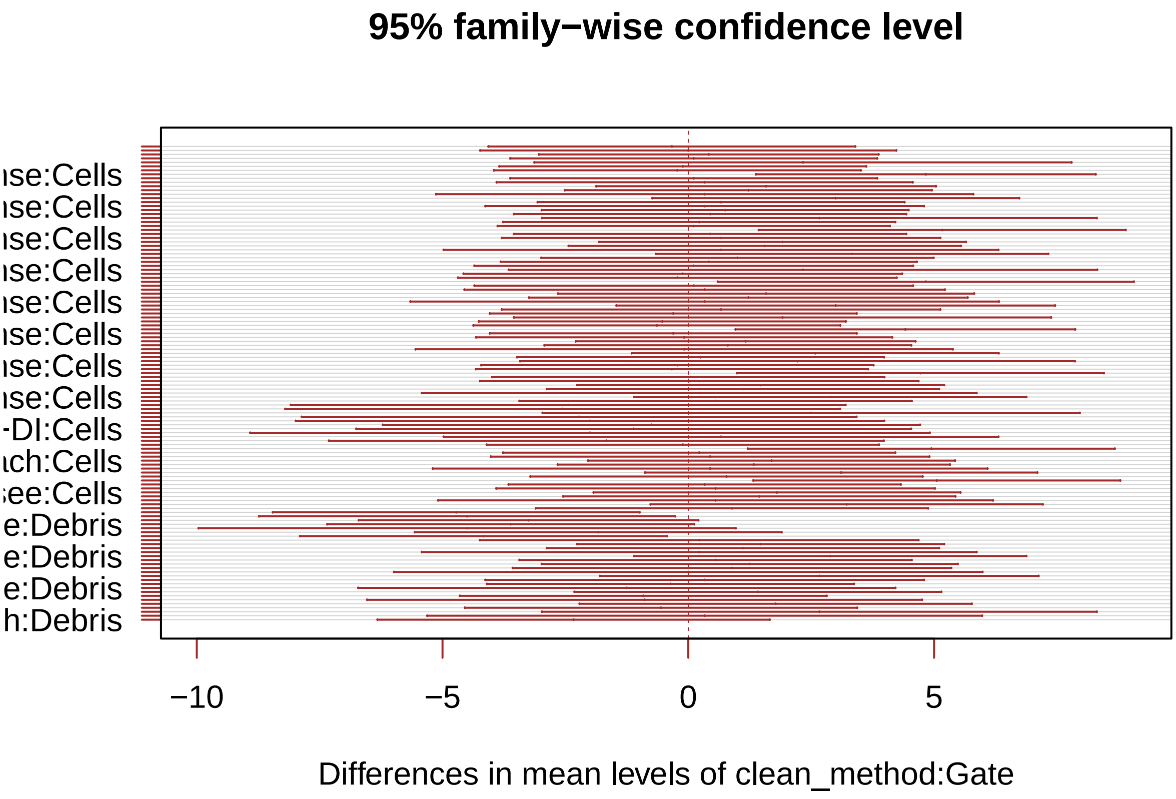


```r
generate_label_df <- function(TUKEY, variable){

     # Extract labels and factor levels from Tukey post-hoc
     Tukey.levels <- TUKEY[[variable]][,4]
     Tukey.labels <- data.frame(multcompLetters(Tukey.levels)['Letters'])

     #I need to put the labels in the same order as in the boxplot :
     Tukey.labels$treatment=rownames(Tukey.labels)
```

```
     Tukey.labels=Tukey.labels[order(Tukey.labels$treatment) , ]
     return(Tukey.labels)
     }

# Apply the function on my dataset
#LABELS <- generate_label_df(TUKEY , "steady_states$strain_and_trt")
LABELS <- generate_label_df(TUKEY , "clean_method:Gate")

names(LABELS)[2] ="combo"

neww["combo"] <-""
neww$combo<-paste(neww$clean_method,neww$Gate,sep=":")
neww<- merge(neww, LABELS, by = "combo",
                  all.x = TRUE)

ggplot(neww, aes(x =fct_relevel(clean_method,"20X yeast","Bleached yeast", "Rinsee","Rinse+Bleach","Blea
               y = Count, color = Gate)) +
  geom_boxplot(outlier.shape = NA) +
  geom_point(alpha = 0.7, position = position_beeswarm(dodge.width = 0.75)) +
  scale_y_continuous(trans=scales::pseudo_log_trans(base = 10)) +
  labs(y = "Cell Count") +
  scale_color_manual(values = c("darkgrey", "maroon")) +
  scale_fill_manual(values = c("darkgrey", "maroon")) +
  theme_classic() +
  theme(legend.position = c(.8, .7),
        axis.text.x = element_text(angle = 30, hjust = 1, vjust = 1),
        axis.title.x = element_blank())+
  geom_text(data = neww, aes(x = clean_method , y = 900, label = Letters),
            size = 3.4, vjust = 0, hjust = -0.5,
            position =position_dodge(width = 1) )+
  scale_x_discrete(labels=labelconv)
```

```
## Warning: 2 unknown levels in `f`: 20X yeast and Bleached yeast
## 2 unknown levels in `f`: 20X yeast and Bleached yeast
```

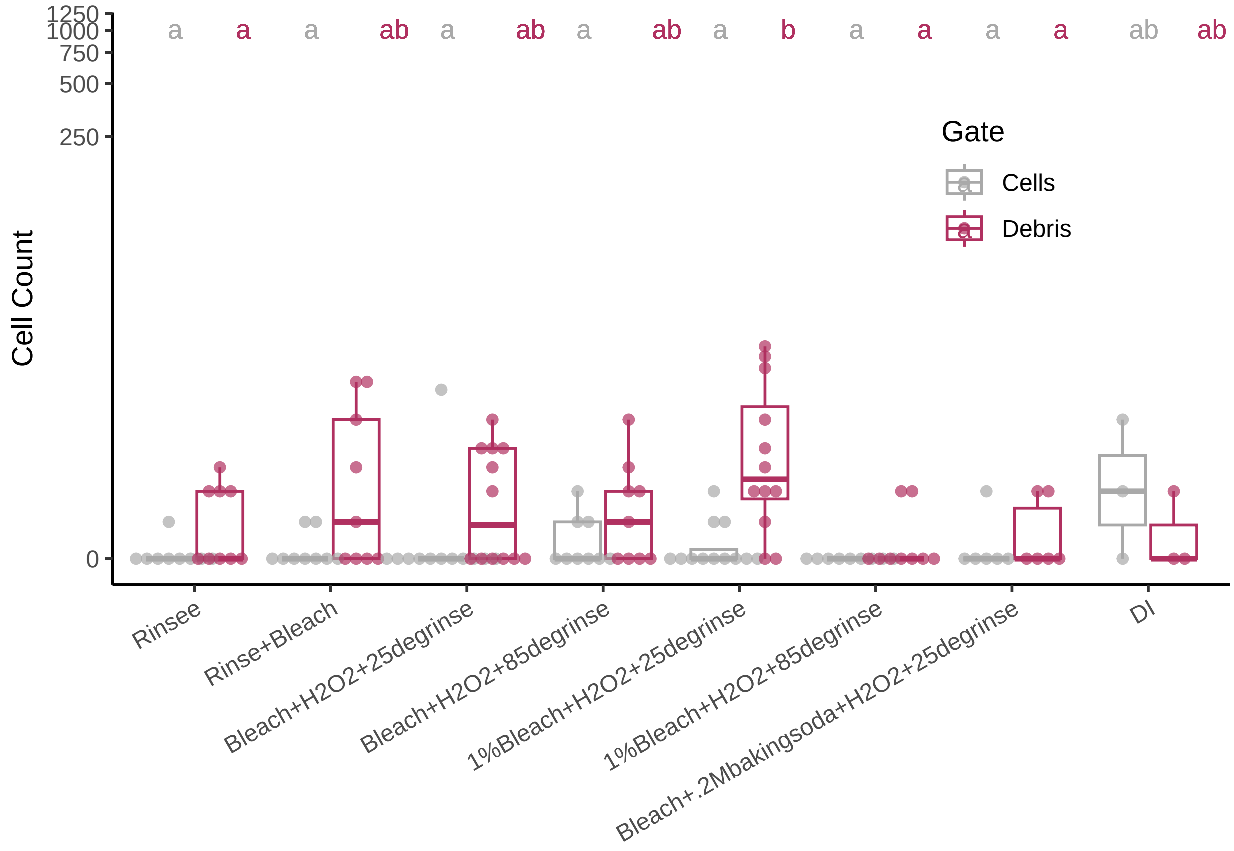


#Figure S6 reading in and annotating data

```
data1 <- read.csv("rawdata/20230423_singleintegraged.csv")
data2 <- read.csv("rawdata/20230423_fullclean.csv")

data1 <- data1[!grepl("All Events", data1$Gate),] #taking away an unnecessary datapoint
data1 <- data1[order(data1$Sample),]
data1$cleaned<-'cleaned'

row <- c("row 1","row 2","row 3","row 4","row 5","row 6","row 7","row 8")
row <- unlist(lapply(row, rep,24))

data1['row'] <- row
data1$grp <- paste(data1[,5],data1[,6])

data2 <- data2[!grepl("All Events", data2$Gate),]
data2 <- data2[order(data2$Sample),]
data2$cleaned<-'fresh'
data2["row"]<-row
data2$grp <- paste(data2[,5],data2[,6])
neww2<- rbind(data1,data2)
```

ANOVA and Tukey's

```
model=lm(Count ~ grp*Gate, data = neww2, na.rm=T)
```

```
## Warning: In lm.fit(x, y, offset = offset, singular.ok = singular.ok, ...) :
##  extra argument 'na.rm' will be disregarded
```

```r
ANOVA=aov(model,na.rm=T)
```

```
## Warning: In lm.fit(x, y, offset = offset, singular.ok = singular.ok, ...) :
##  extra argument 'na.rm' will be disregarded
```

```r
summary(ANOVA)
```

```
##              Df Sum Sq Mean Sq F value   Pr(>F)
## grp          15 123661    8244   13.84  < 2e-16 ***
## Gate          1  35209   35209   59.09 1.51e-13 ***
## grp:Gate     15 122617    8174   13.72  < 2e-16 ***
## Residuals   352 209726     596
## ---
## Signif. codes:  0 '***' 0.001 '**' 0.01 '*' 0.05 '.' 0.1 ' ' 1
```

```r
TUKEY <- TukeyHSD(x=ANOVA, 'grp:Gate', conf.level=0.95)
plot(TUKEY , las=1 , col="brown")
```

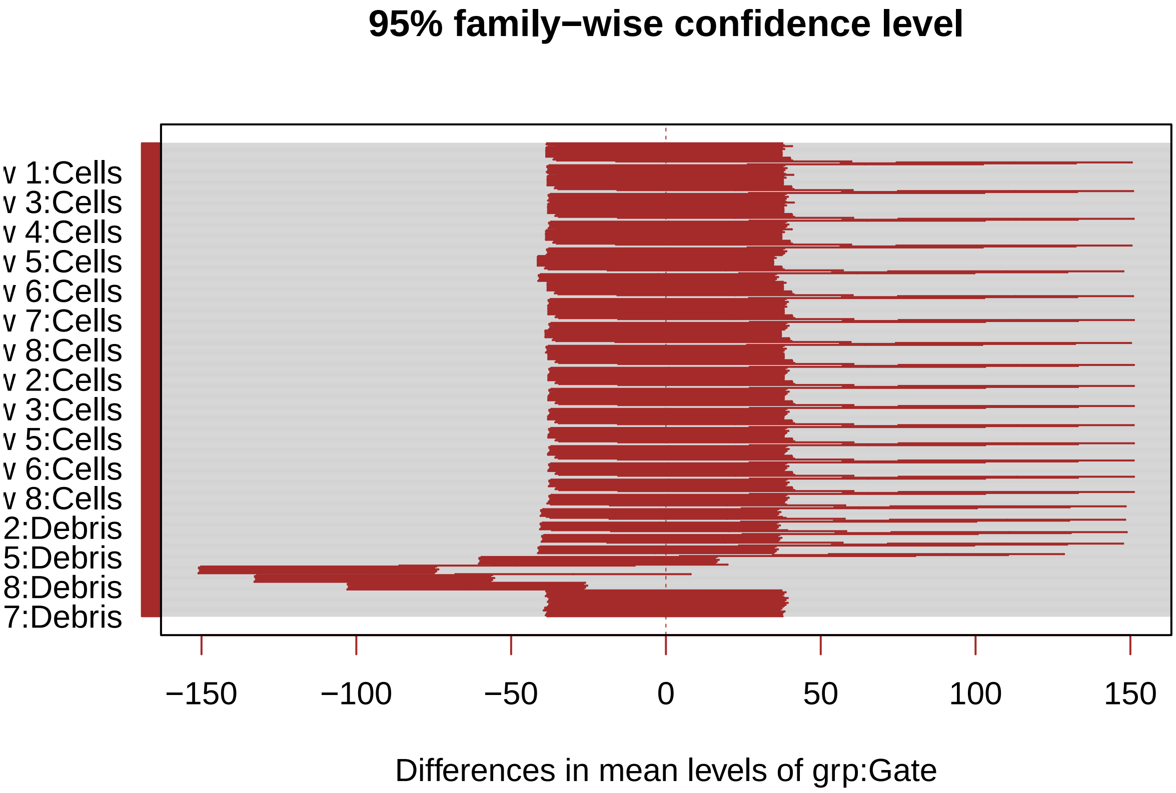


```r
generate_label_df <- function(TUKEY, variable){

     # Extract labels and factor levels from Tukey post-hoc
     Tukey.levels <- TUKEY[[variable]][,4]
     Tukey.labels <- data.frame(multcompLetters(Tukey.levels)['Letters'])

     #I need to put the labels in the same order as in the boxplot :
     Tukey.labels$treatment=rownames(Tukey.labels)
     Tukey.labels=Tukey.labels[order(Tukey.labels$treatment) , ]
     return(Tukey.labels)
     }
```

```
# Apply the function on my dataset
#LABELS <- generate_label_df(TUKEY , "steady_states$strain_and_trt")
LABELS <- generate_label_df(TUKEY , "grp:Gate")

names(LABELS)[2] ="combo"

neww2["combo"] <-""
neww2$combo<-paste(neww2$grp,neww2$Gate,sep=":")
neww2 <- merge(neww2, LABELS, by = "combo",
                 all.x = TRUE)
```

```
labelconv<- c("20X yeast"="5% of initial\nyeast count", "Rinsee" = "Two Rinses","Rinse+Bleach"= "Rinse +


ggplot(neww2, aes(x =fct_relevel(grp,"fresh row 1", "cleaned row 1","fresh row 2", "cleaned row 2","fres
  geom_boxplot(outlier.shape = NA) +
  geom_point(alpha = 0.7, position = position_beeswarm(dodge.width = 0.75)) +
  scale_y_continuous(trans=scales::pseudo_log_trans(base = 10)) +
  labs(y = "Coefficient of Variation") +
  scale_color_manual(values = c("darkgray", "maroon")) +
  scale_fill_manual(values = c("darkgray", "maroon")) +
  theme_classic() +
  theme(legend.position = c(.2, .6),
        axis.text.x = element_text(angle = 30, hjust = 1, vjust = 1),
        axis.title.x = element_blank()) +geom_text(data = neww2, aes(x =grp , y = 900, label = Letters)
```

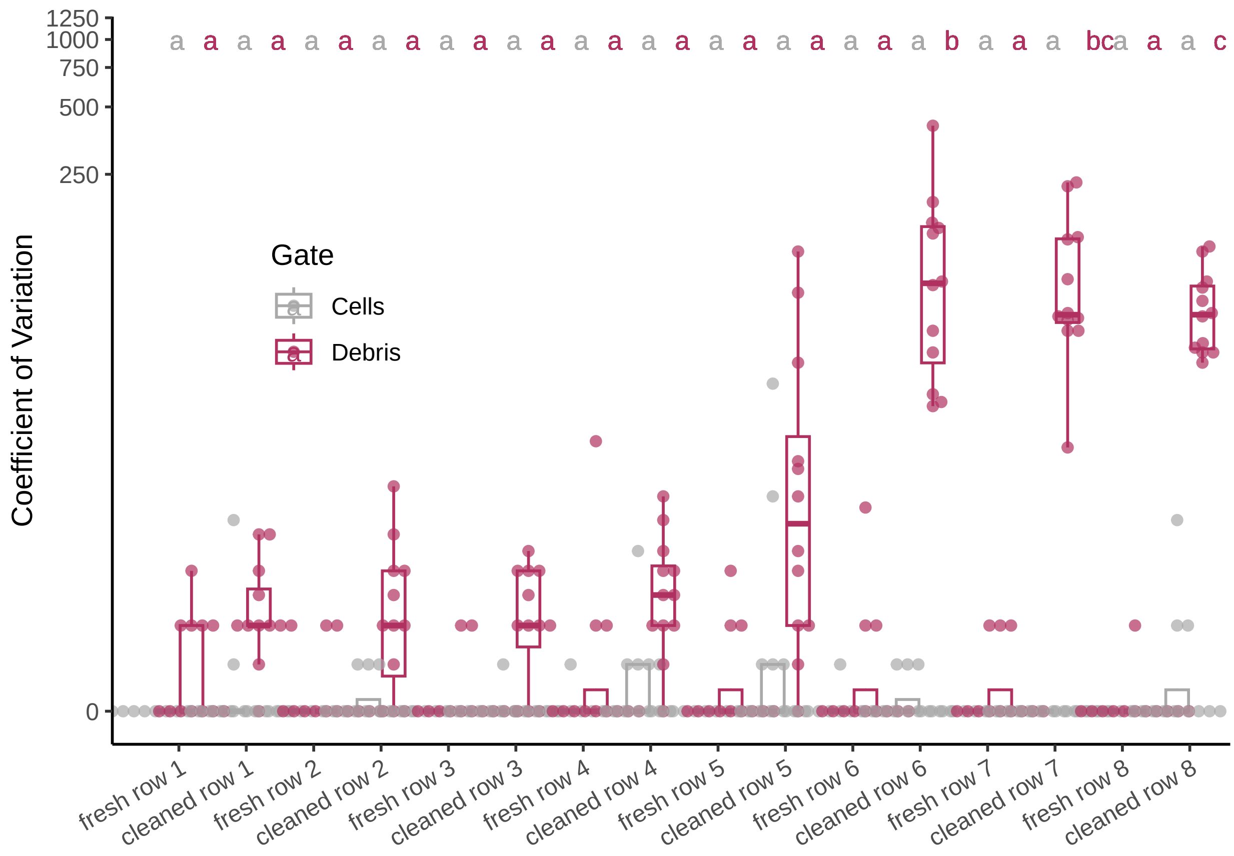
